# Predicting drought-induced tree mortality in Swiss beech forests hinges upon predisposing and inciting factors

**DOI:** 10.64898/2025.12.15.694299

**Authors:** G. Marano, U. Hiltner, K. Meusburger, T. Hands, H. Bugmann

**Affiliations:** Forest Ecology, Institute of Terrestrial Ecosystems, Department of Environmental Systems Science, ETH Zurich, Switzerland; Swiss Federal Institute for Forest, Snow and Landscape Research (WSL), Birmensdorf, Switzerland

**Keywords:** Tree mortality, Beech decline, Soil water deficit, ForClim, Dynamic Vegetation Model

## Abstract

The increase in the frequency and severity of drought-induced tree mortality in many European low-elevation forests poses considerable challenges to forest management and requires an understanding of its causes. We propose a novel framework for integrating the factors underlying drought-induced tree mortality in a dynamic vegetation model.

We evaluate whether this framework accurately reproduces drought-related mortality in six mesic beech-dominated stands in 2018-2020, and over multiple years in a xeric Scots pine-dominated stand in Switzerland. Additionally, we investigate its behavior along a large climatic gradient in central Europe. We employ a three-step approach.

First, we evaluate multiple drought indices for capturing tree growth responses to extreme drought. We find that in contrast to widespread indices such as SPI and SPEI, the ForClim drought index captures growth responses to drought intensity during summer, the growing period, and annually.

Second, we assess in detail the capability of the ForClim soil moisture model to simulate soil water dynamics, comparing it to the mechanistic soil-vegetation-atmosphere model LWFBrook90. The ForClim soil moisture model adequately simulates soil water dynamics, particularly in extreme drought years.

Third, based on Manion’s Decline Disease Theory, we develop a novel mortality sub-model that combines predisposing and inciting factors. Its integration in ForClim captures drought-induced mortality events in the mesic beech forests as well as the multi-year mortality at the xeric Scots pine-dominated stand. Along a climatic gradient in central Europe, the model provides good quantifications of Potential Natural Vegetation.

The novel framework to capture drought-related tree mortality is simple yet produces accurate results. The underlying hypothesis regarding the factors leading to drought-induced tree mortality is promising but requires further tests for its generality.

## 1 Introduction

Over the last decades, climate-driven disturbances have progressively exposed temperate forests to stress, often compromising ecosystem functioning and the provision of ecosystem services (Breshears et al., 2005; Gilliam, 2016; Senf & Seidl, 2021). An increase in the frequency, intensity, and duration of droughts (Buras et al., 2019; Jia et al., 2019; Rakovec et al., 2022) has already led to substantial changes of forest structure in Europe (Schuldt et al. 2020) and worldwide (Allen et al. 2015, Millar et al. 2015). Droughts, which span from weekly to seasonal to multi-decadal, are important drivers of forest dynamics (Cook et al., 2015), triggering declines in productivity and eventually increasing the likelihood of mortality of both deciduous and conifer trees (Allen et al. 2010, Anderegg et al., 2013).

While spruce (*Picea abies*) dieback has been observed for a long time in low-elevation forests in Europe (Klimo et al., 2000; Bosela et al. 2021), the extreme droughts since 2015 have caused unprecedented mortality events also in beech (*Fagus sylvatica*) forests, including areas previously not considered to be drought-prone (Schuldt et al., 2020; Frei et al., 2022; Neycken et al., 2022). It is not clear whether this drought-induced tree mortality, often preceded by canopy decline, is the result of prolonged or rapid stress, or both (Herguido et al., 2016; Neycken et al., 2022; Klesse et al., 2022; Schmied et al., 2023), and to what extent it is the result of the interplay of abiotic and biotic stressors (Manion, 1981).

Declines in tree productivity are commonly assessed based on tree-ring data and drought indices (Bigler et al. 2006, Bose et al. 2021). However, the most frequently employed drought indices neglect the role of soil water, although it is a major driver of productivity decline, at least in more recent drought events (Liu et al., 2020, Carminati & Javaux, 2020). To predict forest responses to climate change, process-based models are needed because they allow to assess the interactions among the multiple factors that shape forest dynamics (Manusch et al., 2014; Huber et al., 2020; Bugmann & Seidl, 2022). Yet, to date dynamic vegetation models have failed to reproduce drought-induced tree mortality (McDowell et al., 2013; Steinkamp & Hickler, 2015; Hendrik & Cailleret, 2017; De Kauwe et al., 2022). This may be due to i) non-linear interactions among processes (Wang et al., 2012; Xu et al. 2013; Camarero, 2021), ii) high process complexity that is difficult to capture (Orth et al., 2015; Lin & Yang, 2022), and iii) scarcity of long-term, stand-scale observations with at least an annual resolution (Parolari et al. 2014; Hartmann et al., 2018; Camarero, 2021).

Drought-induced tree mortality is generally thought to be due to the synergistic action of multiple factors rather than isolated causes (McDowell et al., 2008-2011; Adams et al., 2017; Gazol & Camarero, 2022). Three common hypotheses in this context are i) hydraulic failure (Martínez-Vilalta et al., 2002; Brodribb & Cochard, 2008), ii) carbon starvation (Sevanto et al., 2014), and iii) biotic agents (Neely & Manion, 1991). The interplay among these and likely further factors may explain drought-induced tree mortality (“Decline Disease” framework, Manion, 1981), including non-linear interactions among multiple stressors and specifically distinguishing predisposing, inciting, and contributing factors (hereafter termed the “predisposing and inciting factor” (PIC) scheme). The PIC scheme posits that predisposing factors (chronic stressors) enhance the vulnerability of trees to inciting (short-term acute stress) and contributing factors (secondary, long-term stress agents), eventually resulting in tree death (Manion, 1981; Pedersen, 1998; Wang et al., 2012). However, in dynamic vegetation models, the focus has predominantly been on either predisposing factors (e.g., low growth rate, growth efficiency) or inciting factors (e.g., hydraulic failure), rather than their combination (Wang et al., 2012). A few models that incorporated both processes (TRIPLEX by Liu et al. 2021, ED(X) by Moorcroft et al. 2001, GOTILWA+ by Nadal-Sala et al. 2017) still were limited in their ability to capture observed tree mortality patterns. In these studies, indicators of hydraulic failure and carbon starvation indicators were derived from the same data to both calibrate the mortality formulation and determine its causes—without independent validation or incorporation of temporal dynamics—thus potentially involving circular reasoning.

In this study, we developed a model based on the PIC scheme and tested whether it captures drought-induced mortality. We then compared its performance to an earlier model version that does not distinguish between predisposing and inciting factors. Specifically, we addressed the following questions:

1. *Which drought index is best suited for capturing tree growth response to drought under non-extreme climatic conditions?* We posit that drought indices lacking information on soil water content provide inferior performance, regardless of their temporal resolution.
2. *What complexity is required to simulate the soil water balance in extremely dry years accurately?* We hypothesize that a low-complexity approach, e.g. a single-layer soil bucket model at monthly temporal resolution, provides a good compromise between ease of parameterization and process representation.
3. *Do trees succumb to reduced growth alone, or is it due to compound events such as prolonged stress followed by short-intense events?* According to the PIC scheme, trees exposed to prolonged stress are most susceptible to mortality when a short-term, intense event such as a seasonal drought occurs. We suggest that the PIC scheme helps to elucidate the drought-induced mortality observed at many low-elevation sites in Europe during 2018-2020.

## 2. Materials and Methods

### 2.1. Site selection

We selected six even-aged mature beech-dominated forests in Switzerland that had experienced varying degrees of canopy decline and tree mortality in 2018-2020 (Neycken et al., 2022) (*Figure 1*, *Table 1*). They are located on the Swiss Plateau (Blattenberg, Grosszinggibrunn, Rossberg, Tüeliboden, Vogtacher) and in the Jura mountains (Usserholz), featuring highly varying soil properties as evident from the Available Water holding Capacity (hereafter AWC, 86 to 155 mm), and annual precipitation sum (914 to 1325 mm). Despite the observed mortality events, these sites are characteristic of the mesic climate of central Europe, where drought is usually just mellow and drought-related mortality is quite rare (Supplementary Material, Figure SM A1).

**Figure 1:**
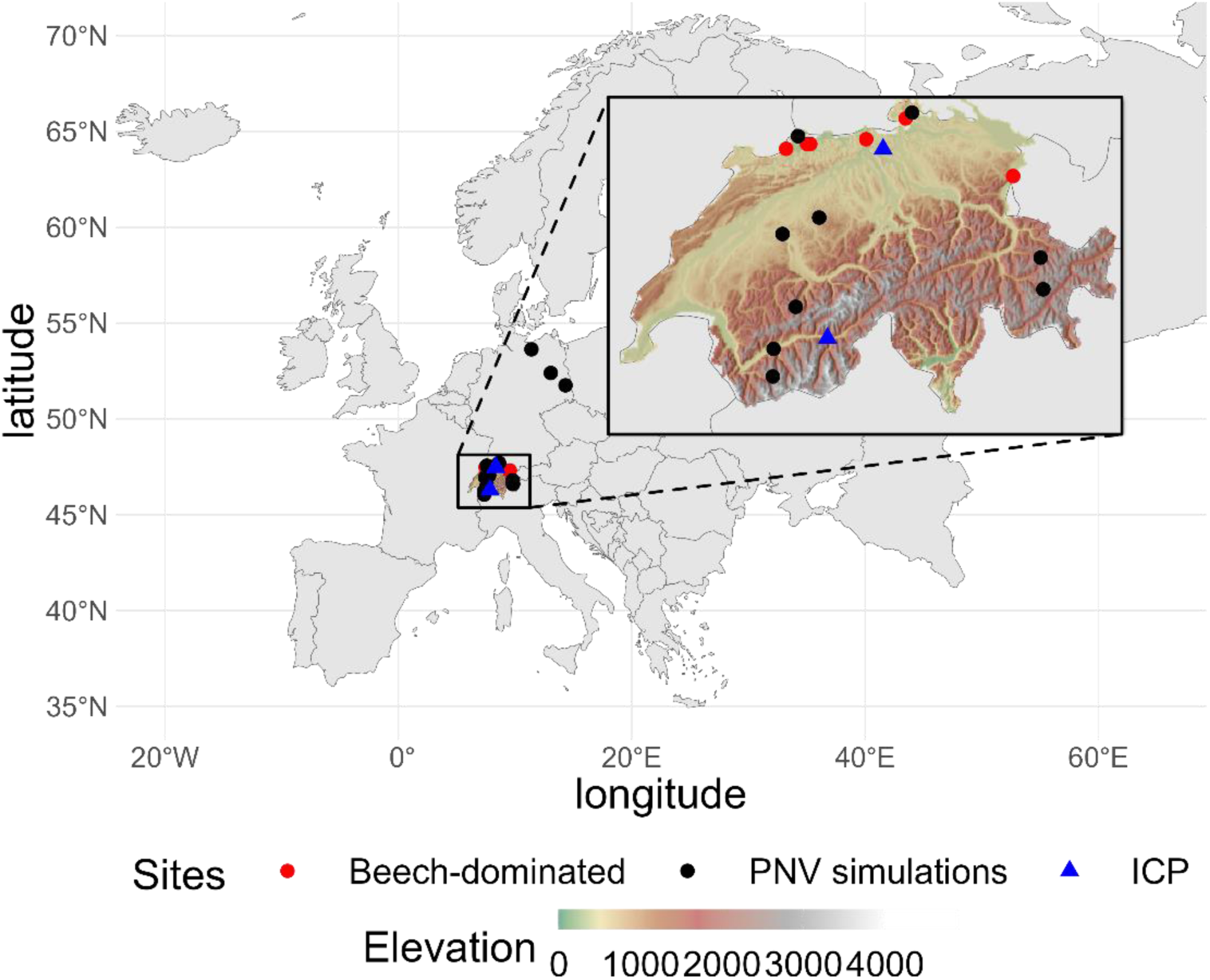
Study sites classified as main sites (red dots) where drought-induced mortality in 2018-2020 was observed, the ICP-Level II plots (abbr. ICP, blue triangles) and sites where Potential Natural Vegetation was simulated along an elevational and climatic gradient (*Bugmann & Solomon, 2000*) in Switzerland and Germany (black dots). Elevational map of Switzerland from Swisstopo (2014).

**Table 1.**
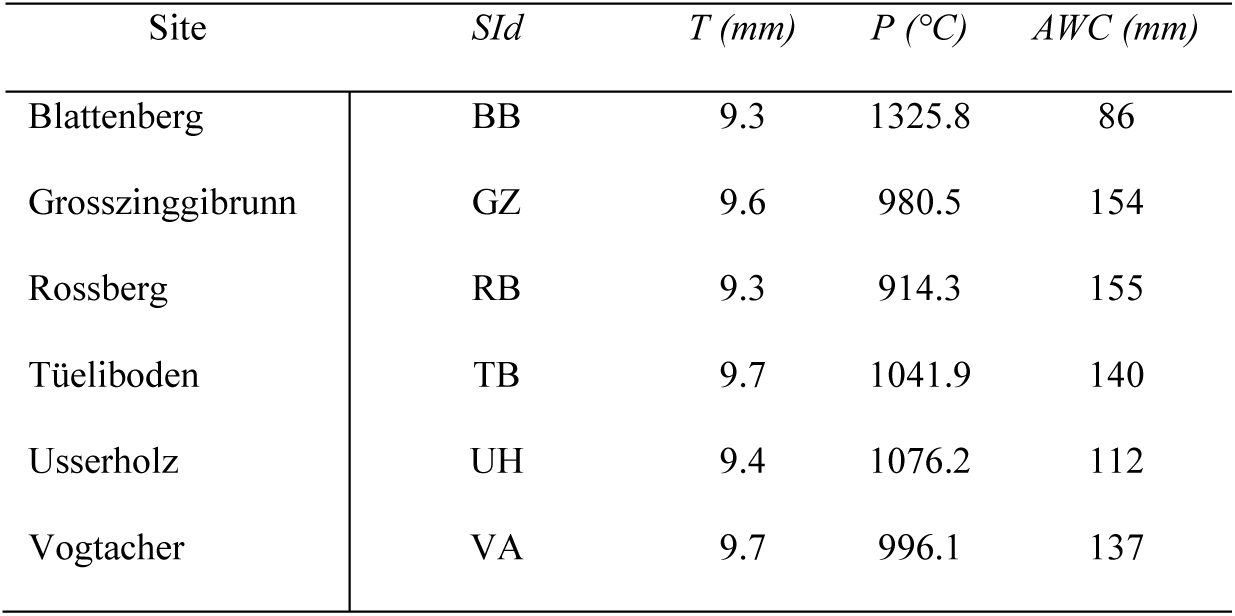
Summary of site-specific properties for the six beech sites (site acronym, SId) on the Swiss Plateau: annual mean temperature (T, °C), annual precipitation sum (P, mm) for the period 1981-2022 (data source: MeteoTest) and available water capacity (AWC, cm) estimated using pedotransfer functions.

| Site | SId | $T$ (mm) | $P$ (°C) | $AWC$ (mm) |
| --- | --- | --- | --- | --- |
| Blattenberg | BB | 9.3 | 1325.8 | 86 |
| Grosszinggibrunn | GZ | 9.6 | 980.5 | 154 |
| Rosserberg | RB | 9.3 | 914.3 | 155 |
| Tüeliboden | TB | 9.7 | 1041.9 | 140 |
| Usserholz | UH | 9.4 | 1076.2 | 112 |
| Vogtacher | VA | 9.7 | 996.1 | 137 |

Additionally, we included the Lägeren site, which is part of the European ICP Level II network (Etzold et al., 2010), is also dominated by *Fagus sylvatica* and has comparable ecological characteristics, e.g. an average annual precipitation of 1051 mm (MeteoSwiss, 2023). Importantly, measured data on actual and potential evapotranspiration rates have been collected at Lägeren—data unavailable for the other beech-dominated sites. Including Lägeren thus allows for an assessment of the water balance during recent drought years under environmental conditions analogous to those of the six beech stands.

We also selected a xeric forest dominated by Scots pine (*Pinus sylvestris*) in Visp, located at 650 m a.s.l on a steep north-facing slope in the Valais (Switzerland), to test the generality of our framework. This site is also part of the ICP Level II network (Rebetez & Dobbertin, 2004). At Visp, drought-induced mortality events occurred regularly in the past (Bigler et al., 2006) and have been documented in detail since 1996 (Rigling & Cherubini, 1999; Hunziker et al., 2022), thus offering a test bed for our framework in the context of the impacts of frequent and severe water stress on forests in the south-central Alps.

Lastly, to encompass a still broader climatic and edaphic spectrum, we studied multi-species forest dynamics at twelve sites across Switzerland and Germany. They span a gradient from cold-mesic to warm-xeric climates, with mean annual precipitation ranging from 600 to 1350 mm and AWC from 100 to 240 mm (Bugmann & Solomon, 2000).

### 2.2 Tree-ring data and drought indices

We retrieved cross-dated tree-ring width data from Neycken et al. (2022) for the six beech sites. We standardized and detrended the individual tree-ring chronologies using a power-transformed Age-Dependent Spline (Obladen et al., 2021; Neycken et al., 2022). All detrended series were averaged into a site chronology using a bi-weight robust mean for each site (cf. SM B1). We assessed the tree growth response to extreme events using pointer year analysis after calculating z-transformations of the detrended site chronologies based on a bi-weight robust mean to detect extreme events at the stand level (SM B2, Jetschke et al., 2019). All analyses were performed in R (v4.2, R Core Team, 2023) using the packages *pointRes* (van der Maaten-Theunissen et al., 2021) and *dplR* (Bunn, 2008, 2010).

We selected a range of drought indices with varying levels of complexity. From the most widely and commonly used single-variable indices, we used the *Standardized Precipitation Index* (SPI, McKee et al., 1993), which is defined as the difference of precipitation from its mean over a specific time scale divided by its standard deviation. Moving towards higher complexity, we included the *Climatic Water Balance*, defined as the difference between monthly precipitation and monthly potential evapotranspiration (Thornthwaite, 1948; Thornthwaite & Mather, 1954). We furthermore selected the *Standardized Precipitation Evapotranspiration Index*, SPEI (Beguería et al., 2014), computed as the standardized difference between precipitation and potential evapotranspiration. We also used the *self-calibrated Palmer Drought Severity Index*, scPDSI (Wells et al., 2004), which takes into account AWC. Lastly, we evaluated the *ForClim drought index* (hereafter termed ‘ForClim’, Bugmann & Solomon, 2000), which additionally distinguishes between tree water supply and demand (see section *ForClim model* below). It is computed at a monthly time step during either the growing season (for deciduous species) or across the year (for evergreen species), provided that the temperature is sufficiently high for tree growth.

The R package *spei* (Beguería et al., 2014) was used to calculate SPI, CWB and SPEI. A modified version of the *pdsi* R wrapper function was used to calculate scPDSI (Zang, 2018). The ForClim drought index was calculated using the R package *ForDrought* (cf. Data Availability).

To consistently compare these drought indices, we computed all indices at a temporal resolution of one month and averaged the monthly values for different periods: autumn (Sept through Nov), winter (Dec through Feb), spring (March through May), summer (June through Aug), the complete growing season (April to October), and the entire year (annual). We standardized the scPDSI and ForClim drought indices using a z-transformation to facilitate direct comparison with the indices that are standardized by definition (i.e., SPI and SPEI). We further rescaled all indices according to McKee et al. (1993) to assign drought severity classes within the following ranges: *mild* (-0 – -0.99), *moderate* (-1 – -1.49), *severe* (-1.5 – -1.99) and *extreme* (≤ -2).

To assess the strength of the relationship between tree growth and drought intensity, we computed the bootstrapped Spearman correlation between the six detrended residual tree-ring chronologies (ring width indices, hereafter RWI, Figure SM B2) and the detrended drought indices (Table SM B1) in the common period 1980–2020 for the selected temporal extents and both the previous and the current year. To identify autocorrelation, we utilized residual tree-ring chronologies and conducted a Mann-Kendall test to evaluate the extent of long-term trends in the predictors (i.e., the drought indices). When autocorrelation was found, we detrended the series using Seasonal Trend Decomposition based on Loess (hereafter STL, Cleveland et al., 1990; for full details cf. SM B3), which effectively separates a time series into its seasonal trend and residual components.

### 2.3. Soil moisture models of contrasting complexity: LWFBrook90 and ForClim

We evaluated the complexity required to simulate soil water dynamics regarding atmospheric water demand (potential evapotranspiration, hereafter PET) and supply (soil moisture and actual evapotranspiration, hereafter SM and AET, respectively) by comparing the performance of the single-layer soil moisture balance model in ForClim (for details, see below) and the detailed multi-layer Soil-Vegetation-Atmosphere Transfer (SVAT) model LWFBrook90 (Hammel & Kennel, 2001). LWFBrook90, an extension of Brook90 (Federer, 2002; Federer & Lash, 1978), simulates daily transpiration and interception from the canopy using a single plant layer (“big leaf” approach), as well as soil water fluxes by numerically solving the Richards equation. In contrast to ForClim, LWFBrook90 does not impose a field capacity constraint, allowing soil moisture to exceed this threshold and enabling a more detailed simulation of soil water dynamics. This fundamental difference presents a considerable challenge when comparing the behavior of the two models (Guswa et al., 2022).

In a first step, we compared the measured above-canopy AET at the site Lägeren with simulated AET. Simulated AET data for LWFBrook90 were obtained from Meusburger et al. (2022) for the period 2013-2019. For ForClim, we simulated AET by forcing the model with measured temperature and precipitation obtained from MeteoTest (2020).

In a second step, we simulated soil moisture, PET and AET at the six beech-dominated sites and compared model behavior. LWFBrook90 was forced with daily weather time series (MeteoTest, 2020); elevation and slope angle were provided from the digital elevation model DHM25 (spatial resolution of 25 m; Swisstopo, 2004). Additionally, Leaf Area Index (hereafter LAI) data were obtained from the MODIS global Leaf Area Index and Fraction of Photosynthetically Active Radiation (FPAR) product (Myneni et al., 2021), while stand height was retrieved from the global forest canopy height model by Potapov et al. (2020). Full details on the simulation setup and parameters are provided in SM C (cf. Table C2.1). We calculated AWC from soil texture properties (Baltensweiler et al., 2021) using the pedotransfer function of Wessolek et al. (2009) (cf. sections D1 and D2 in SM, Eq. 10-12).

Model performance was evaluated as follows: at Lägeren, we computed the root mean squared error (RMSE) and the mean absolute error (MAE; cf. *Eq. 1-2*), while for the six beech sites where no soil moisture data were available, we assessed the similarity of the simulated water balance variables using cumulative density functions (CDFs), in particular cumulated soil moisture (CSM), and the Bland Altman method (BAM), which is commonly applied in medicine and genomics (Giavarina, 2015; Bland & Altman, 1986). BAM aims to describe the degree of agreement between two methods or data sources. The y-axis of BAM plots represents the difference between each pair of simulated variables from the two methods, whereas the x-axis shows the mean of each pair of values among the two models (*Eq. 3*). The mean bias (β, *Eq. 4*) and the limits of agreement (*Eq. 5*) were also calculated, where σ is the standard deviation of the differences in each pair of values. According to BAM, 95% of the scatter points should reside within the limits of agreement, representing ±1.96σ from the mean difference between the data of the two models. If the mean difference of the data is not significantly different from zero (based on a one-sample t-test), this indicates good agreement between the two methods.

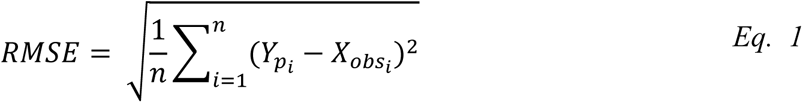

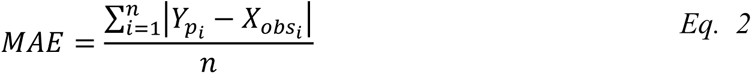

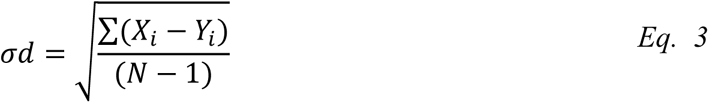

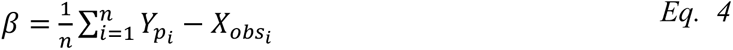

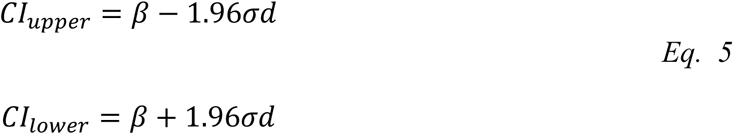

### 2.4 ForClim Model

#### 2.4.1. Overview

ForClim is a forest gap model (Botkin et al., 1972) originally designed to capture the long-term (i.e., decades to centuries) growth, mortality and regeneration of trees in temperate forests of central Europe and account for climate change effects on forest dynamics (Bugmann, 1994, 1996). In gap models, forest dynamics are simulated on small areas (‘patches’), usually with a size of 400-1000 m^2^, each representing one out of many stochastic realizations that are spatially independent of each other. The soil water balance is calculated using a monolayer “bucket” model that stores all incident water until its capacity (“bucket size”, kBS) is reached, which corresponds to AWC. The bucket model has a fixed field capacity, beyond which soil moisture cannot increase. This simplification facilitates computations but limits the representation of hydrological processes such as percolation and soil moisture variability above field capacity. For details, cf. Bugmann & Cramer (1998) and Bugmann & Solomon (2000).

This bucket model is directly connected to plant dynamics, as tree growth is determined as a species-specific potential (i.e., under optimum conditions) that is reduced via growth-reduction factors (*GRF*) accounting for light availability, crown condition, temperature, nitrogen and soil water supply vs. demand (Bugmann & Solomon, 2000). Tree mortality from stress is assumed to occur when diameter growth is below a specific stem diameter increment or a fraction of the maximum increment for several years (Solomon, 1986). Thus, *predisposing* stress is assumed to increase the mortality rate, but there is no formulation of *inciting* stress (cf. the PIC scheme).

#### 2.4.2. Integrating predisposing and inciting stress factors

To enhance the response of tree growth to environmental extremes (temperature and soil water dynamics), we modified the original growth reduction formulation (Huber et al. 2020) by applying Liebig’s “law of the minimum” for the temperature (DDGF) and soil moisture (SMGF) growth factors (Eq. 6; cf. Liebig et al., 1842), rather than multiplying them:

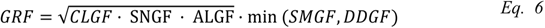

where GRF is the overall growth reduction factor, and CLGF, SNGF and ALGF are the crown-related, nitrogen-related and light-related growth factors, respectively. The underlying rationale is that in the temperate and boreal regions, conditions are typically either dry *or* cold, but not dry *and* cold (cf. Bugmann, 1996b). To preserve the species-specific response to environmental stress although the level of GRF is changing due to the use of *Eq. 6*, we had to re-estimate a temperature-related species-specific parameter (for details cf. SM D6), and we also modified the dynamic formulation of site index, which reflects the response of maximum tree height to temperature and drought and their changes over time (for details cf. SM D7).

To disentangle predisposing and inciting factors leading to drought-related mortality, we identified short-term (within a year) and long-term (multi-year) stressors linked to drought duration and intensity as well as carbon starvation, as explained below.

First, we accounted for the effect of long-lasting droughts by *a predisposing factor* using a drought memory term (*DrM*; Wang et al., 2012; *Eq. 7*):

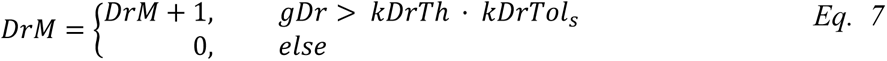

This formulation counts all continuous years in which drought intensity, represented by the ForClim drought index (*gDr*), exceeds a fraction (*kDrTh*) of the species-specific drought tolerance (*kDrTol_s_*). In this manner, we account for the species-specific resistance to multi-annual drought stress. To be consistent, we also modified the “slow growth” factor 𝑆𝐺𝑟 to better mimic the impact of carbon reserves on mortality risk (for details see SM D3).

Second, to deal with the inciting factor we defined drought duration within any given year (*gDrD*, *Eq. 8*) as the ratio of the number of dry months relative to the total number of months *m* of the growing period (for deciduous species, 𝑚_𝑔𝑝_; annual for evergreen species, 𝑚_𝑎𝑛_). The algorithm selects those months in which average temperature (*T_m_*) is above a threshold *kJ* (5.5 °C) and water supply (i.e., transpiration, *gE_m_*, cm) relative to water demand from the soil (*gDm*, cm) is below a threshold *kEg*; i.e., *gE_m_*/*gD_m_* < *kEg*; and *SM_s_* < *kBS*). The term 𝟙 represents an indicator function, which equals 1 if the condition is true for a given month, and 0 otherwise.

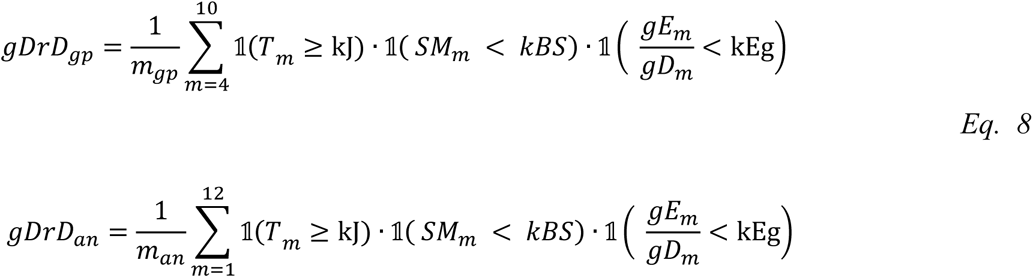

Third, under conditions of low demand (i.e., in spring and fall) this index would not record any drought, although soil moisture (*SM_m_*) may be limiting e.g. for bud break in spring. Thus, we used two variables (*Eq. 9*) to capture limiting soil moisture levels in spring and fall along with a threshold (*kDu*) for the duration of the drought to define an *inciting* factor for drought stress (*IncFDr; Eq. 9*).

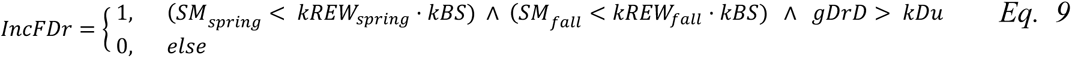

This formulation for the seasonal water deficit in spring and fall is based on the concept of ‘relative extractable water’ (REW; Bréda et al., 2006; Granier et al., 1999; SM D4). Specifically, the spring component of *IncFDr* reflects the need of trees to mobilize water for bud break and cell division and elongation, while the fall component reflects the need of accumulating carbon reserves for the subsequent year (Figure 3). The threshold *kDu* was set to 0.28, corresponding to two months out of a seven-month growing period (2/7 ≈ 0.28) for broad-leaves, and three to four months out of the whole year (3.5/12 ≈ 0.29) for evergreen species, provided that winters are warm enough (cf. Hidy et al., 2021; Merganičová, 2023). The seasonal soil moisture levels (*SM_fall_, SM_spring_*) are calculated for the fall (September to November) and spring (March to May) periods for both evergreen and deciduous species.

**Figure 2:**
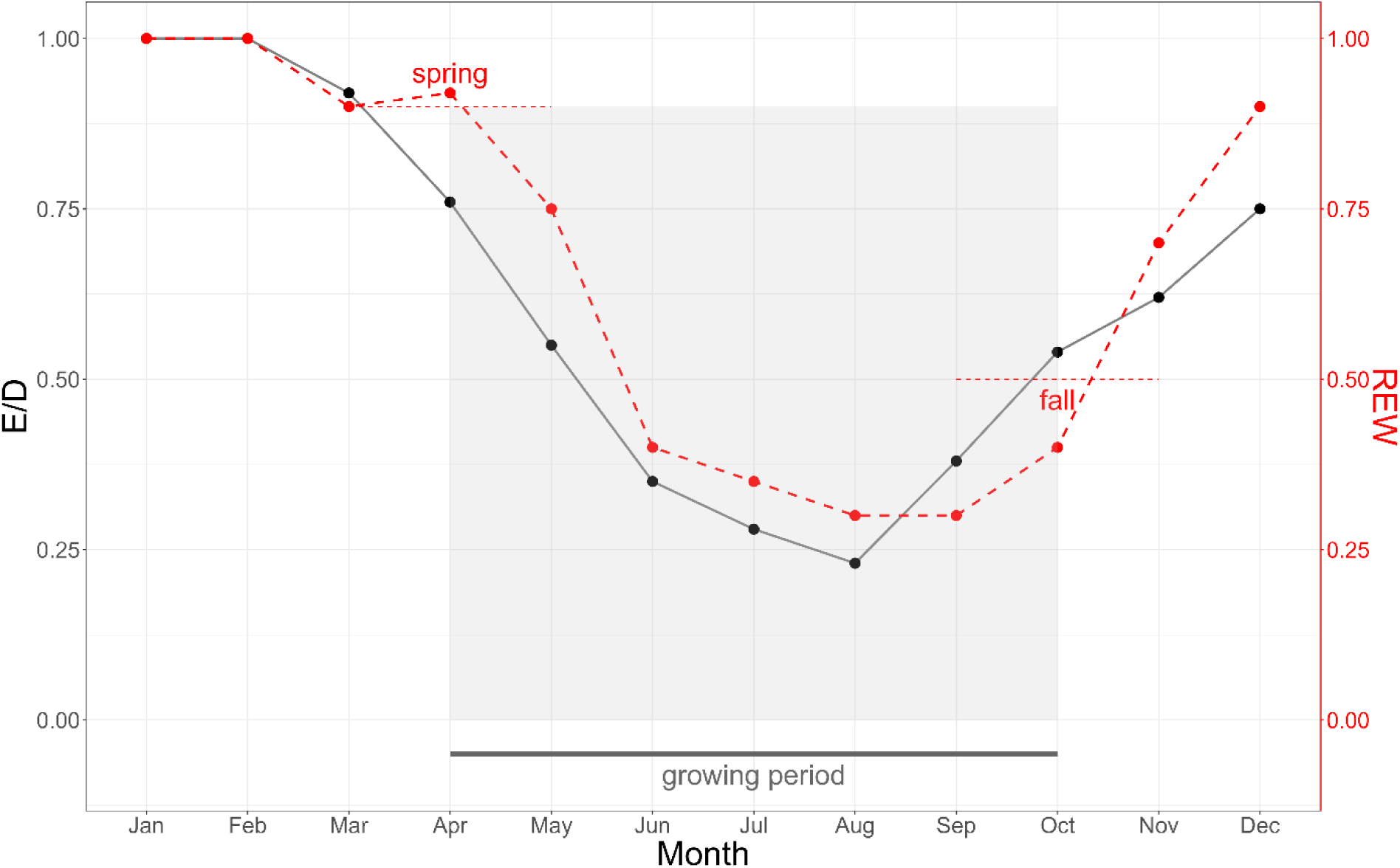
Inciting factor scheme showing the stress factors operating in the short term (one year): during the growing period, the formulation selects only those months in which (1) water demand (ratio of Evapotranspiration E on the water Demand D, unitless) is sub-optimal (<0.9), and (2) soil moisture falls below a threshold in spring (0.9) and fall (0.5). Both axes are unitless.

**Figure 3:**
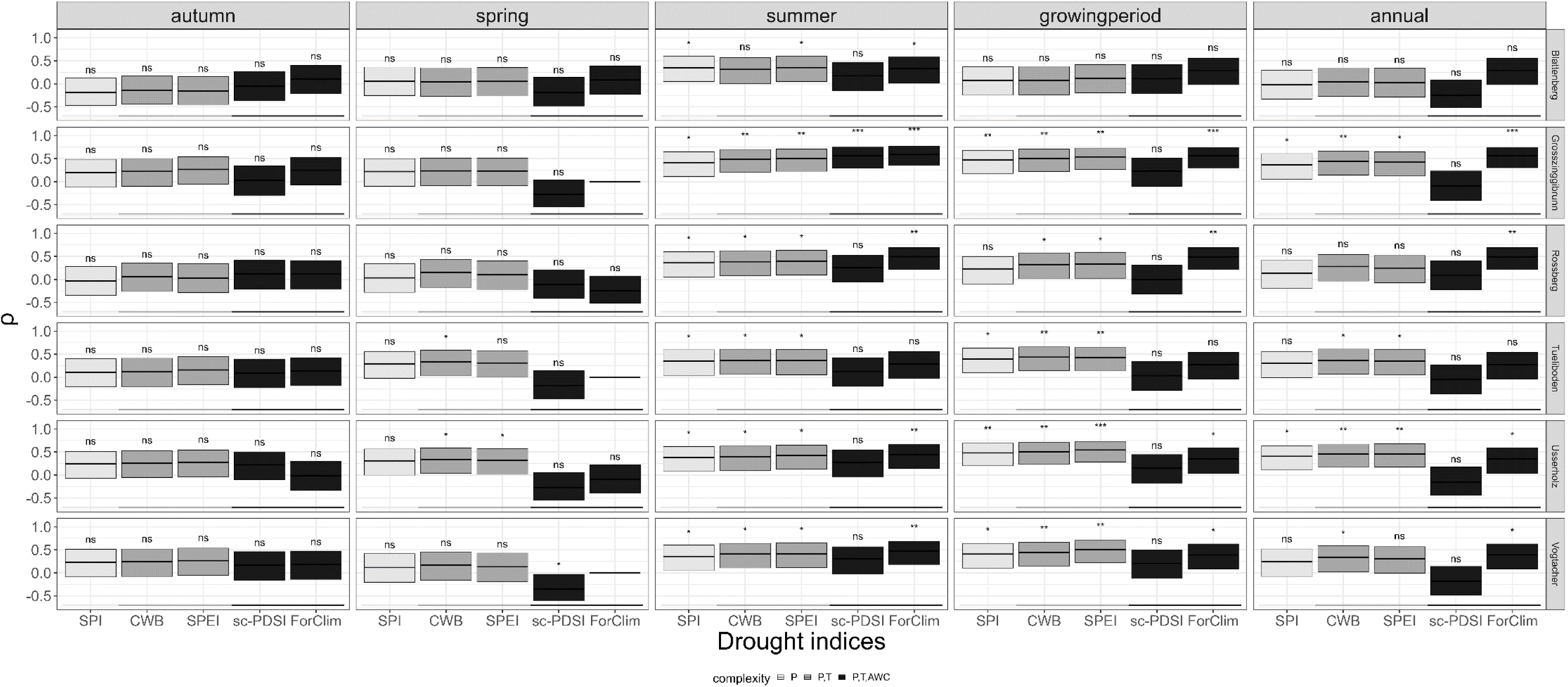
Seasonal Spearman rank correlation between detrended tree-ring indices and detrended drought indices of increasing complexity (from indices including precipitation (P) only (SPI) to precipitation and temperature (CWB, SPEI) to precipitation, temperature and available water capacity (AWC; scPSDSI, ForClim). The significance levels (p-values) are indicated by symbols (“***”, “**”, “*”, “ns” from 10^-3^ to >0.05). When no variance was observed, an intercept y=0 was used (i.e. at Grosszinggibrunn, Tüeliboden and Vogtacher for the ForClim drought index).

Lastly, the overall stress-induced mortality probability (*gPStr*), including the carbon memory and integrating predisposing as well as inciting factors, is formulated as follows:

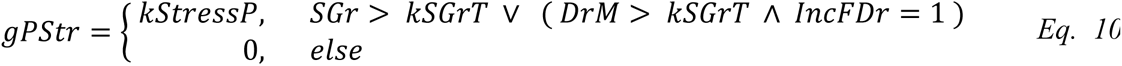

where 𝑘𝑆𝑡𝑟𝑒𝑠𝑠𝑃 is the stress-induced enhanced mortality probability, 𝑆𝐺𝑟 is the slow-growth counter, and 𝑘𝑆𝐺𝑟𝑇 indicates the number of stress years that are tolerated until mortality probability is enhanced (Peltier et al., 2023). The first term of *Eq. 10* (*SGr* condition) captures the probability that a tree may die due to slow growth induced by whatever cause (e.g., insufficient light), whereas the second term (*DrM* and *IncFDr* conditions) reflects that a string of dry years can enhance mortality under a particularly prolonged summer drought coupled to early- and/or late-season soil moisture depletion.

The ensemble of these features gives rise to ForClim v4.1. Since contributing factors (Manion, 1981) such as insect damage are currently not included, we refer to the concept underlying ForClim v4.1 as the “PI framework”, rather than PIC. Note that none of the parameters of the new drought-related mortality framework (Eqs. 7-10) were calibrated against any data sources, least those used for the simulation studies that are explained below.

#### 2.4.3 Model initialization

**Six beech sites** – We compiled forest stand data using relascopic sampling (Grosenbaugh, 1958, Bitterlich, 1948) in combination with the single-tree inventory conducted by Neycken et al. (2022) in 2019 and 2020, resulting in a total of 14 small sampling plots of 10 m radius for Blattenberg, 35 for Grosszinggibrunn, 14 for Rossberg, 10 for Tüeliboden, 11 for Usserholz, and 36 for Vogtacher. For each site, we calculated the mean basal area and its standard deviation for the respective years (cf. Table A1, SM A). Individual DBH data were collected for the target trees, defined as those sampled by Neycken et al. (2022) for dendrochronological analysis and crown vitality assessment. Additionally, DBH measurements for neighboring trees within a 10 m radius of each target tree were included in the dataset (Neycken et al., 2022). To further enhance the inventory data, we incorporated information from trees felled following observed mortality at each site. These records were used to retrospectively reconstruct stand structure from 2018 back to 2000, which was employed to initialize the model for the year 2000. This process was carried out by reintroducing the dead trees into each of the sampled plots at the start of the simulation time, and the same process was applied for the neighboring trees.

For model initialization, the inventory data of vital, declining, and dead/cut trees were randomly distributed on *n* = 200 patches, each with a size of 0.085 ha, as simulations need to be replicated to represent *n* stochastic realizations of the processes simulated in ForClim (cf. Bugmann, 2001). The simulations were forced by interpolated, site-specific monthly meteorological time series of precipitation and temperature (MeteoTest, 2020), while soil properties were obtained from Baltensweiler et al. (2021). For each site, AWC (*kBS*) was calculated using the Wessolek pedotransfer function (cf. Eqs. 10-12 in SM D1 and D2). These stands are experiencing high atmospheric nitrogen (N) deposition (Roy et al., 2021), and thus we set N availability to a high value of 180 kg ha^-1^, assuming N not to be limiting (cf. Table S7, SM D7).

**ICP Level II sites** – For both Lägeren and Visp, the state of the forest was determined by simulating Potential Natural Vegetation from bare ground (i.e., no stand initialization) to equilibrium under current climate conditions, assuming no management. At Lägeren, this state was used to initialize the simulations for the period 2011-2020. Similarly, for Visp we forced the model with precipitation and temperature time series from the LWF Visp meteorological station for the period 1997-2004 (Haeni et al., 2019).

**European climate gradient** (large-scale Potential Natural Vegetation) – We performed simulations starting from bare ground over 200 patches (patch size 0.08 ha) for 1500 years to determine the equilibrium between forest vegetation and climate. We averaged the simulated species-specific basal area between the simulation years 1300 and 1500 (for details cf. SM D8). The weather time series were obtained by randomly sampling years from the site-specific climatology (Bugmann, 1994, 1996; cf. SM D8).

## 3. Results

### 3.1 Selecting the most suitable Drought Index for tree growth response

As expected, we found only weak relationships between tree growth and drought intensity in the mesic mature beech forests: correlations were low to medium, yet with distinctive seasonal differences (Figure 3). Still, the drought indices featured rather distinct behavior across the six sites and seasons, suggesting that it is possible to identify an index that best captures extreme events.

Specifically, there was a tendency for indices of low complexity, particularly SPI, to have lower correlations across all seasons and sites, while indices of higher complexity performed better, yet with notable exceptions. Surprisingly, the scPDSI demonstrated poorer performance compared to simpler indices during spring, the growing period, and the entire year. Similarly, the ForClim drought index exhibited low performance in spring, but outperformed the other indices during summer, the entire growing period and throughout the year. In the autumn period, the ForClim drought index showed the highest, yet non-significant, correlations at three out of six sites.

Non-parametric correlations between the current tree-ring index and the previous year’s drought index exhibited consistent patterns across sites, yet they were non-significant for all seasons (Figure SM B3). Notably, ForClim showed the highest correlations during the growing period and at the annual time scale at most sites for the previous year. It also featured significant correlations in autumn for Grosszinggibrunn and Usserholz, followed by Blattenberg, and in spring except for Usserholz. However, due to the zero variance of the ForClim index at three of the beech sites—Grosszinggibrunn, Tüeliboden, and Vogtacher—correlations could not be calculated for the current and previous year at these sites.

Overall, our analysis indicated that ForClim is capable of picking up the relatively weak drought signals at many of these mesic beech sites, being the most suitable index for the entire growing period and the entire year; all indices performed similarly for the summer, and – importantly – the use of any of these indices is discouraged for spring and autumn.

### 3.2 Simulating the water balance: how much complexity is needed?

At the site Lägeren, observed versus simulated AET indicated that both ForClim and LWFBrook90 captured the seasonal cycle of AET well (Figure 4A,B), yet with distinct differences, surprisingly indicating a somewhat lower performance of the more complex model, LWFBrook90, compared to ForClim, regardless of the metric being used (Table 2). Meusburger et al. (2022) attributed the performance problems of LWFBrook90 to the omission of lateral water fluxes at Lägeren, which resulted in lower simulated AET (Figure 4A). The ForClim model almost consistently underestimated AET, especially during winter. However, it performed well during the dry years of 2015 and 2018-2019, showing a reduction in AET, particularly in the summer months. In contrast, the LWF-Brook90 model featured both under- and overestimations of AET (Figure 4B). Notably, it significantly overestimated AET during the summer months of 2015-2016 and 2018-2019, followed by a substantial reduction in fall.

**Figure 4:**
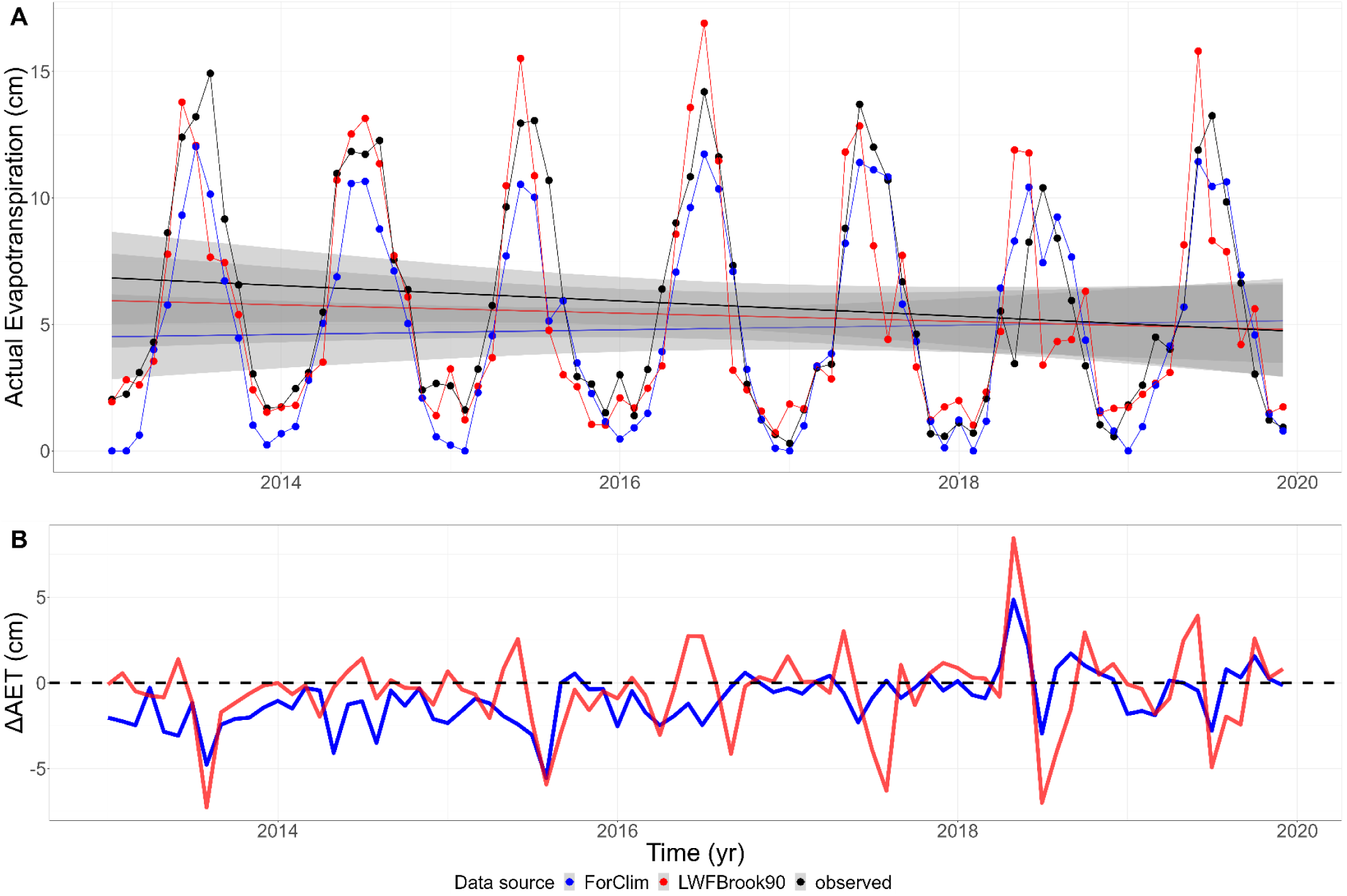
**(A)** Simulated and observed monthly actual evapotranspiration (AET) at the ICP Level II site Lägeren (Switzerland). The trend lines in panel A indicate a progressive reduction in simulated and measured AET from 2015. Both models showed a slight negative temporal trend in simulated AET. **(B)** Differences between simulated and observed AET across years. Observed data are indicated with a black line (A) and a dashed line (B), respectively.

**Table 2.** Summary statistics of observed vs. simulated AET at the site Lägeren across the years 2012-2020 (observational data from Meusburger et al., 2022). *MSE* = Mean Squared Error; *MAE* = Mean absolute Error; *RMSE* = Root Mean Squared Error.

| Model | $MSE$ | $MAE$ | $RMSE$ |
| --- | --- | --- | --- |
| ForClim | 3.33 | 1.39 | 1.82 |
| LWFBrook90 | 5.74 | 1.61 | 2.4 |

Soil moisture simulated by ForClim at the six beech sites was consistently higher than for LWFBrook90 (Figure 5A) except for Blattenberg (cf. Fig.SM C2.1), where simulated soil moisture was larger in LWFBrook90 (at average +16%). For Tüeliboden, ForClim yielded higher values (+17%), and considerably higher values for Grosszinggibrunn, Usserholz and Vogtacher (+28 to +30%; cf. Table 3). The higher soil water availability simulated by ForClim occurred mostly during the cold seasons, i.e. autumn and winter (cf. SM C, Fig. C2.2).

**Figure 5:**
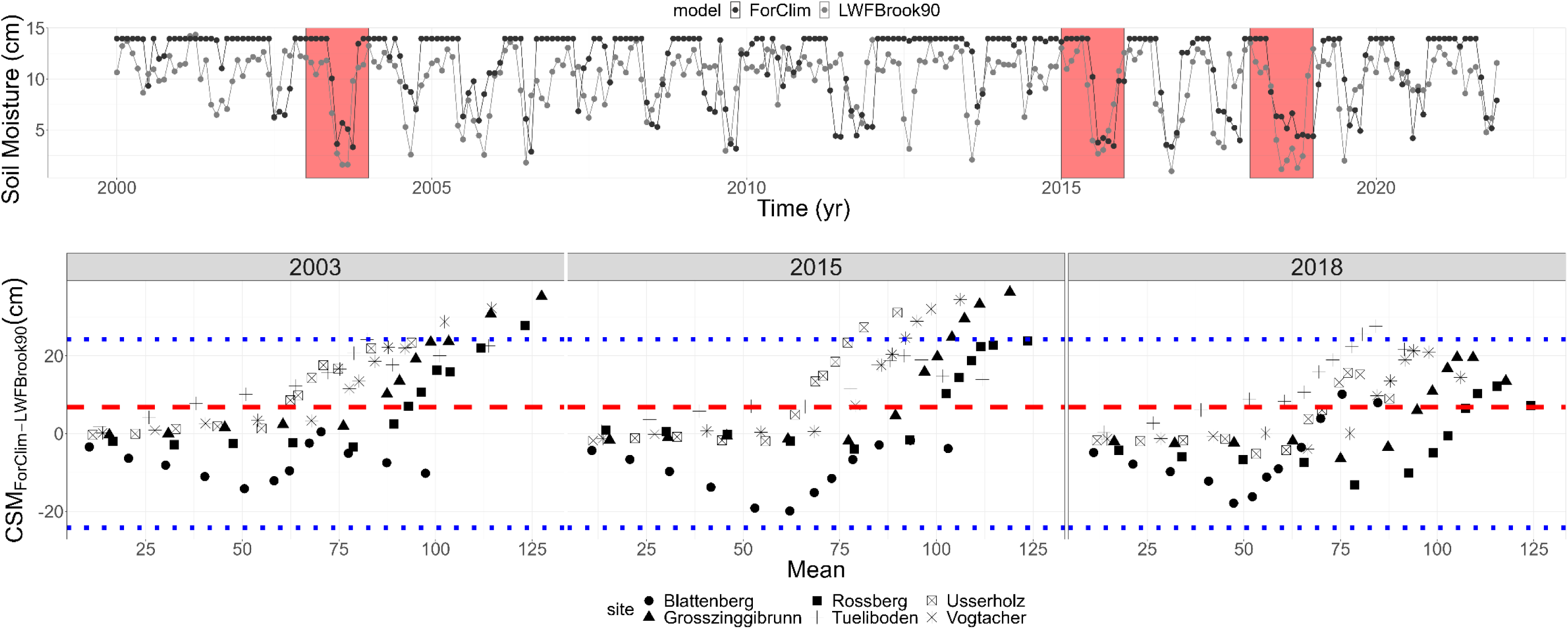
Simulated soil moisture with LWFBrook90 and ForClim. **(A)** Monthly variation of simulated soil moisture at the site of Tüeliboden for the model ForClim (black line and black dots) and LWFBrook90 (grey lines and grey dots). The highlighted red areas indicate the driest years 2003, 2015 and 2018. **(B)** Differences of Cumulative Soil Moisture (CSM, cm on the y-axis) in Bland-Altmann plots for the driest years (2003, 2015, 2018) at the six beech-dominated sites (dots with different shapes) simulated with ForClim and LWFBrook90. The x-axis shows the mean of each pair of values among the two models. The limits of agreement are indicated with the blue dashed intercepts. The intercept on the y-axis indicated by the red dashed line in Figure 5B shows the mean bias. Positive deviations from the mean bias between ForClim and LWFBrook90 indicate a higher simulated soil moisture for ForClim compared to LWFBrook90.

**Table 3.** Ratio of Cumulative Soil Moisture (CSM) between the ForClim and LWFBrook90 models (ForClim:LWFBrook90) for each year until the month of December, averaged across the years 2000-2020 along with standard deviations.

| Site | $CSM_{\text{ForClim:LWFBrook90}}$ |
| --- | --- |
| Blattenberg | $0.86 \pm 0.09$ |
| Grosszinggibrunn | $1.28 \pm 0.12$ |
| Rosserberg | $1.24 \pm 0.12$ |
| Tüeliboden | $1.17 \pm 0.12$ |
| Usserholz | $1.3 \pm 0.14$ |
| Vogtacher | $1.3 \pm 0.11$ |

The cumulative density functions (CDFs) for AET showed good agreement between the models at Grosszinggibrunn, Rossberg and Vogtacher, but ForClim produced notably lower values at Usserholz, Blattenberg, and Tüeliboden. For PET, slightly lower values were observed from ForClim across all sites, ranging from -2.93 cm (Vogtacher) to -5.83 cm (Usserholz) (cf. Fig. SM C2.3).

The Bland-Altmann analyses (Figure 5B) revealed varying degrees of agreement between ForClim and LWFBrook90 in simulating cumulative soil moisture (CSM, cm) across the six beech sites and across the years. The models showed closer alignment during dry years (2003, 2015, and 2018; Figure 5B) compared to wetter years, albeit with year-specific differences (cf. Fig. SM C2.4). Notably, in 2018 —the driest year — the points were more tightly clustered around the zero-bias line compared to 2003 and 2015, indicating higher consistency between the models during this extreme drought. Moreover, no significant difference was found between the mean CSM simulated by the two models for 2018 (t = 1.02, p = 0.21), although some site-specific discrepancies were evident, particularly at Tüeliboden.

These results suggest that both models capture the same trend of soil moisture depletion during extreme droughts, and the ForClim model tends to yield higher soil moisture in wetter periods and lower values in drier periods at some sites compared to the more complex LWFBrook90. Overall, LWFBrook90 featured a less conservative behavior with soil moisture, which likely is a result of its daily resolution, thus better capturing extreme situations. ForClim also captured the summer drying of the soil and the extreme summer droughts in the years 2003, 2015, and 2018, while simulated soil moisture remained close to saturation for the remainder of the year (cf. Fig. SM C2.2).

### 3.3 Is tree mortality driven by compound stress events?

The previous model version, ForClim v4.0.1, lacked the capability to simulate drought-induced mortality at the six beech-dominated sites, whereas the implementation of the PI framework in ForClim v4.1 enabled us to quantify this phenomenon well, except for the site Blattenberg (Figure 6). In both ForClim versions, the simulated basal area showed no drought-related mortality events in the period 2000-2017, not even in the very dry years 2003 and 2015, whereas ForClim v4.1 featured a notable reduction starting in 2018 (Figure 6). These simulated mortality events are consistent with observations, yet with some site-specific over- and underestimation. The simulations for Rossberg featured the lowest MAE and RMSE among sites (Table 4), indicating the highest accuracy, followed by Usserholz and Tüeliboden, suggesting an overall good to medium level of consistency in prediction errors across sites. Grosszinggibrunn and Vogtacher featured a higher variability (dispersion of errors) compared to the other sites. Blattenberg featured the lowest accuracy and only a slight decrease in basal area, i.e. only little mortality.

**Figure 6:**
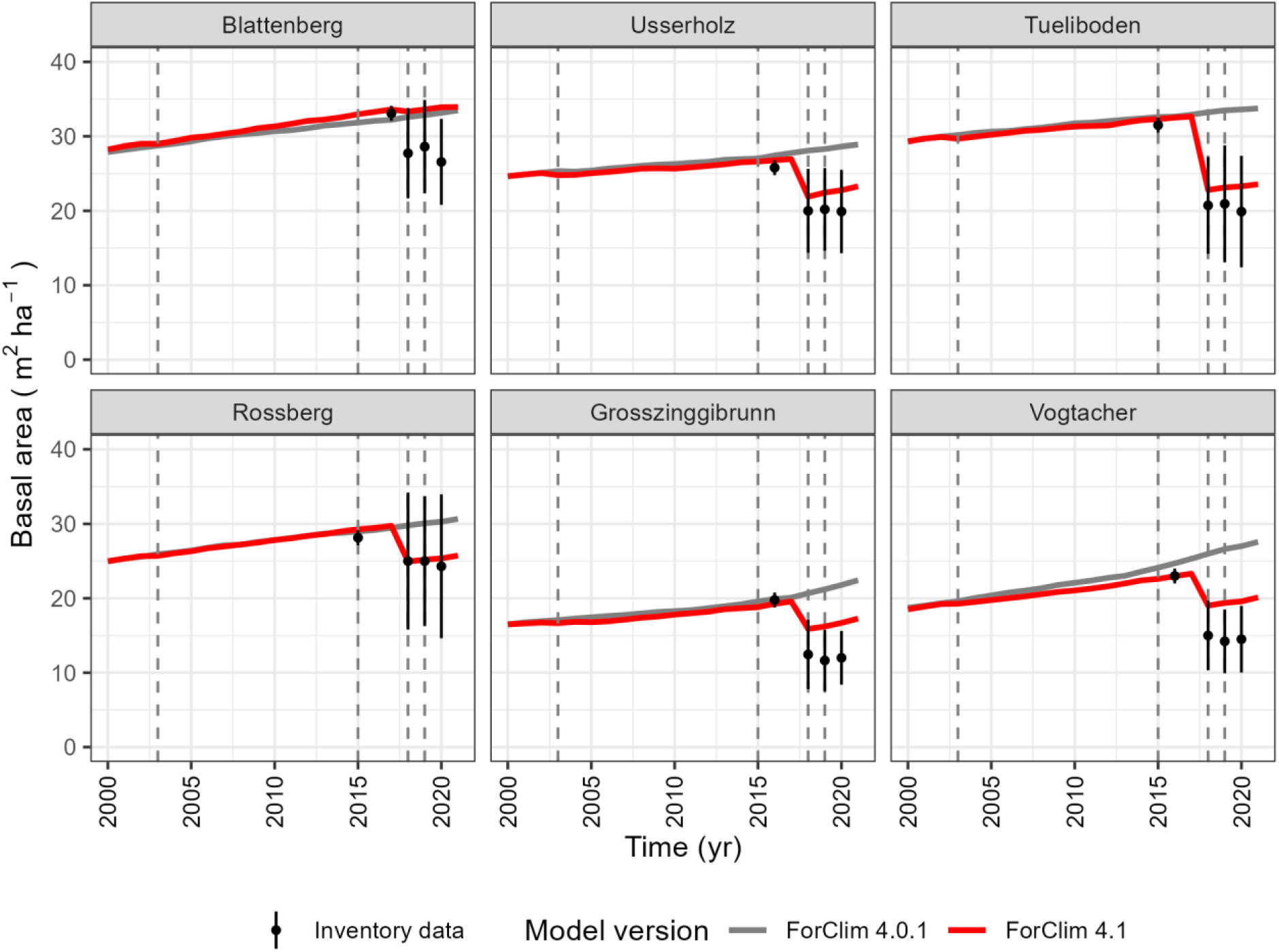
Simulated basal area of *Fagus sylvatica* over time and observed basal area (black dots) in the years 2018-2020 (Neycken et al., 2022) with the newly developed predisposing and inciting factor scheme (ForClim 4.1) and the earlier model version (ForClim 4.0.1). The dashed lines indicate the drought years 2003, 2015, 2018 and 2019.

**Table 4.** Model statistics for the years 2018, 2019 and 2020 (data source for inventory: Neycken et al., 2022) and simulated basal area losses at the six beech sites expressed as the percent changes from the long-term mean (2000-2015 for Rossberg and Tüeliboden, 2000-2016 for Grosszinggibrunn, Vogtacher and Usserholz, 2000-2017 for Blattenberg).

| *Site* | *MAE* | *RMSE* | *Year* | *$\Delta BA_{sim}$ (%)* |
| --- | --- | --- | --- | --- |
| Blattenberg | 5.97 | 6.05 | 2018 | 7.9 |
|  |  |  | 2019 | 8.8 |
|  |  |  | 2020 | 9.7 |
| Usserholz | 2.34 | 2.37 | 2018 | -14.4 |
|  |  |  | 2019 | -12.3 |
|  |  |  | 2020 | -11.0 |
| Tüeliboden | 2.55 | 2.62 | 2018 | -25.8 |
|  |  |  | 2019 | -24.7 |
|  |  |  | 2020 | -24.2 |
| Rossberg | 0.44 | 0.62 | 2018 | -7.9 |
|  |  |  | 2019 | -7.0 |
|  |  |  | 2020 | -6.4 |
| Grosszinggi-brunn | 4.23 | 4.27 | 2018 | -9.5 |
|  |  |  | 2019 | -7.6 |
|  |  |  | 2020 | -4.9 |
| Vogtacher | 4.75 | 4.78 | 2018 | -7.8 |
|  |  |  | 2019 | -6.1 |
|  |  |  | 2020 | -5.1 |

The reduction of simulated basal area in ForClim v4.1 across all six beech sites indicated a consistent trend over the three driest years (Table 3). Tüeliboden had the strongest decrease compared to the previous years, followed by Usserholz and Grosszinggibrunn. Vogtacher and Rossberg showed a smaller decrease than the other sites, while an increase of simulated basal area was observed in Blattenberg in the year 2018.

When simulating stand dynamics at the ICP-Level II site of Visp (Figure 7), basal area and stem number featured a strong reduction in the years 1999 and 2003. This pattern is in line with the mortality observed by Dobbertin et al. (2004) and Hunziker et al. (2022) for that same period, amounting to ∼75.6% from 1997 to 2004.

**Figure 7:**
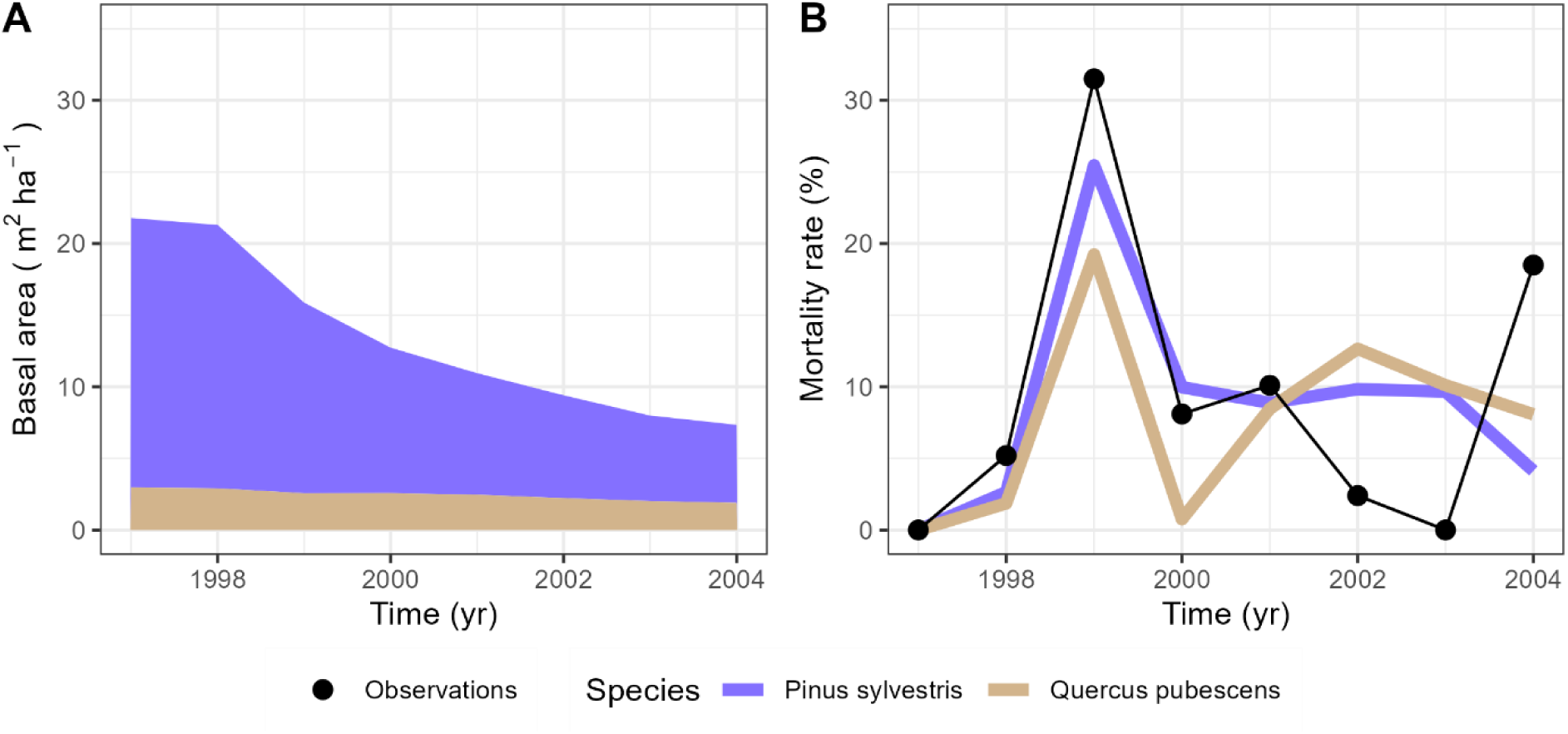
Simulated basal area (**A**) and mortality rate (**B**) over the years 1997-2004 at the site Visp for *Scots pine* and *Quercus pubescens*. The black dots indicate the observed mortality rates according to previous studies (Hunziker et al. 2022).

When analyzing model behavior across an extended environmental gradient in Europe, ForClim v4.1 tended to feature lower basal area compared to ForClim 4.0.1 (Figure 8). The differences were particularly revealing when analyzing the species composition along the elevational and climatic gradient, as explained below.

**Figure 8:**
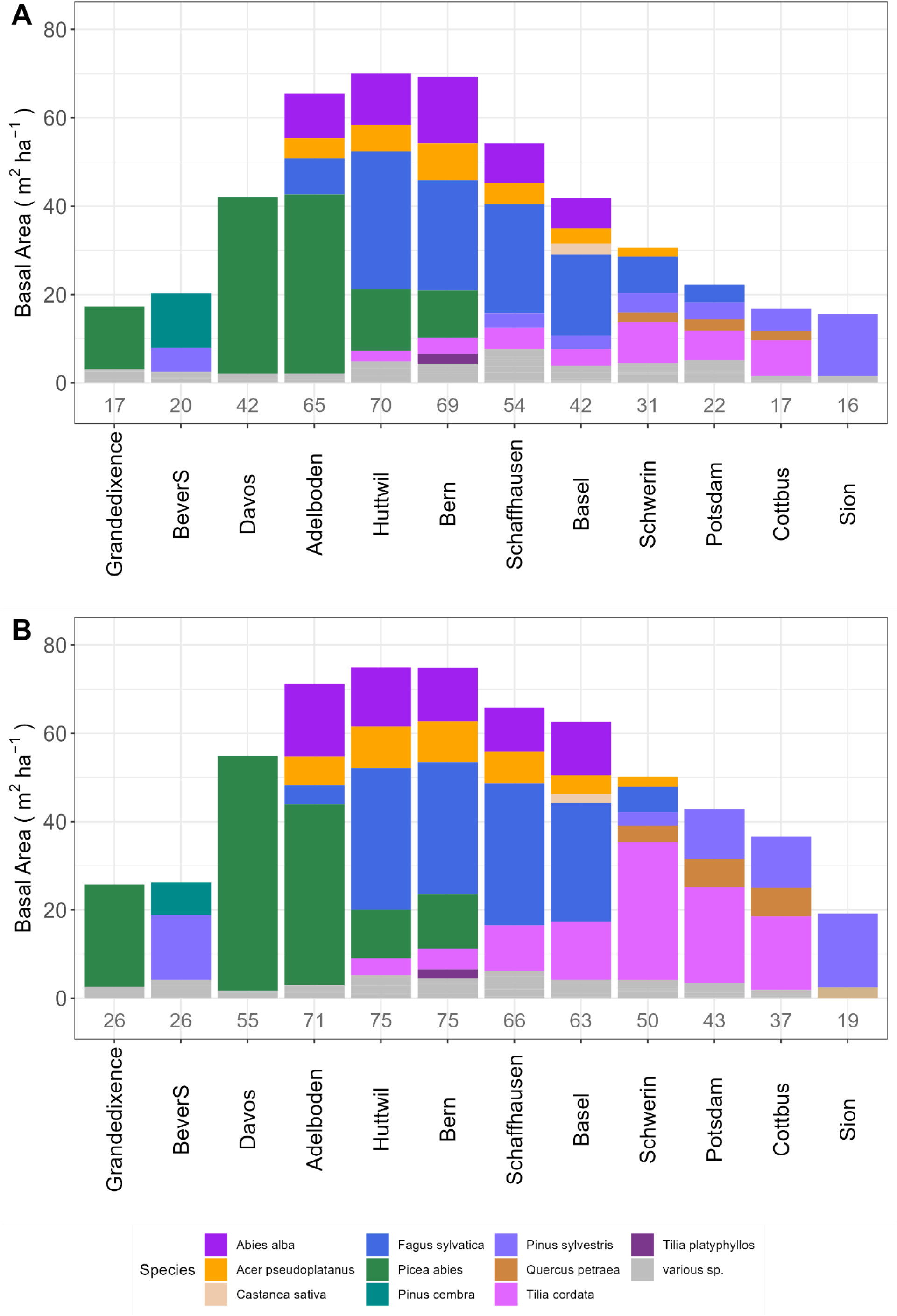
ForClim model version comparison along the EU gradient of sites. **(A)** Basal area (BA) at the equilibrium of potential natural vegetation (PNV) simulated with ForClim version 4.1 and **(B)** and with the older ForClim version 4.0.1 across 12 sites in Switzerland and Germany (Bugmann, 1996 b). Species with BA ≤ 2 m^2^ ha^-1^ are summarized in the “various sp.” category.

In ForClim v4.1. the major features of forest basal area and species composition along the extended climatic gradient from cold-wet to warm-dry conditions (Figure 8A) were retained compared to the predecessor version 4.0.1 (Figure 8B): Conifers dominate the high-elevation cold-wet sites (Grande Dixence to Adelboden), mixed conifer-deciduous forests are found at mid-elevations, which also feature the highest basal area along the gradient. Beech (*Fagus sylvatica*) dominance extends from Huttwil to Basel, and towards the dry edge of the gradient drought-adapted species like oaks (*Quercus* spp.) and lime (*Tilia* spp.) are gaining dominance, ending up with pine dominance towards the dry treeline (Sion).

Yet, there are marked difference in the performance of ForClim v4.1 compared to v4.0.1. Specifically, we found a consistent reduction in the share of *Picea abies* (Grande Dixence, Davos, Bern), and the share of *Abies alba* decreased slightly at the cool-wet sites. Also, a substantial decrease of *Tilia cordata* was found (cf. Figure SM D7). On the contrary, an increase in the share of *Pinus sylvestris* (Schaffhausen, Basel) was evident from ForClim v4.1. At the south-facing site Bever in the continental high Alps, the share of *Pinus sylvestris* was reduced substantially in favor of *Pinus cembra*. Importantly, the share of *Fagus sylvatica* increased moderately in Adelboden, Schwerin and Potsdam, while it decreased somewhat at the drier sites of the Swiss Plateau (Schaffhausen and Basel).

## 4. Discussion

### 4.1 Seasonal drought periods modulate growth responses to drought

Drought indices are often used to capture the relationship between drought intensity and tree growth. Also, they are important for monitoring forest health and resilience as they help to identify areas under drought stress (Zargar et al., 2011; Vicente-Serrano et al., 2012). As expected, we found that different indices perform quite differently. The performance of a drought index is contingent on its complexity, expressed among others by the number of predictors and the temporal extent. In our study, particularly those indices that are used widely in dendrochronology, i.e. SPI and SPEI (Speich, 2019; Schwarz et al., 2020) performed less well at all sites and across different temporal extents. Surprisingly, the Palmer Drought Severity Index performed rather poorly as well, particularly in summer, despite being used as a standard index to capture drought patterns (e.g. as a component of the US Drought Monitor, 2019). Thus, while these indices may be suitable to capture strong droughts in semi-arid or arid areas, they are less suitable to characterize drought occurrence in mesic areas, as shown in the present study.

In contrast, the ForClim drought index, which embeds a simple single-layer soil moisture model, was able to capture the distinct drought intensity signals in summer, across the vegetation period, and at the annual level. These results underline that water availability is key for shaping the response of trees to drought (Speich et al., 2018; Gessler et al., 2022; Klesse et al., 2022; Meusburger et al., 2022). Thus, the ForClim Drought Index has notable predictive capability in detecting drought signals during the growing season even under mesic conditions. This underscores its robustness in regions with moderate drought, highlighting its utility for assessing drought impacts on tree growth across a wide range of conditions.

The rather poor performance of the ForClim Drought Index in spring and autumn suggests that it cannot capture early- or late-season water deficits, as it is tied to the ratio of water supply to demand alone. Yet, recent studies highlight that forests in northern as well as southern Europe have experienced widespread increases in drought sensitivity in the spring to autumn periods (Jin et al., 2023), underscoring the importance of detecting such early-season droughts more effectively. To this end, an alternative approach to the ratio of supply and demand needs to be found for capturing drought intensity, because in spring and fall demand (Potential Evapotranspiration) tends to be low, but trees may still suffer if there is insufficient soil moisture to supply their needs (cf. Granier et al. 1980-2006, Breda et al. 1999).

### 4.2 A simple soil water dynamics model captures drought stress in extreme years

Model selection for simulating soil water dynamics must be made with care, as simple bucket models (Koster and Suarez, 1994) and Soil-Vegetation-Atmosphere Transfer (SVAT) models are built on fundamentally different principles. Each model type is tailored to address specific research objectives, justifying varying temporal and spatial resolutions (Ohrt et al., 2015).

On the one hand, the SVAT LWFBrook90 incorporates detailed soil hydraulic properties through the Mualem-van Genuchten parameterization, which enhances the spatial (vertical stratification) and temporal (sub-monthly) resolution of the simulated variables. However, this comes with the challenge of high data demand (cf. Table SM C2.2) and the need for extensive model calibration. This becomes a particular challenge for forest sites where meteorological data of high spatial and temporal resolution are scarce and long records of daily soil moisture data are typically absent. This can lead to high uncertainty in parameter estimation and, ultimately, model projections (Schmidt-Walter et al. 2020, Franks et al. 1997).

On the other hand, ForClim’s simple soil moisture scheme with its monthly time step and a single soil bucket layer was found to perform well for detecting strong droughts, as shown in the comparison with (1) tree-ring series at the six beech sites and (2) evapotranspiration dynamics at Lägeren. However, the simplicity of a bucket model may have limitations during wet seasons and in moist years (cf. Romano et al. 2011, Fischer et al. 2011), where ForClim tends to simulate lower water demand and higher water supply compared to LWFBrook90 (cf. Hagemann and Stacke, 2015). This behavior depends on the structural simplifications underlying bucket models that, while aimed at reducing computational complexity, tend to ignore processes such as water retention and movement across soil layers (Kondo, 1993).

Despite these differences, which are least pronounced during dry spells, both models are highly suitable for simulating the soil moisture balance, particularly in the context of longterm forest growth and development, i.e. when the focus is on seasonal patterns and inter-annual variability rather than short periods of time (e.g., days to weeks), where only LWFBrook90 would provide the required level of detail.

Overall, despite their structural and methodological differences, ForClim and LWFBrook90 feature strong agreement in simulated soil moisture depletion and declining actual evapotranspiration during dry years, in line with previous studies (e.g., Meusburger et al., 2022). Thus, although ForClim is based on a very simple framework compared to the complex SVAT approach of LWFBrook90, it is still effective in capturing strong drought events even under mesic conditions. In addition, its simplicity in terms of parameter requirements and temporal resolution render it a viable tool for studies where ease of use and low input requirements are important.

### 4.3 Predisposing and inciting factors allow to capture drought-induced tree mortality

The combination of predisposing and inciting factors (Manion, 1981) in ForClim v4.1 enabled to capture the exceptional mortality events induced by prolonged and intense droughts in Swiss temperate forests that are not usually prone to drought. Specifically, the PI framework revealed the importance of separately treating predisposing stress factors that operate over longer periods (Neycken et al. 2022), and short-term, intense stress factors that affect trees already under stress (Etzold et al., 2019). A long-term reduction in growth alone is insufficient to capture the mortality probability (Steinkamp & Hickler, 2015, DeSoto et al., 2020), and the same goes for a short-term stressor such as hydraulic failure (Körner, 2019). Our analysis suggests that a simple framework combining both aspects is suitable for predicting drought-related tree mortality, even though – or perhaps exactly because – plant-physiological processes are not modeled explicitly.

Regarding the predisposing factors, the capability to recover from stress (Schwalm et al., 2017; Ovenden et al., 2021) was captured well by a growth memory term in our PI scheme (cf. Cailleret et al., 2017; Bottero et al., 2021; Zamora-Pereira et al., 2021). By incorporating a drought-induced reduction in diameter increment over the long term, our approach closely aligns with the concept of carbon starvation (Peltier et al. 2023, Gessler et al. 2018). Our approach assesses tree vitality without engaging with the highly complex and uncertain aspects of tree carbon balance and allocation (Sala et al. 2010; Hartmann 2015; Fatichi et al. 2014). Instead, we rely on an integrative measure, i.e. tree-ring width, as a proxy of tree vitality (cf. Waring, 1980, Cherubini et al., 2021). Yet, the inclusion of tree recovery mechanisms in dynamic forest models is essential to reflect the adaptive mechanisms of trees and assess the impacts of droughts on the long-term stability of ecosystems as well as future trajectories under changing climatic conditions (Gessler, 2020).

Regarding the inciting factors, the combination of seasonal drought duration, limitations to the evapotranspiration rate along with the early- and late-seasonal soil water deficit turned out to be highly suitable for approximating the combination of low precipitation and pronounced soil moisture deficits that were found to be characteristic of the hot-dry droughts of the last decades (Breshears et al., 2005; Adams et al., 2009). Note that high Vapor Pressure Deficit (VPD), which was found to be particularly pronounced in the 2018 drought in low-elevation forests (e.g., Gharun et al., 2020) is closely correlated with a low ratio of water supply to demand, which is at the core of our formulation of the inciting factor. Soil moisture in spring and fall is critical for budburst and reserve building, respectively (Michelot et al. 2012; Massonet et al., 2021), and its integration via Relative Extractable Water (Breda et al., 1999) clearly improved the model’s capability of capturing the prolonged water deficits experienced during extreme droughts such as in 2018 (Brun et al., 2020; Gharun et al., 2020; Meusburger et al., 2022), which are projected to increase under future climatic conditions (Ruosteenoja et al., 2018).

Modeling frameworks that incorporated both carbon starvation and hydraulic failure often have failed to capture tree mortality well (e.g., Hajek et al., 2022; Fischer et al., 2024). Thus, even dynamic forest models that feature detailed physiological process representations may still lack methods, processes or feedbacks that account for multiple causes of tree mortality, thereby missing key elements of tree death (Anderegg et al. 2012, Bugmann & Seidl 2022). Our framework suggests that mortality arises from a combination of prolonged, chronic stress (which may include carbon starvation, but is not restricted to it) that is weakening trees over multiple years, followed by acute, severe stress (which may or may not include hydraulic failure) within a growing season. This integrative approach accounts for the compounding effects of long-term physiological decline and sudden environmental extremes, offering a potentially holistic depiction how trees succumb to drought. By focusing on the compound effect of chronic and acute stress, our model adds to the conventional view of a dichotomy between carbon starvation and hydraulic failure and highlights the importance of capturing stress dynamics across different scales and time frames.

The simulations for a xeric site dominated by Scots pine (site Visp) demonstrate the generality of our approach beyond the mesic beech sites. It should be noted that we did not adjust any parameters for these simulations, although the climatic setting as well as the dominant species are entirely different. To our own surprise, the simulated increase of the mortality rate under these xeric conditions was consistent with previous research (Dobbertin et al., 2004; Bigler et al., 2006; Wohlgemuth et al., 2018), suggesting that our framework may not be restricted to beech and highlighting the general importance of short, intense droughts in addition to less severe, longer-lasting droughts in the context of tree death (cf. Rigling and Cherubini, 1999).

Ultimately, our approach shows that strong model simplifications may not be a problem but a virtue, rendering it possible to assess drought-related tree mortality via the combination of growth-related and growth-independent limitations, not necessitating a physiologically based approach (Fatichi et al., 2014; Körner, 2015). Our choice reflects the trade-off between a small number of ecological assumptions while still accounting for the critical role of limiting factors, such as temperature and soil moisture, in shaping long-term tree growth responses under drought (cf. Huber et al., 2020, 2021).

### 4.4 Model behavior along a large gradient of temperature and precipitation

Since its early development (versions ≤4.0.1), ForClim featured a distinct species distribution pattern across an extended gradient’s elevational belts and climatic regions in Europe, which was attributable to the main underlying ecological assumptions (Bugmann, 1994). Our study demonstrates the continued realism of the simulations of Potential Natural Vegetation by ForClim v4.1 along the very same gradient of temperature and precipitation (Bugmann and Solomon, 2000), thus corroborating the robustness of the new formulation of drought-related mortality across sites and species. Yet, more detailed studies are required to further substantiate this, using data e.g. from the ICP Level-I and Level-II network (Bussotti et al. 2024) regarding widespread, recently drought-inflicted species such as beech (*Fagus sylvatica*), spruce (*Picea abies*) or Scots pine (*Pinus sylvestris*), to generalize from our case studies to larger areas (Knapp et al. 2024).

The simulations with the new model version (v4.1) along this gradient featured not only a reduction of total basal area to more plausible values (cf. Idoate et al., 2024), but also distinct differences at the level of the abundance of individual species. The reduction in the abundance of spruce (*Picea abies*) in low-elevation areas is consistent with descriptions by Frehner et al. (2005), and the same goes for the decrease of beech (*Fagus sylvatica*) towards drier sites (Bohn & Gollub, 2006). Yet, in ForClim v4.1 beech continued to dominate forests on the Swiss Plateau and strongly suppressed other species via light competition, which is also in line with previous studies (cf. Heiri et al. 2009). Towards the dry end of the gradient, beech lost dominance in favor of lime (*Tilia* spp.) and Scots pine (*Pinus sylvestris*), which is confirmed by studies e.g. from the northern German lowlands (Diers et al., 2023) as well as general vegetation mapping efforts (Bohn & Gollub, 2006). Thus, the PI scheme that we introduced did not only improve model behavior under drought conditions, but it generally improved the depiction of competitive relationships along this very wide environmental gradient.

### 4.5 Limitations and outlook

The quantitative testing of our PI framework relied on data from a small set of temperate, mesic beech-dominated sites and one Scots-pine dominated stand, which may limit the general applicability of our findings. Additionally, the absence of comprehensive inventory data and long-term observations required us to reconstruct stand structure partly based on circumstantial evidence, which may have reduced the accuracy of our results. While drought events like the one in 2018 seemed rare or even erratic at the time of observation, their repeated occurrence over the last years highlights the need for long-term monitoring, including annual, tree-level mortality assessments, to better capture the ongoing effects of these extreme events.

To simulate Potential Natural Vegetation, we decided to use a gradient of sites across a wide range of climatic conditions in Switzerland and Germany. This was primarily a modeling choice rather than one rooted in the reality of the landscape, which strongly limits the quantitative comparison of simulated vs. ‘real’ data, as there are few if any primeval forests in Europe that could be considered to be in equilibrium with climate. A way forward would be to validate the model against data from strict forest reserves (cf. Käber et al. 2024 for tree regeneration), thus providing a more tangible baseline for quantitative comparisons (cf. Brang and Bolliger, 2015).

A clear limitation of our study concerns soil water dynamics and the comparison of the ForClim and LWFBrook90 models, which was difficult due to the scarcity of long-term data on key components of the water balance in forests beyond precipitation, i.e. soil moisture, evapotranspiration, and runoff. The analyses of simulated soil water dynamics would benefit from an in-depth data-model comparison using long-term time series data from multiple sites.

Lastly, the lack of measured soil data (i.e., quantitative analyses of samples from soil profiles) introduced some degree of uncertainty in the simulated soil moisture dynamics as well as the simulated forest responses to drought. As soil properties vary tremendously in space, particularly in forest stands, high-resolution soil maps are key tools to derive estimates of AWC (cf. Baltensweiler et al. 2022). These products, however, are characterized by substantial uncertainty, particularly when it comes to soil depth and rock fraction, which play a major role in determining AWC. Constraining this uncertainty and enhancing the quality of soil maps is key to better constrain and assess the impacts of drought stress on forests (Walthert et al. 2020, Klesse et al., 2022).

## 5. Conclusions

We demonstrated that widely applied drought indices like the SPI or SPEI, although showing significant responses in the summer months, are overall not capable of detecting drought signals at mesic sites across multiple seasons, even though these droughts have led to tree mortality. By contrast, indices of higher complexity, particularly the ForClim drought index, are suitable for identifying such dry years and capturing their effect on tree growth and demography. We therefore recommend using indices such as the ForClim drought index; the essential feature is that such indices must consider soil water storage in addition to mere climatic data.

Our comparison of the highly detailed LWFBrook90 model with the simple ForClim soil moisture balance model, which is based on a monthly resolution, suggests that annual drought severity can be captured well with a highly simplified scheme, although it is clear that for capturing any higher-resolution features in either time or space, particularly under milder drought conditions, SVAT models may be preferred. In the context of drought effects on growth and mortality of forest trees, we posit that simple soil moisture balance models are sufficient, bringing the distinct advantage of ease of parameterization and low runtime.

The novel framework for drought-related tree mortality that we developed suggests that the combination of predisposing and inciting factors is pivotal. It furthermore indicates that trees do not normally die due to carbon starvation or hydraulic failure alone, but due to the combination of prolonged stress with sudden (within a year) severe stress that covers the entire growing period. We even posit that it is possible to model these ecophysiological processes in a highly simplified way, specifically that they can be formulated at an aggregate, general level. Even if simplified, our framework permits to (1) isolate the multiple factors as drivers of tree responses to drought stress and (2) accurately simulate beech mortality in response to the 2018 drought, even at sites not usually prone to drought-related mortality. The new mortality model also features high robustness when applied to a xeric system with another species (Scots pine) as well as along an extended climatic gradient and mixed species stands.

## Acknowledgements

We are thankful to Dr. Anna Neycken and Dr. Mathieu Lévesque (Chair of Silviculture, Dept. Environmental Systems Sciences, ETH Zürich) for providing the inventory and tree-ring data of the six beech sites. We thank Prof. Dr. Christof Bigler (Chair of Forest Ecology, Dept. Environmental Systems Sciences, ETH Zürich) for the discussions on detrending predictors and response variables altogether as good scientific practice in dendrochronology. We are thankful to Dr. Fabian Bernard (Oeschger Centre for Climate Change Research, University of Bern; WSL, Birmensdorf) for advice on the selection of LWFBrook90 outputs and how to aggregate them, and to Dr. Lorenz Walthert (WSL, Birmensdorf) for references on the relationship between soil water and drought stress. The ForClim source code was curated by Hussain Abbas and Dr. Thomas Oliver Hands (Forest Ecology, ETH Zurich), and they deserve a special acknowledgment for their excellent support.

Gina Marano was supported by the project entitled “Embracing structural uncertainty in models of forest dynamics”, under Grant Agreement N. 188882 which has received funding by the Swiss National Science Foundation (SNSF).

## Credit authorship contribution statement

Gina Marano, Harald Bugmann and Ulrike Hiltner conceived the idea, developed the concept and framing of the paper; Gina Marano conducted the analyses, created the graphics, and drafted the main manuscript; Katrin Meusburger supported the work with LWFBrook90 simulations, input data and provided advice regarding the analyses; Gina Marano and Thomas Oliver Hands curated the ForClim source code. All authors reviewed the manuscript and approved its submission.

## Data availability

All R scripts, R packages and codes are archived in a Zenodo repository and can be accessed at: https://zenodo.org/records/14234972

## Supplementary Material

### SM A: Climatic anomalies and site inventories

**Figure A1:**
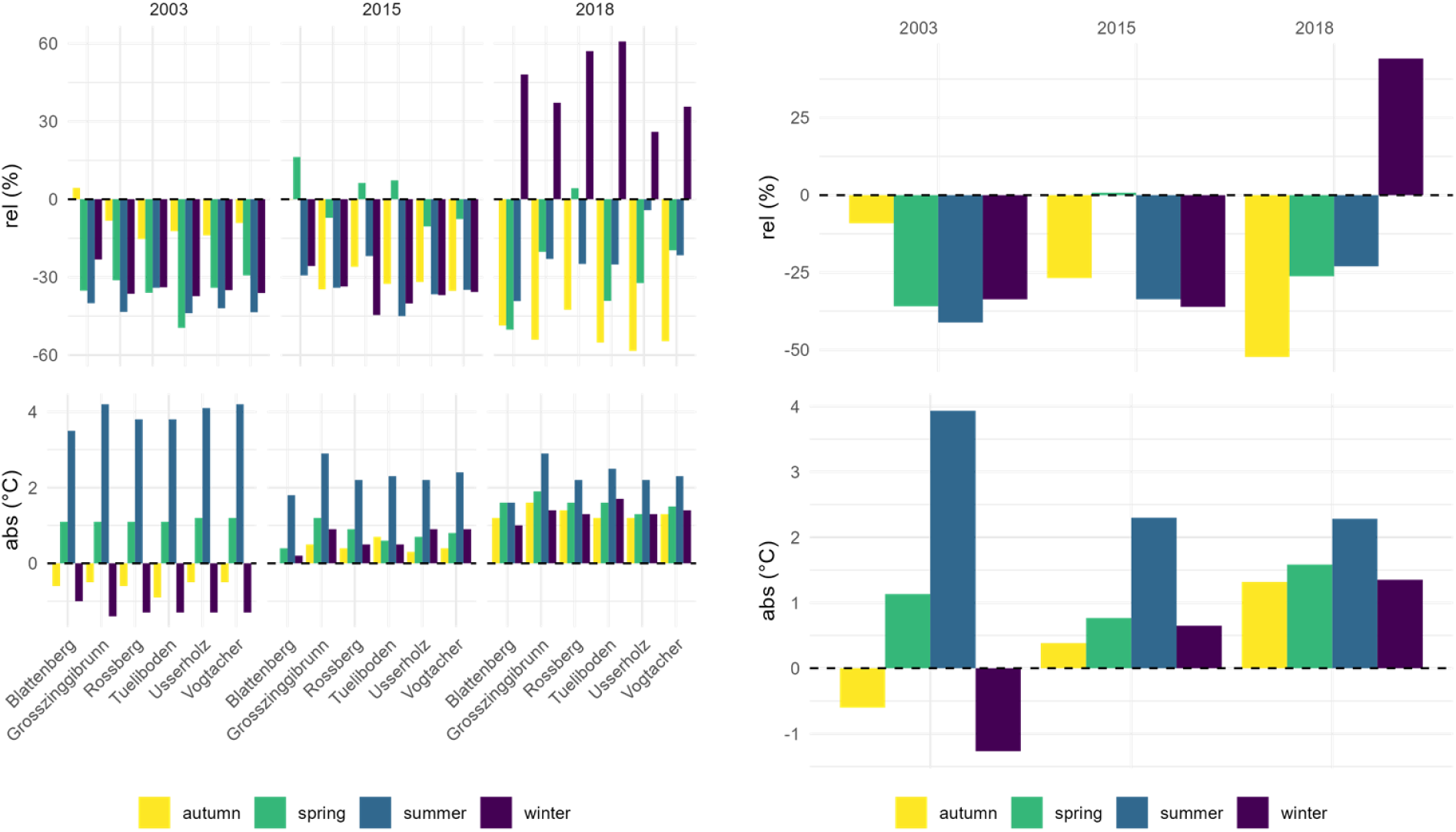
Precipitation and temperature anomalies for the years 2003, 2015 and 2018 relative to the reference period 1980-2010 across sites (left panel) and averaged across the six beech sites (right panel).

**Table A1:** Measured basal area (m^2^ ha^-1^) at the six beech sites from 2015 to 2020, as retrieved from Neycken et al. (2022).

| *Site* | *Time* | *Basal area* |
| --- | --- | --- |
| Usserholz | 2016 | $25.8 \pm 1.7$ |
| | 2018 | $20.0 \pm 5.6$ |
| | 2019 | $20.2 \pm 5.5$ |
| | 2020 | $19.9 \pm 5.6$ |
| Vogtacher | 2016 | $23.0 \pm 1.4$ |
| | 2018 | $15.0 \pm 4.6$ |
| | 2019 | $14.2 \pm 4.2$ |
| | 2020 | $14.5 \pm 4.4$ |
| Tüeliboden | 2015 | $31.5 \pm 1.2$ |
| | 2018 | $20.7 \pm 6.4$ |
| | 2019 | $20.9 \pm 7.8$ |
| | 2020 | $19.9 \pm 7.4$ |
| Rossberg | 2015 | $28.1 \pm 1.0$ |
| | 2018 | $25.0 \pm 9.2$ |
| | 2019 | $25.0 \pm 8.7$ |
| | 2020 | $24.3 \pm 9.6$ |
| Grosszinggibrunn | 2016 | $19.7 \pm 1.5$ |
| | 2018 | $12.4 \pm 4.6$ |
| | 2019 | $11.6 \pm 4.1$ |
| | 2020 | $12.0 \pm 3.6$ |
| Blattenberg | 2017 | $33.0 \pm 1.0$ |
| | 2018 | $27.7 \pm 6.0$ |
| | 2019 | $28.6 \pm 6.2$ |
| | 2020 | $26.5 \pm 5.7$ |

### SM B: Tree-ring data and drought indices

#### B1: Ring-width indices

**Fig B1.1:**
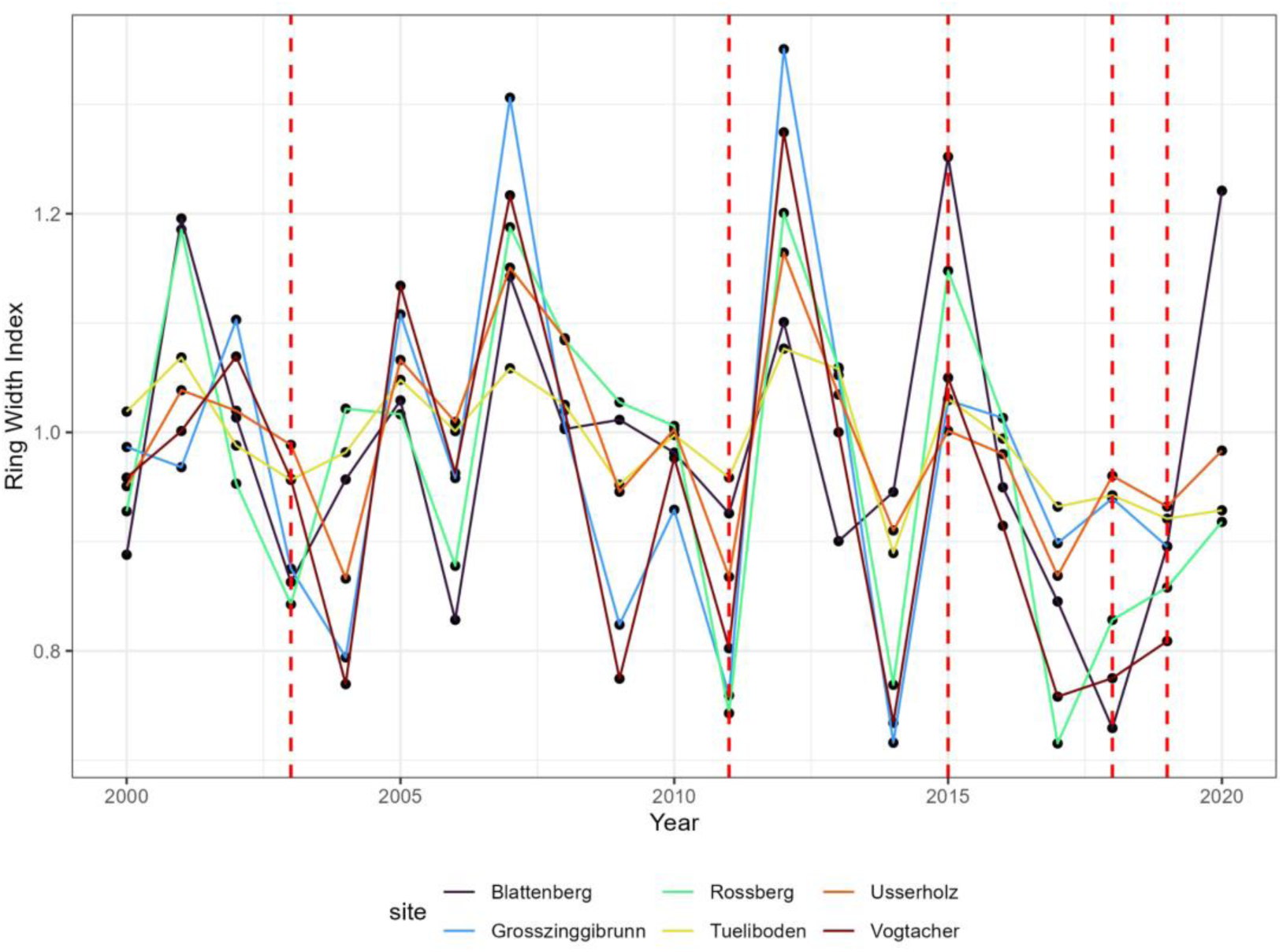
Ring-width Index (RWI) standard and residual chronologies of the six sites. Red dashed lines indicate the drought years 2003, 2011, 2015, 2018-2019.

**Fig B1.2:**
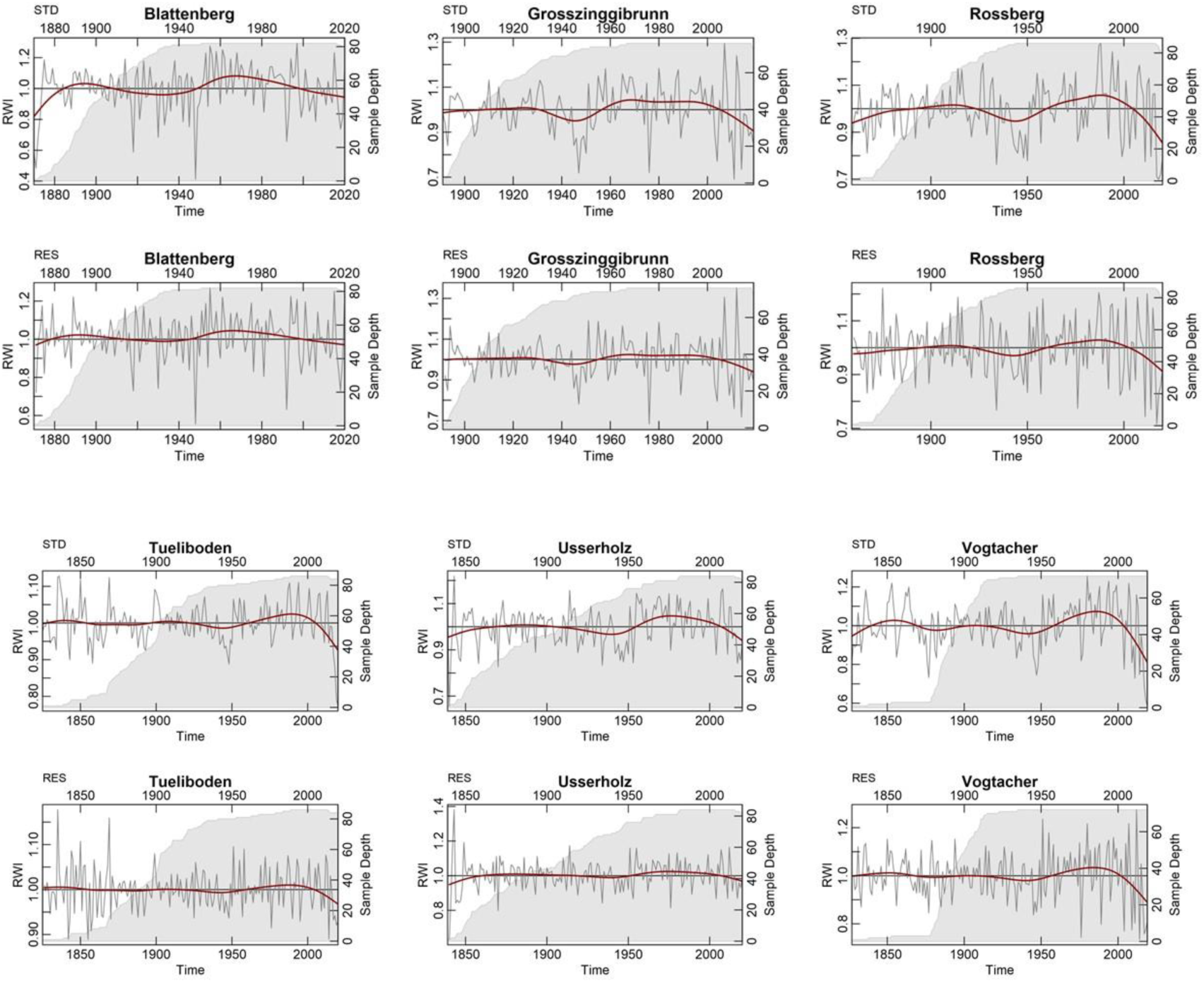
Tree-ring chronologies for the six beech sites. RES stands for residual chronology, which was obtained using auto-regressive models (AR) of the individual tree series, while STD indicates the standard chronology. The shaded area represents the sample depth. The main spline is indicated by the red line (i.e., low-pass filter).

#### B2: Z-chronologies for the identification of pointer years

**Fig B2:**
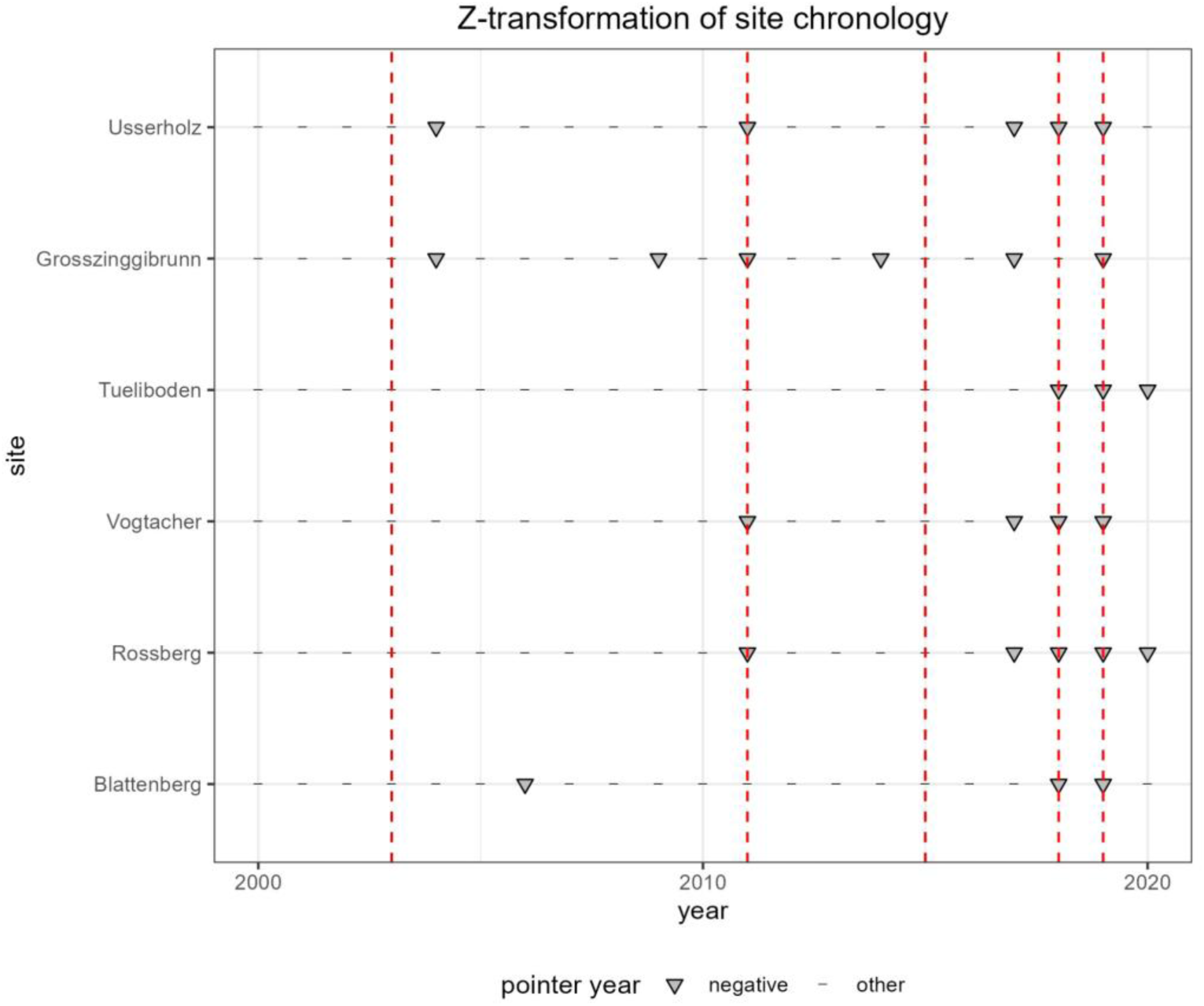
Calculation of pointer years for the six sites using a z-transformation of the site chronologies (Tukey’s biweight robust mean site chronology). The dashed lines indicated dry years 2003, 2011, 2015, 2018 and 2019.

#### B3: Tree-rings indices and drought indices

**Table B3.1:** Mann-Kendall seasonal test for the five drought indices at the six beech sites.

| *Site* | *p-value* | $\tau$ | *Drought Index* | *significant* |
| --- | --- | --- | --- | --- |
| *Blattenberg* | 0.073690743 | -0.053947096 | CWB | FALSE |
| *Grosszinggibrunn* | 0.218507857 | -0.037115382 | CWB | FALSE |
| *Rossberg* | 0.464476155 | -0.022063907 | CWB | FALSE |
| *Tüeliboden* | 0.105823985 | -0.048780892 | CWB | FALSE |
| *Usserholz* | 0.156277559 | -0.042762109 | CWB | FALSE |
| *Vogtacher* | 0.313238929 | -0.030417431 | CWB | FALSE |
| *Blattenberg* | 0.215672487 | -0.037384032 | SPI | FALSE |
| *Grosszinggibrunn* | 0.527478963 | -0.019058972 | SPI | FALSE |
| *Rossberg* | 0.8687562 | -0.004984186 | SPI | FALSE |
| *Tüeliboden* | 0.180862792 | -0.040362473 | SPI | FALSE |
| *Usserholz* | 0.223478476 | -0.036718589 | SPI | FALSE |
| *Vogtacher* | 0.532513542 | -0.018827465 | SPI | FALSE |
| *Blattenberg* | 0.062978216 | -0.056083123 | SPEI | FALSE |
| *Grosszinggibrunn* | 0.06721257 | -0.05520507 | SPEI | FALSE |
| *Rossberg* | 0.308520949 | -0.030715478 | SPEI | FALSE |
| *Tüeliboden* | 0.053881759 | -0.063989469 | SPEI | FALSE |
| *Usserholz* | 0.040411808 | -0.059270359 | SPEI | FALSE |
| *Vogtacher* | 0.117173918 | -0.047257129 | SPEI | FALSE |
| *Blattenberg* | 0 | -0.348273502 | sc-PDSI | TRUE |
| *Grosszinggibrunn* | 1.97373E-09 | -0.183332548 | sc-PDSI | TRUE |
| *Rossberg* | 6.24286E-06 | -0.138075743 | sc-PDSI | TRUE |
| *Tüeliboden* | 3.21965E-14 | -0.231929032 | sc-PDSI | TRUE |
| *Usserholz* | 1.3468E-10 | -0.19622966 | sc-PDSI | TRUE |
| *Vogtacher* | 1.91588E-08 | -0.171724536 | sc-PDSI | TRUE |
| *Blattenberg* | 0.271524187 | 0.039440173 | ForClim | FALSE |
| *Grosszinggibrunn* | 0.251986335 | 0.041176087 | ForClim | FALSE |
| *Rossberg* | 0.526524962 | 0.022411501 | ForClim | FALSE |
| *Tüeliboden* | 0.313488755 | 0.036149312 | ForClim | FALSE |
| *Usserholz* | 0.828500556 | -0.007769694 | ForClim | FALSE |
| *Vogtacher* | 0.354662213 | 0.03323285 | ForClim | FALSE |

**Table B3.2:** Mann-Kendall test on the detrended sc-PDSI index by applying the STL method

| *Site* | *p-value* | $\tau$ | *Drought Index* | *significant* |
| --- | --- | --- | --- | --- |
| *Blattenberg* | 0.058372779 | -0.059543321 | sc-PDSI | FALSE |
| Grosszinggibrunn | 0.88910471 | -0.004205785 | sc-PDSI | FALSE |
| Rossberg | 0.119632891 | -0.046942523 | sc-PDSI | FALSE |
| Tüeliboden | 0.543383497 | 0.018329938 | sc-PDSI | FALSE |
| Usserholz | 0.548127808 | -0.018114681 | sc-PDSI | FALSE |
| Vogtacher | 0.686588218 | -0.012170284 | sc-PDSI | FALSE |

**Figure B3:**
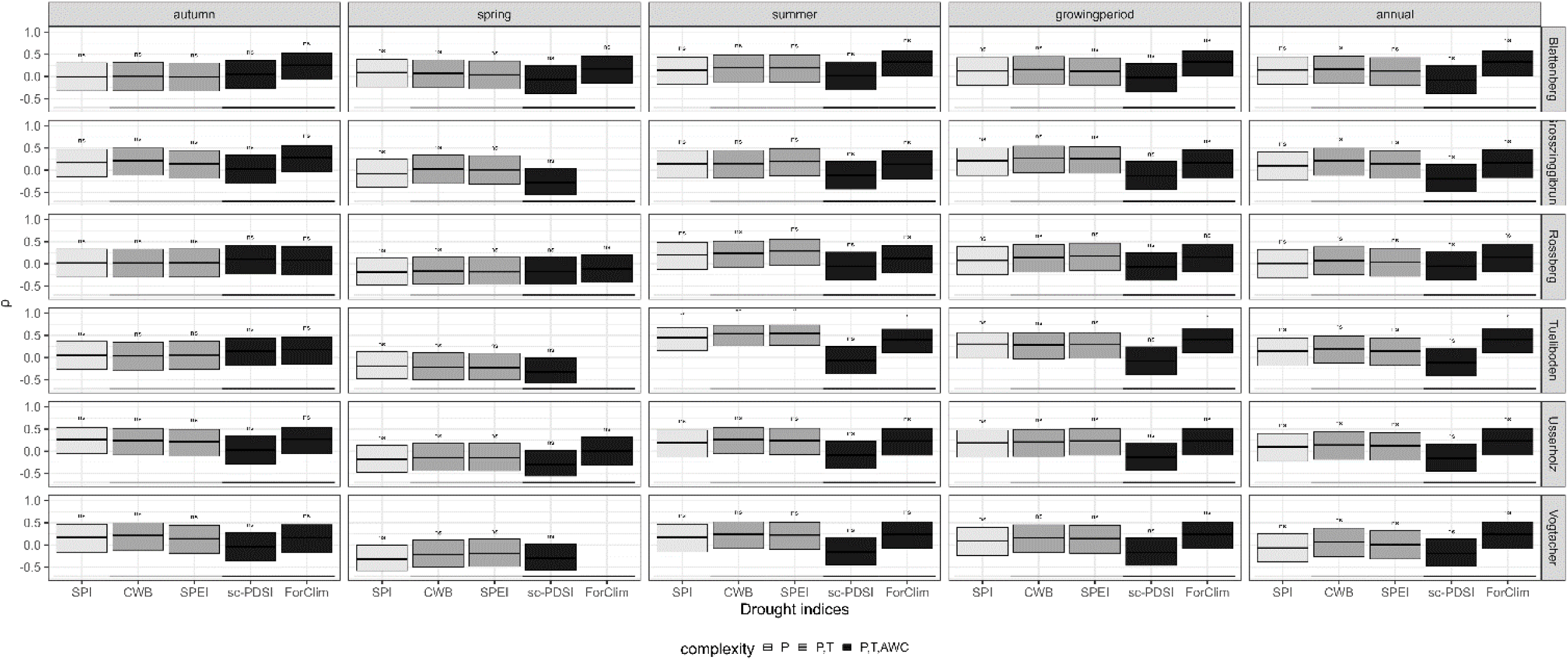
Seasonal non-parametric correlations between tree-ring width indices and drought indices of previous year (lag=1).

### SM C: LWFBrook90 and ForClim model comparison

#### C1: Lägeren simulated and observed soil moisture

**Fig C1:**
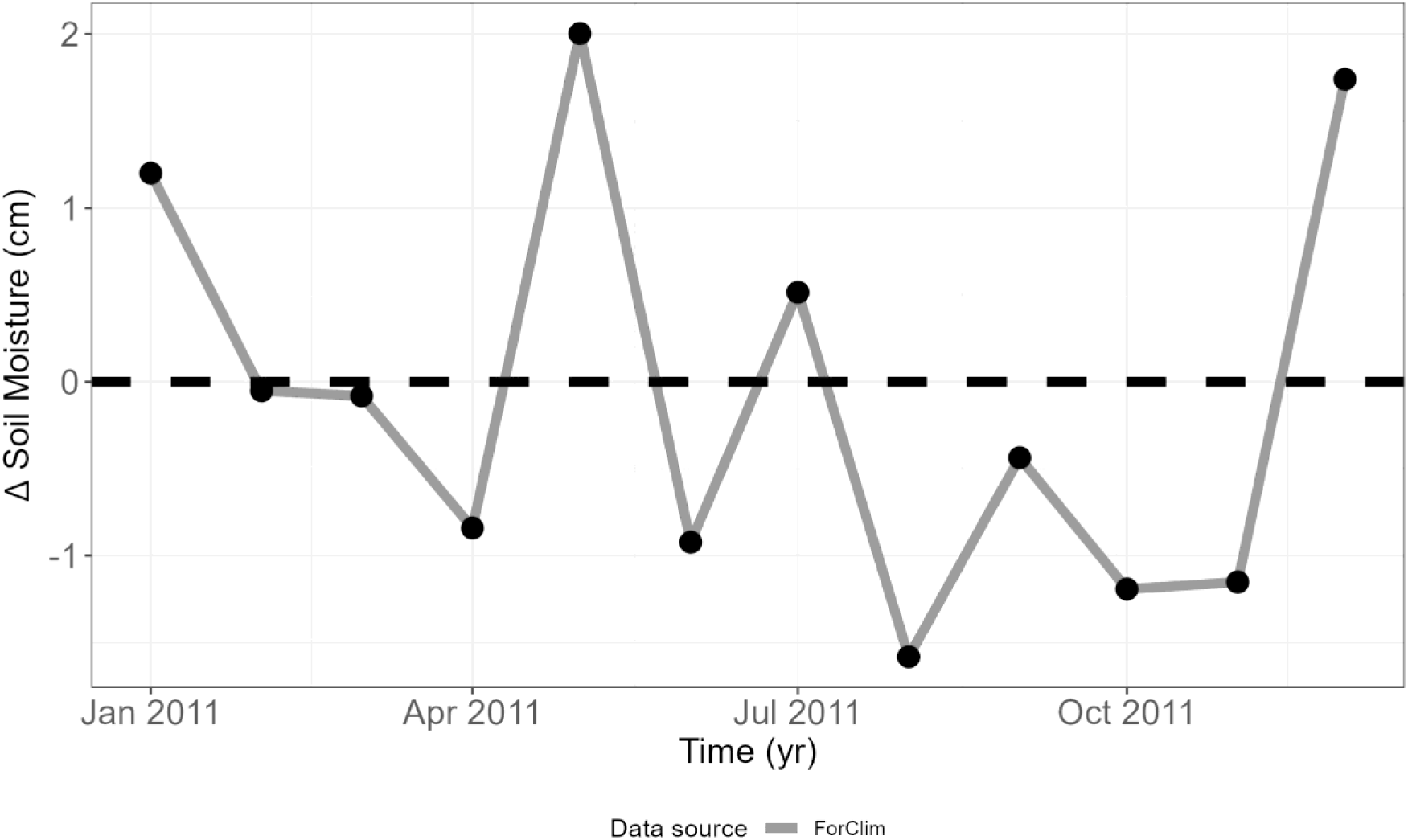
Difference between observed and simulated (ForClim) soil moisture in the year 2011. The dashed black line indicates the reference (observed data) indicating no difference between observed and simulated soil moisture. Observed data were retrieved from Pastorello, G., Trotta, C., Canfora, E. *et al*. The FLUXNET2015 dataset and the ONEFlux processing pipeline for eddy covariance data. *Sci Data* **7,** 225 (2020). https://doi.org/10.1038/s41597-020-0534-3

#### C2: Simulation setup for LWFBrook90

**Table C2.1.** Parameter set LWFBrook90 for the Lägeren. Sid: site id, Sp: species, El: elevation (m a.s.l), Sl: slope, As: aspect, Bas: basal area (m^2^ ha^-1^), Ag: age (years), He: height (m), Lm: maximum LAI (m^2^ m^-2^).

| *SId* | *Lat* | *Lon* | *Sp* | *El* | *Sl* | *As* | *Bas* | *Ag* | *He* | *Lm* |
| --- | --- | --- | --- | --- | --- | --- | --- | --- | --- | --- |
| GZ | 47.51 | 7.67 | Fagus sylvatica | 407.3 | 0.26 | 0.37 | 11.64 | 99 | 22.3 | 5 |
| VA | 47.51 | 7.7 | Fagus sylvatica | 467.2 | 0.24 | 6.03 | 14.22 | 119 | 24.7 | 5 |
| TB | 47.54 | 8.21 | Fagus sylvatica | 468.3 | 0.28 | 2.67 | 19.9 | 128 | 23.8 | 5 |
| UH | 47.48 | 7.48 | Fagus sylvatica | 570.8 | 0.12 | 2.72 | 19.91 | 132 | 22 | 6 |
| BB | 47.31 | 9.55 | Fagus sylvatica | 539.49 | 0.39 | 1.48 | 26.57 | 122 | 24.3 | 5 |
| RB | 47.67 | 8.57 | Fagus sylvatica | 515.1 | 0.09 | 2.55 | 24.29 | 122 | 24.6 | 5 |

**Table C2.2:**
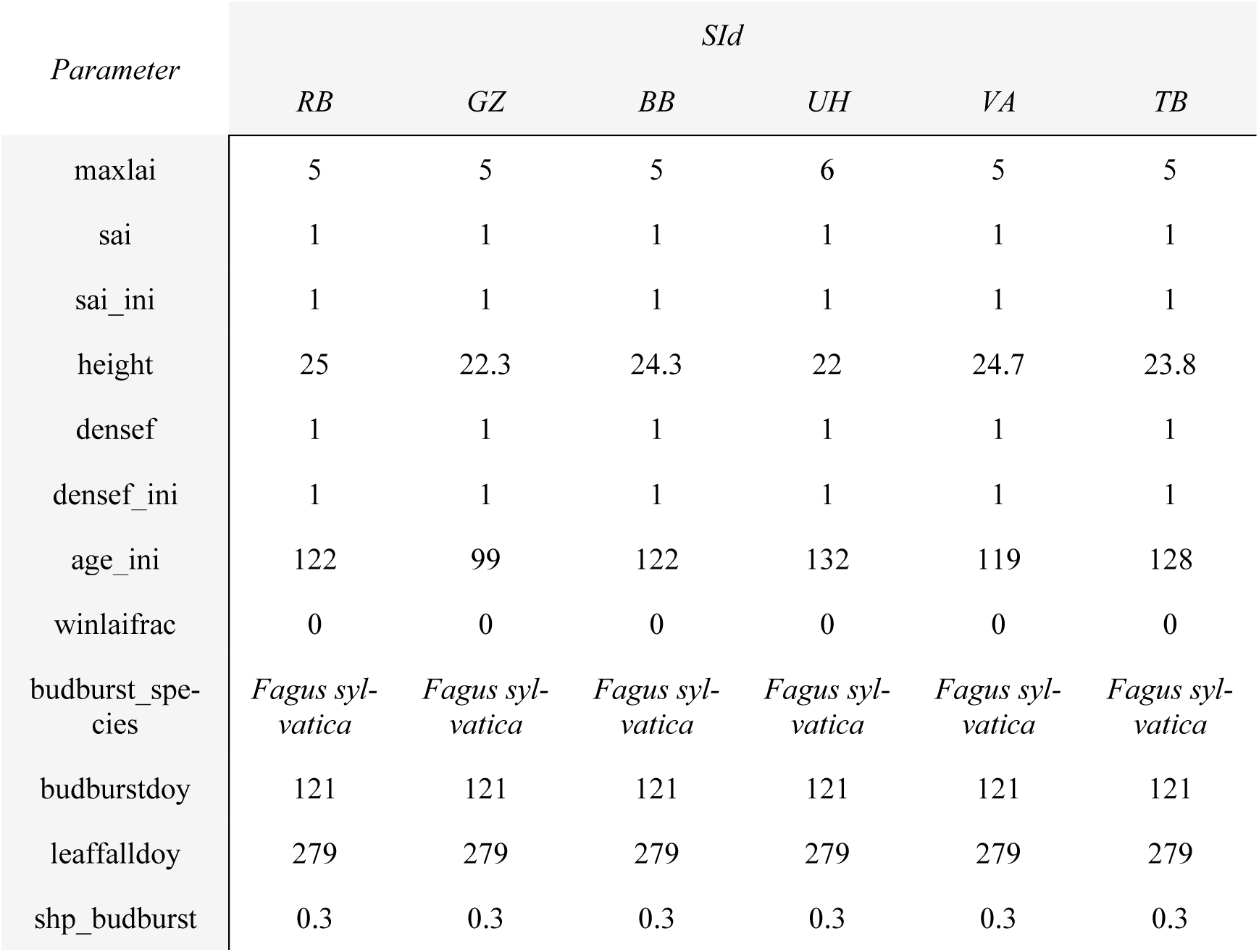
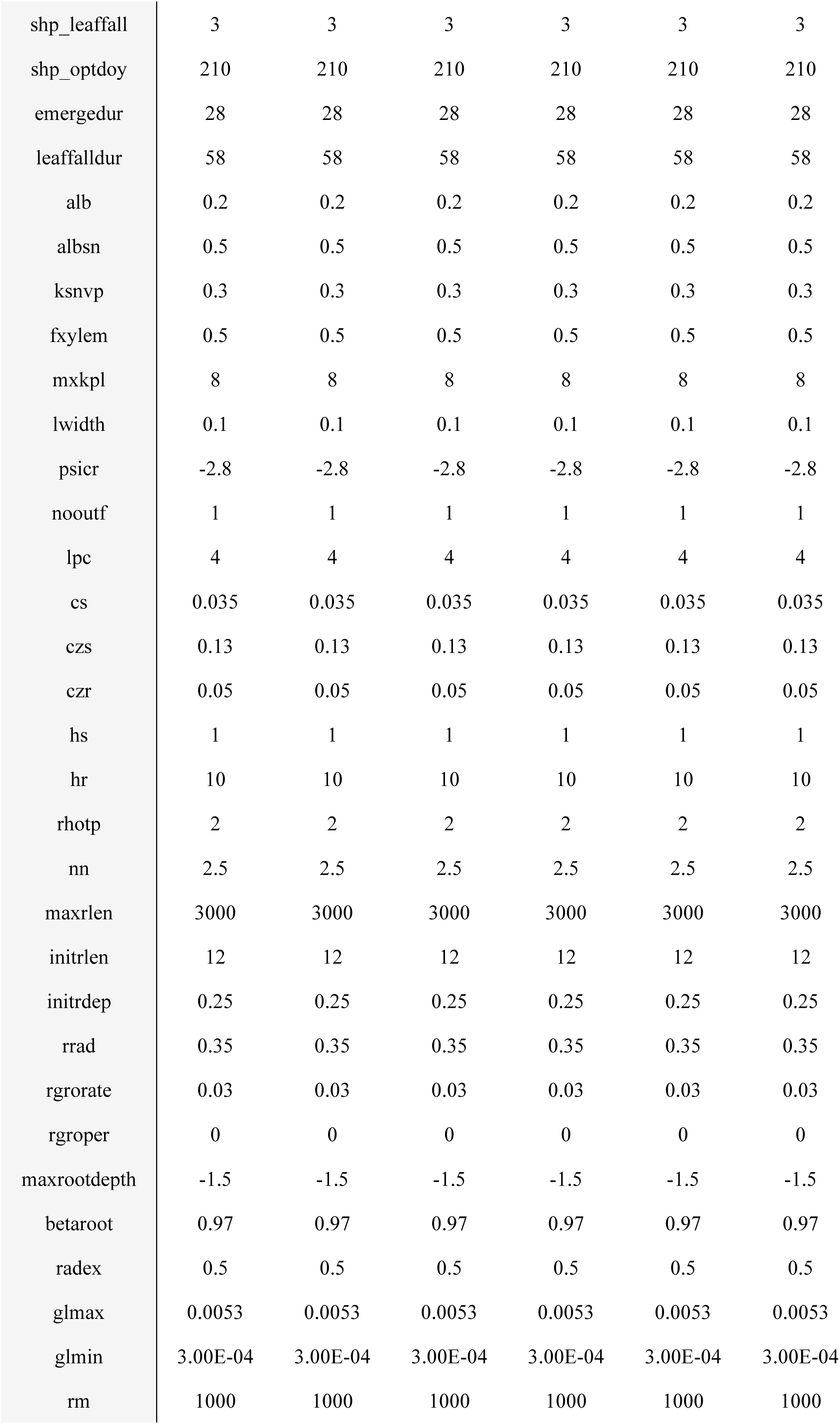
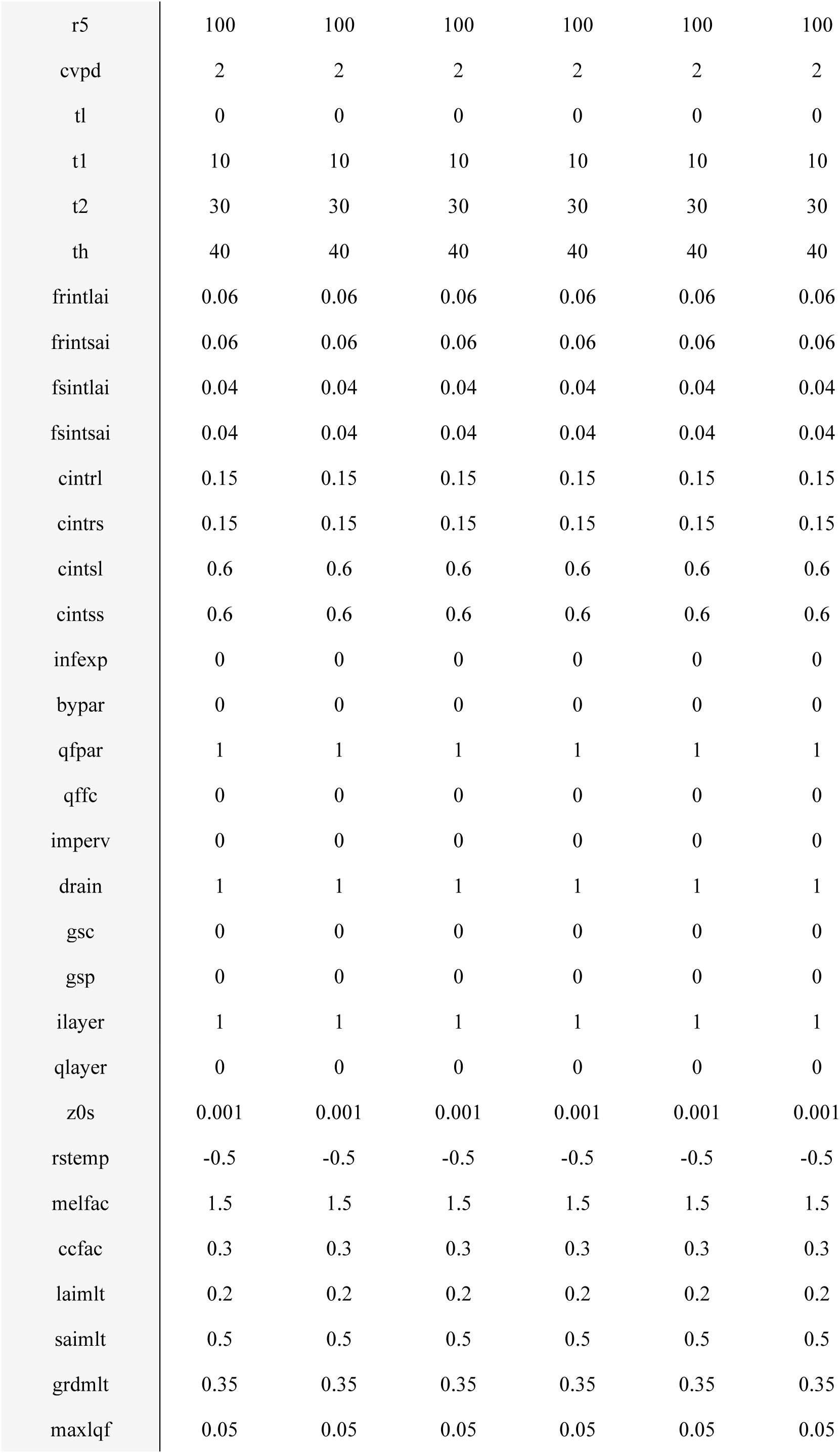
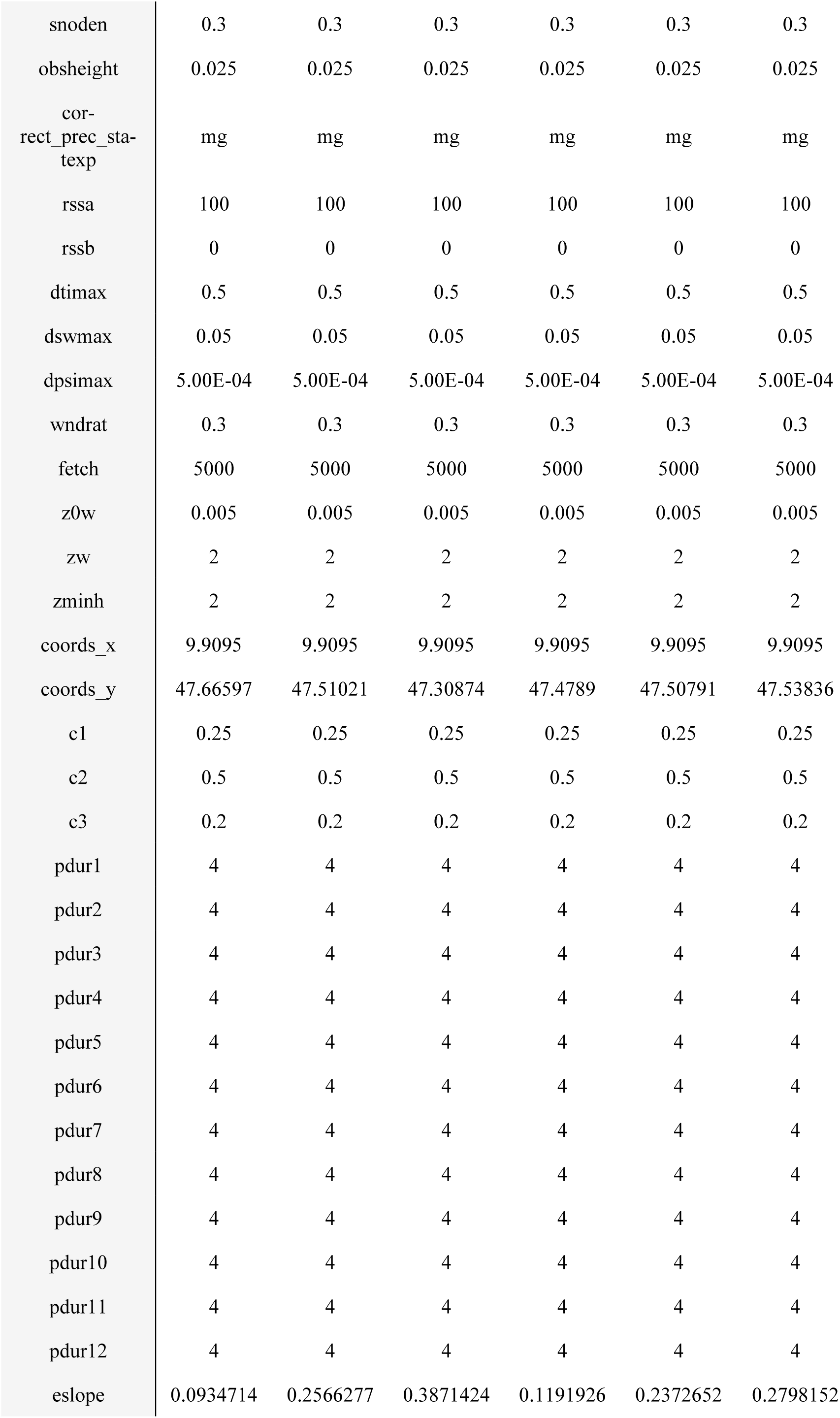
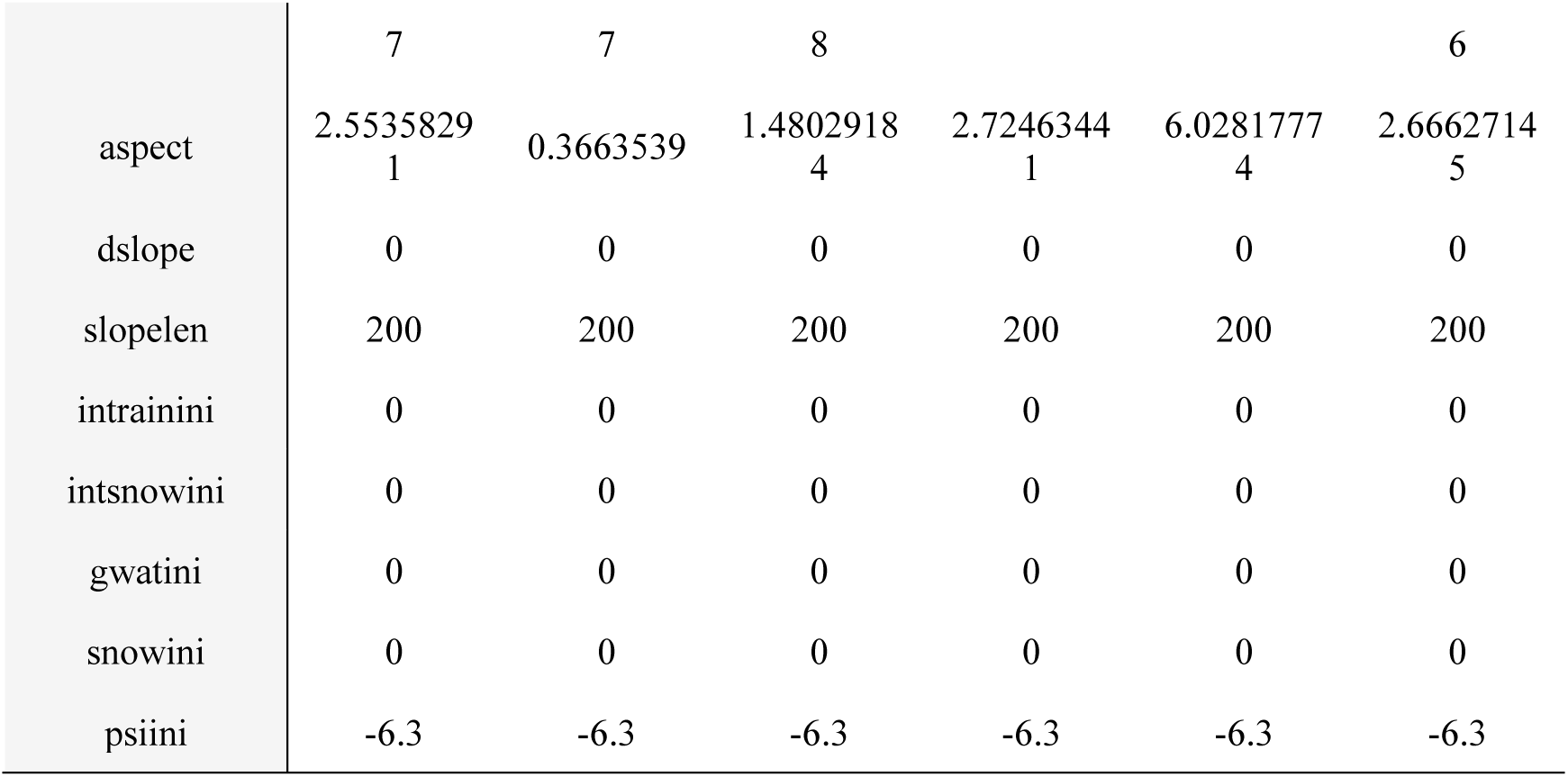
Parameter set for LWFBrook90 simulations for the six beech-dominated sites (site id, SId). For a detailed description of each parameter consult Schmidt-Walter et al. (2020).

**Fig C2.1:**
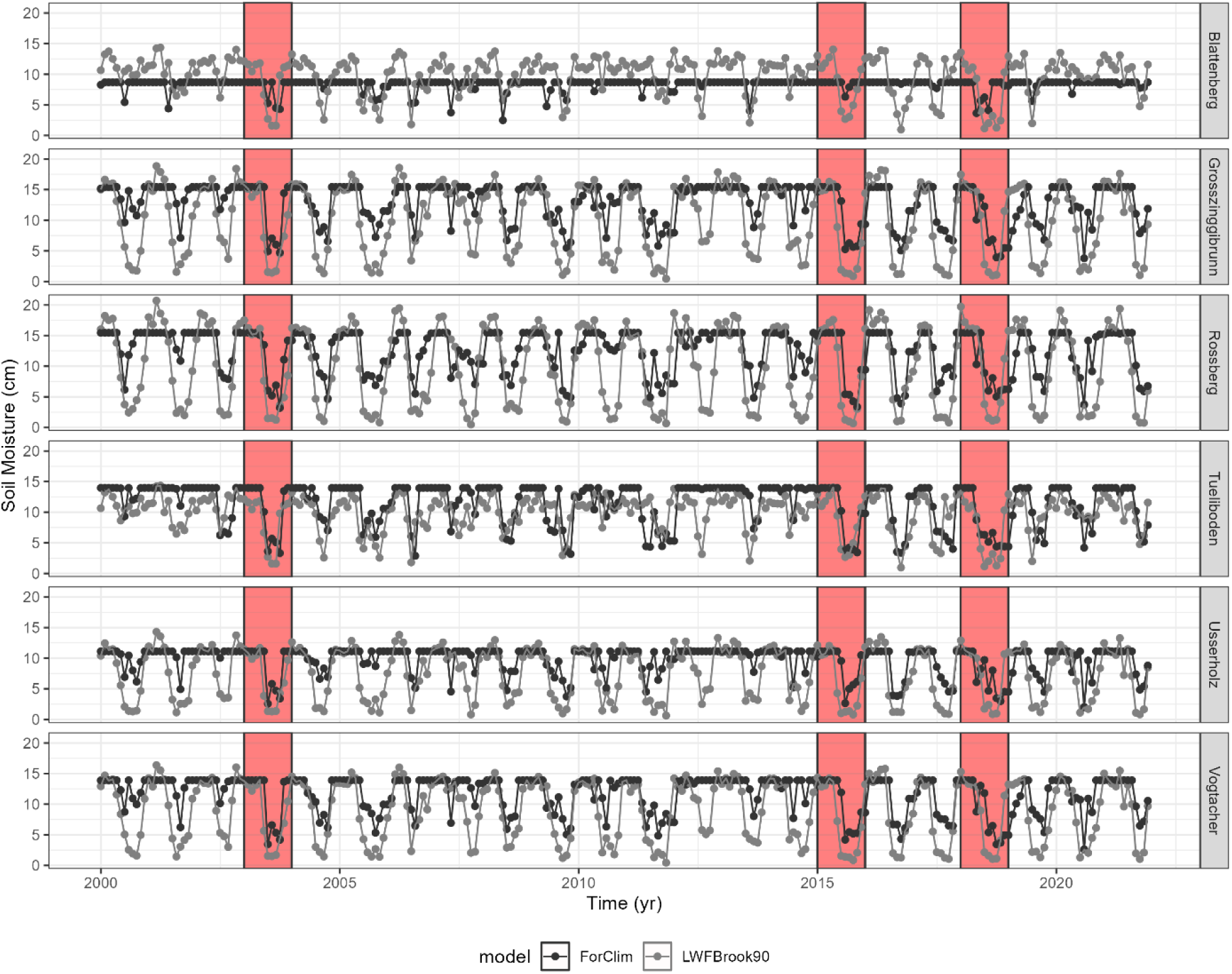
Simulated monthly soil moisture at the six beech sites. Drought years 2003, 2015 and 2018 are highlighted in red.

**Fig C2.2:**
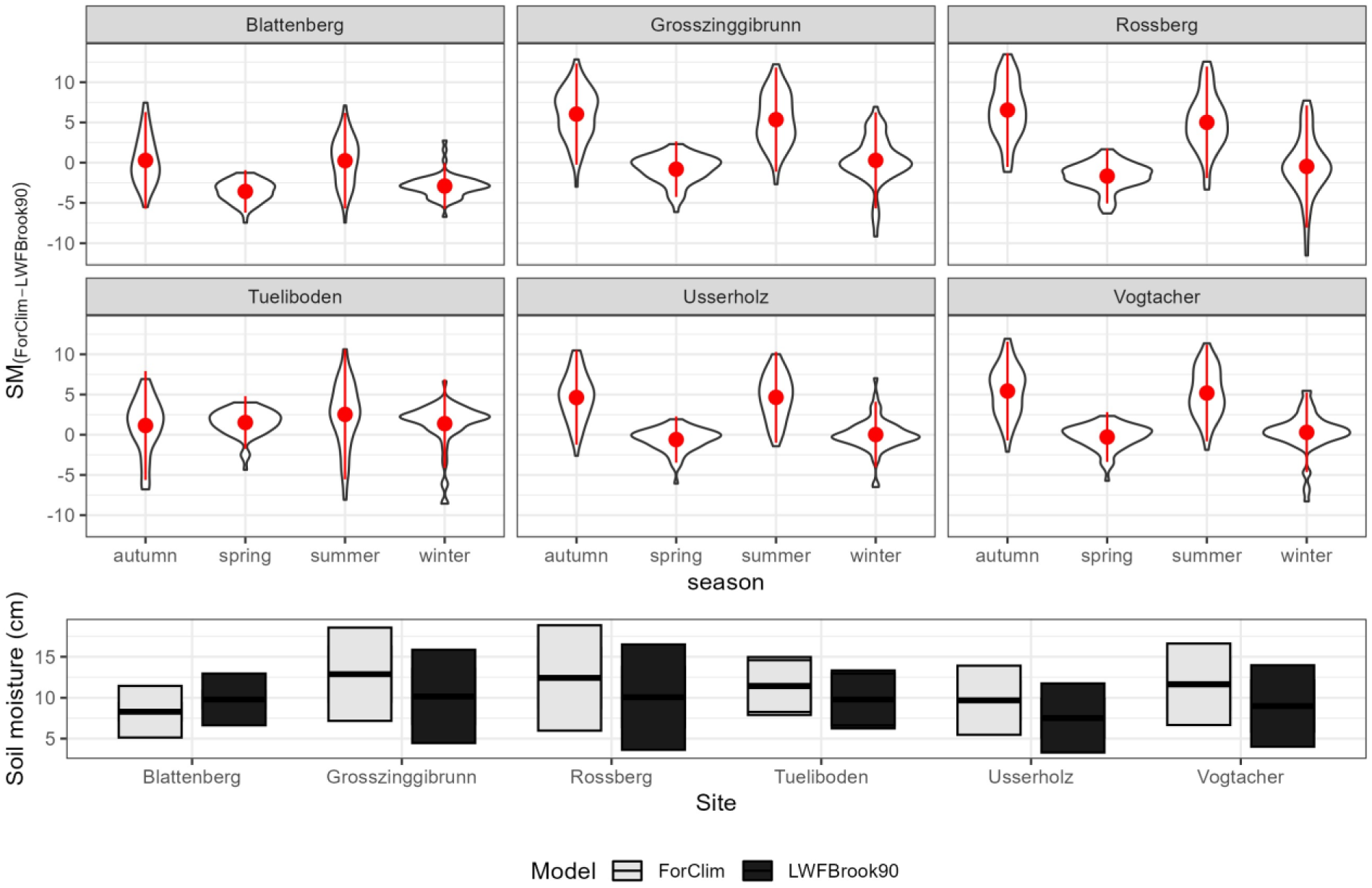
Soil moisture differences between ForClim and LWFBrook90 across seasons (upper panel) and yearly means (lower panel) at the six beech sites.

**Fig C2.3:**
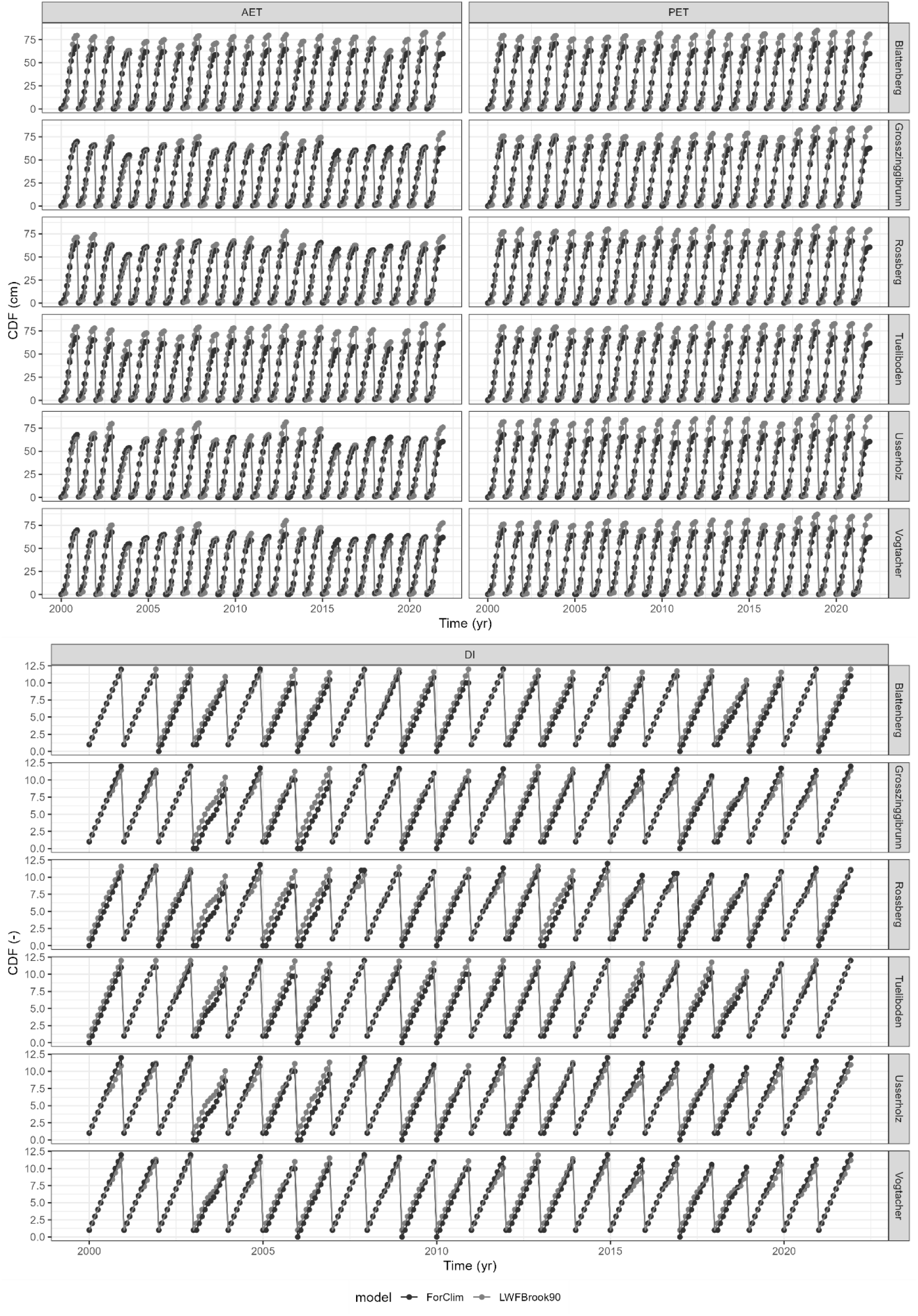
Annual cumulative AET, PET and DI (AET/PET) from the two models at the six beech sites.

**Figure C2.4:**
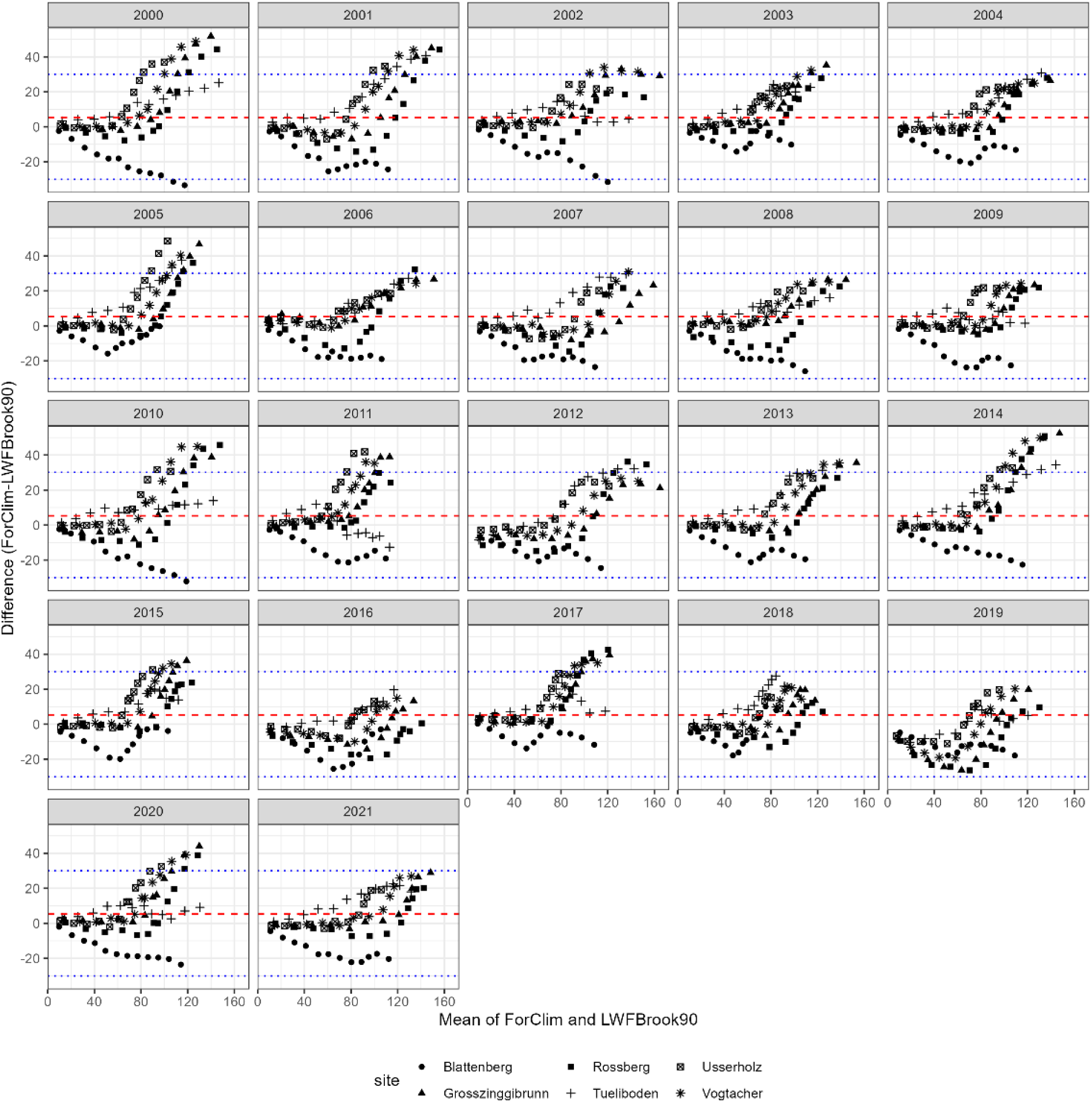
Bland-Altmann plots of yearly cumulated soil moisture (CSM) across the six beech sites of the two models for the for the distinct years of the simulation period.

### SM D: Description of new features in ForClim 4.1

#### D1 Distributed soil properties per patch (kBS distribution)

Soils feature high spatial variability in forests as the result of the interplay of geology, geomorphology, vegetation, and time (Binkley & Fisher, 2013; Boyle, 2005; Osman, 2013). Notably, one of the most influential factors in this context is soil depth, determined by weathering and affecting nutrient availability. Within this mosaic of soil properties, hydraulic properties pose a notable source of uncertainty, inherently linked to physical attributes. This uncertainty can be interpreted as being stochastic because there is no soil data at meter resolution across entire forest stands. Understanding and quantifying this apparent stochasticity is pivotal for accurately characterizing soil properties, with implications for ecosystem functioning, hydrological processes, and sustainable land management practices in forested environments.

To accurately capture the influence of drought within specific time frames, such as the drought events of 2003, 2015, and 2018, time series of precipitation and temperature need to be used. Such time series provide a detailed account of weather conditions, allowing us to accurately model and analyze the drought signals provided that we can capture the spatial variability of soil physical properties. To achieve this, we attributed distinct soil properties, particularly soil water availability, to each forest patch of ForClim, thereby mimicking the heterogeneity present in natural ecosystems. This approach ensured that the simulation experiments appropriately reflected the intricate interplay between drought, weather patterns, and spatial heterogeneity in soil properties across forest stands.

The lognormal probability distribution is often used to model skewed and left-truncated variables (such as the “bucket size” of ForClim). It was in this case selected to attribute to each forest patch its own soil properties.

In the ForClim site file, values for kBS_mean_ and kBS_min_ must be provided, and the standard deviation (*σ*) and mean (*μ*) of the lognormal distribution are calculated using these values. The standard deviation (*σ*) is determined as:

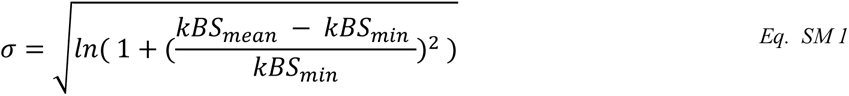

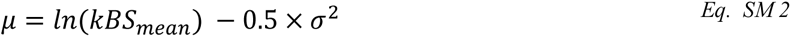

A loop is initiated to sample *kBS* values from the lognormal distribution until a valid value greater than or equal to the minimum is obtained. The sampling is performed using the Box-Muller transform to generate a lognormally distributed random variable based on the calculated *μ* and *σ*.

As the water balance module in ForClim did not undergo changes compared to version 4.0.1 (Huber et al., 2020), the mathematical formulation of E (transpiration), D (water demand), PET (potential evapotranspitation), AET (actual evapotranspiration) and SM (soil moisture) are not reported here but can be found in the ForClim documentation (cf. Data Availability).

#### D2 Parameter derivation: pedotransfer functions

The determination of kBS_mean_ and kBS_min_ is based on soil textural properties (sand, silt, clay), along with additional parameters such as upper and lower boundaries, thickness, and gravel content per genetic horizon. From these data, the available water capacity is determined by means of pedotransfer functions (Schmidt-Walter et al., 2020).

Soil textural properties were derived from the 25 m resolution product for Swiss soils provided by Baltensweiler et al. (2021) . Three pedotransfer functions were tested: ROSETTA (Zhang & Schaap, 2017), Wessolek (2009) and Puhlmann & von Wilpert (2012).

In the ROSETTA model, van Genuchten (VG) model curves are calculated, and the available water capacity (AWC) is retrieved based on matric potentials. The Wessolek method is based on field capacity, wilting point,and available water content using a pre-defined van Genuchten function. The third method, similar to Wessolek, calculates field capacity, wilting point, and available water content using a pre-defined van Genuchten function, yet it requires bulk density and soil organic carbon data, which are rarely available and uncertain to determine, especially in forest soils.

Ultimately, the Wessolek pedotransfer function was chosen as it was developed for German sites and has been tested for forest soils. It furthermore allowed us better to compare the performance of ForClim and LWFBrook90, as it had also been adopted in Meusburger et al. (2022).

The Wessolek function estimates soil water content (θ) based on pressure head (ψ) using the van Genuchten model. The function takes parameters such as *alpha, npar, mpar, ths, thr,* and calculates wetness and water content accordingly. Then, field capacity (FC) and wilting point (WP) are calculated, and the available water content is retrieved for the given pressure heads (63 hPa for FC and 15’848 hPa for WP) for each layer *n* as:

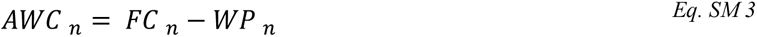

The volumetric available water content is then converted to gravimetric units (cm) by considering soil layer thickness and gravel content of each layer as:

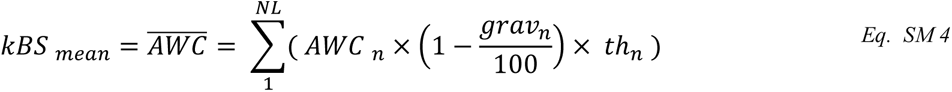

Where *NL* is the number of soil layers. The resulting soil hydraulic properties for each 25 x 25 m cell were averaged to obtain the mean values from which the mean bucket size was determined, as shown in Table D2; for each site, the minimum value of kBS was used to constrain kBS_min_ as:

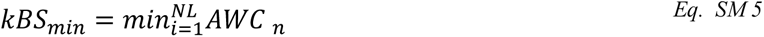

**Table D2:** Soil properties derived from the Wessolec pedotransfer function. For each site (Sid), the number of soil layers (*NL*) and their depth (upper and lower limits, cm) are given. For each soil horizon, texture (text, derived from clay,sand,silt (%)), and gravel content (grav, in %) are retrieved from the soil map of Baltensweiler et al. (2021) and averaged for each 25 m cell, and then used to calculate soil hydraulic properties, namely: *ths* (saturation water content fraction), *thr* (residual water content fraction), *alpha* (alpha parameter of the van Genuchten water retention function, *npar* (n parameter of the van Genuchten water retention function), *mpar* (m parameter of the van Genuchten water retention function), *ksat* (saturated hyraulic conductivity parameter of Mualem hydraulic conductivity function (mm/d)) and *tort* (tortuosity parameter of the Mualem hydraulic conductivity function).

| *Sid* | *NL* | *upper* | *lower* | *text* | *grav* | *ths* | *thr* | *alpha* | *npar* | *mpar* | *ksat* | *tort* |
| --- | --- | --- | --- | --- | --- | --- | --- | --- | --- | --- | --- | --- |
| GZ | 1 | 0 | -0.05 | Lu | 0.01 | 0.43 | 0.05 | 4.32 | 1.17 | 0.14 | 826.8 | -3.23 |
| GZ | 2 | -0.05 | -0.15 | Tu3 | 0.09 | 0.46 | 0 | 5.5 | 1.08 | 0.08 | 1,237.65 | 0 |
| GZ | 3 | -0.15 | -0.3 | Tu3 | 0.09 | 0.46 | 0 | 5.5 | 1.08 | 0.08 | 1,237.65 | 0 |
| GZ | 4 | -0.3 | -0.6 | Lt2 | 0.14 | 0.44 | 0.15 | 7.01 | 1.25 | 0.2 | 625.31 | -3.18 |
| GZ | 5 | -0.6 | -1 | Lu | 0.12 | 0.43 | 0.05 | 4.32 | 1.17 | 0.14 | 826.8 | -3.23 |
| VA | 1 | 0 | -0.05 | Tu3 | 0.06 | 0.46 | 0 | 5.5 | 1.08 | 0.08 | 1,237.65 | 0 |
| VA | 2 | -0.05 | -0.15 | Tu3 | 0.1 | 0.46 | 0 | 5.5 | 1.08 | 0.08 | 1,237.65 | 0 |
| VA | 3 | -0.15 | -0.3 | Tu3 | 0.13 | 0.46 | 0 | 5.5 | 1.08 | 0.08 | 1,237.65 | 0 |
| VA | 4 | -0.3 | -0.6 | Tu3 | 0.22 | 0.46 | 0 | 5.5 | 1.08 | 0.08 | 1,237.65 | 0 |
| VA | 5 | -0.6 | -1 | Lt2 | 0.27 | 0.44 | 0.15 | 7.01 | 1.25 | 0.2 | 625.31 | -3.18 |
| TB | 1 | 0 | -0.05 | Lt2 | 0.03 | 0.44 | 0.15 | 7.01 | 1.25 | 0.2 | 625.31 | -3.18 |
| TB | 2 | -0.05 | -0.15 | Lt3 | 0.05 | 0.45 | 0.16 | 4.95 | 1.17 | 0.15 | 443.4 | -4.1 |
| TB | 3 | -0.15 | -0.3 | Lt3 | 0.09 | 0.45 | 0.16 | 4.95 | 1.17 | 0.15 | 443.4 | -4.1 |
| TB | 4 | -0.3 | -0.6 | Lt3 | 0.13 | 0.45 | 0.16 | 4.95 | 1.17 | 0.15 | 443.4 | -4.1 |
| TB | 5 | -0.6 | -1 | Lt2 | 0.17 | 0.44 | 0.15 | 7.01 | 1.25 | 0.2 | 625.31 | -3.18 |
| UH | 1 | 0 | -0.05 | Tu3 | 0.08 | 0.46 | 0 | 5.5 | 1.08 | 0.08 | 1,237.65 | 0 |
| UH | 2 | -0.05 | -0.15 | Lu | 0.15 | 0.43 | 0.05 | 4.32 | 1.17 | 0.14 | 826.8 | -3.23 |
| UH | 3 | -0.15 | -0.3 | Lt2 | 0.33 | 0.44 | 0.15 | 7.01 | 1.25 | 0.2 | 625.31 | -3.18 |
| UH | 4 | -0.3 | -0.6 | Lt2 | 0.42 | 0.44 | 0.15 | 7.01 | 1.25 | 0.2 | 625.31 | -3.18 |
| UH | 5 | -0.6 | -1 | Lt2 | 0.4 | 0.44 | 0.15 | 7.01 | 1.25 | 0.2 | 625.31 | -3.18 |
| BB | 1 | 0 | -0.05 | Lt2 | 0.07 | 0.44 | 0.15 | 7.01 | 1.25 | 0.2 | 625.31 | -3.18 |
| BB | 2 | -0.05 | -0.15 | Lt2 | 0.17 | 0.44 | 0.15 | 7.01 | 1.25 | 0.2 | 625.31 | -3.18 |
| BB | 3 | -0.15 | -0.3 | Lt2 | 0.28 | 0.44 | 0.15 | 7.01 | 1.25 | 0.2 | 625.31 | -3.18 |
| BB | 4 | -0.3 | -0.6 | Lt2 | 0.38 | 0.44 | 0.15 | 7.01 | 1.25 | 0.2 | 625.31 | -3.18 |
| BB | 5 | -0.6 | -1 | Ls3 | 0.4 | 0.41 | 0.07 | 6.83 | 1.21 | 0.17 | 982 | -3.23 |
| RB | 1 | 0 | -0.05 | Lu | 0.01 | 0.43 | 0.05 | 4.32 | 1.17 | 0.14 | 826.8 | -3.23 |
| RB | 2 | -0.05 | -0.15 | Lu | 0.06 | 0.43 | 0.05 | 4.32 | 1.17 | 0.14 | 826.8 | -3.23 |
| RB | 3 | -0.15 | -0.3 | Lu | 0.07 | 0.43 | 0.05 | 4.32 | 1.17 | 0.14 | 826.8 | -3.23 |
| RB | 4 | -0.3 | -0.6 | Lu | 0.13 | 0.43 | 0.05 | 4.32 | 1.17 | 0.14 | 826.8 | -3.23 |
| RB | 5 | -0.6 | -1 | Lu | 0.17 | 0.43 | 0.05 | 4.32 | 1.17 | 0.14 | 826.8 | -3.23 |

### D3. Modeling of Predisposing and Inciting factors (after Manion, 1981)

#### D3.1. Proxy for the memory of carbon pools

Trees are capable of resisting and recovering from a drought event (Körner, 2019). To mimic this behavior in the absence of a full-tree carbon balance in ForClim, we followed the protocol developed by Zamora-Pereira et al. (2021) to constrain the diameter increment in subsequent years by means of a proxy. The constraints are based on yearly ratios of measured tree-ring width data for *Abies alba, Picea abies* and *Fagus sylvatica* (Bottero et al. 2019 and Cailleret et al. 2017). In this protocol, the smallest and largest ratios of inter-annual changes of tree-ring width are identified, respectively (Table D3.1, bold values). For the sake of consistency and parsimony, we selected the same two approximate values for all species, namely *0.2* and *5.0*.

In the current implementation, the yearly diameter increment of each cohort is corrected based on the values on the previous year’s growth. When the ratio of the current year is lower or higher than the empirical values, growth will be constrained accordingly. No lag effects longer than one year are considered.

**Table D3.1:** Ratios of yearly tree-ring widths according to Maxime Cailleret and Zamora-Pereira et al. (2021).

| Species | Quantile | Ratio* | Ratio** | Ratio*** |
| --- | --- | --- | --- | --- |
| Silver fir | 0.001 | 0.1420 | 0.2456 | **0.2086** |
|  | 0.999 | 4.3799 | 3.5517 | 3.7822 |
| Norway spruce | 0.001 | 0.3266 | 0.1632 | 0.1737 |
|  | 0.999 | 2.8598 | 5.7124 | 5.3877 |
| Beech | 0.001 | 0.1124 | 0.1426 | 0.1401 |
|  | 0.999 | 16.1329 | 6.0804 | **6.9725** |

\* Bottero et al. 2019, \*\* Cailleret et al. 2017, \*\*\* both databases (used for constraints)

#### D3.2. Seasonal soil moisture and REW critical in spring and autumn

The empirical threshold of “critical Relative Extractable Water content” (REW) suggested by Granier et al., 1999 has been used as reference for more than 20 years with a value ∼0.4. However, several studies have reported that REW varies strongly and seems to depend not only on soil properties but also on species-specific responses to drought via stomatal conductance, canopy conductance or, more generally, by transpiration strategies (Lindroth et al., 2018; Niu et al., 2023; Ruehr et al., 2012; Vilhar, 2016). In

Table D3.2.2.1 we summarize the range of variability of REW_c_ and report the temporal extent, species, biomes, and countries in which these values were assessed. These values were crucial to identify a range of plausible variability in the parameter space for the spring and fall period. Based on the literature review, we decided to set the range of variability for spring to [1.0 to 0.9] as in Chuste et al. (2019; cf. their Figure 2-A for control plots C in DoY 100-150), while for the fall period we identified the range [0.7 to 0.5] as in Vilhar et al. (2016). To assess the importance of choosing an exact value as well as to test the plausibility of these parameter ranges, we assessed the impact of the variation of REW across the beech sites in spring and fall, as shown in Figure D3.2.

**Table D3.2:** Critical REW (REW_c_) values according to a literature review. The temporal extent, main investigated species, biome and country are reported.

| $REW_c$ | Temporal extent | Species | Biome | Country | Reference |
| --- | --- | --- | --- | --- | --- |
| 0.7-0.4 | growing season (May-October) | *Fagus sylvatica* | Temperate | Slovenia | Vilhar U. Comparison of drought stress indices in beech forests: a modelling study. (2016) iForest 9: 635-642. doi: 10.3832/for1630-008 |
| 0.4-0.5 | growing season (2014-2015) | *Fagus sylvatica* | Temperate | Lorraine region (France) | Chuste, P. A., Massonnet, C., Gérant, D., Zeller, B., Levillain, J., Hossann, C., Angeli, N., Wortemann, R., Bréda, N., & Maillard, P. (2019). Short-term nitrogen dynamics are impacted by defoliation and drought in *Fagus sylvatica* L. Branches. *Tree Physiology*, 39(5), 792–804. https://doi.org/10.1093/treephys/tpz002 |
| $< 0.1$ | 15 days during growing season | *Quercus* spp. | Temperate | NW China | Niu, Xiaodong, and Shirong Liu. Environmental and stomatal control on evapotranspiration in a natural oak forest." *Ecohydrology* 15.4 (2022): e2423 |
| 0.4-0.5 | Late summer (August-September) | *Ponderosa pine* | Temperate | Central Oregon (USA) | Ruehr, N. K., Martin, J. G., & Law, B. E. (2012). Effects of water availability on carbon and water exchange in a young ponderosa pine forest: Above- and below-ground responses. *Agricultural and Forest Meteorology*, 164, 136–148. https://doi.org/https://doi.org/10.1016/j.agrformet.2012.05.015 |
| $< 0.4$ | annual- growing period | *Quercus aliena* | Warm Temperate | Central China | Niu, X., Chen, Z., Pang, Y., Liu, X., & Liu, S. (2023). Soil moisture shapes the environmental control mechanism on canopy conductance in a natural oak forest. *Science of the Total Environment*, 857. https://doi.org/10.1016/j.scitotenv.2022.159363 |
| $< 0.2$ or $< 0.05-0.5$ | (global sensitivity analysis) | *Pinus sylvestris* | Boreal | Hyytiälä (Finland) | Launiainen, S., Guan, M., Salmivaara, A., and Kieloaho, A.-J.: Modeling boreal forest evapotranspiration and water balance at stand and catchment scales: a spatial approach, *Hydrol. Earth Syst. Sci.*, 23, 3457–3480, https://doi.org/10.5194/hess-23-3457-2019, 2019 |
| 0.04 and 0.37 (Pinus), 0.18 and 0.39 (P. abies) | Late summer (June to September) | *Pinus sylvestris*, *Picea abies* | Boreal | Sweden | Lagergren, F., & Lindroth, A. (2002). Transpiration response to soil moisture in pine and spruce trees in Sweden. In *Agricultural and Forest Meteorology* (Vol. 112). |
| $< 0.4$ | Annual | *Fagus sylvatica*, *Picea abies*, *Pinus sylvestris* | Temperate/ Boreal | Hesse (F), Hainich (D), Sorø (DK), Tharandt (D), Hyytiälä (FI), Loobos (NL) | Bréda, N., Huc, R., Granier, A., & Dreyer, E. (2006). Temperate forest trees and stands under severe drought: a review of ecophysiological responses, adaptation processes and long-term consequences. *Annals of Forest Science*, 63(6), 625-644. |
| 0.33-0.45 | Growing period | *Pinus taeda*, *Liriodendron tulipifera*, *Liquidambar styraciflua*, *Carya* spp., *Quercus* spp. | Temperate | North Carolina (USA) | Oishi, A. C., Palmroth, S., Butnor, J. R., Johnsen, K. H., & Oren, R. (2013). Spatial and temporal variability of soil CO<sub>2</sub> efflux in three proximate temperate forest ecosystems. *Agricultural and Forest Meteorology*, 171–172, 256–269. https://doi.org/10.1016/j.agrformet.2012.12.007 |

**Figure D3.2:**
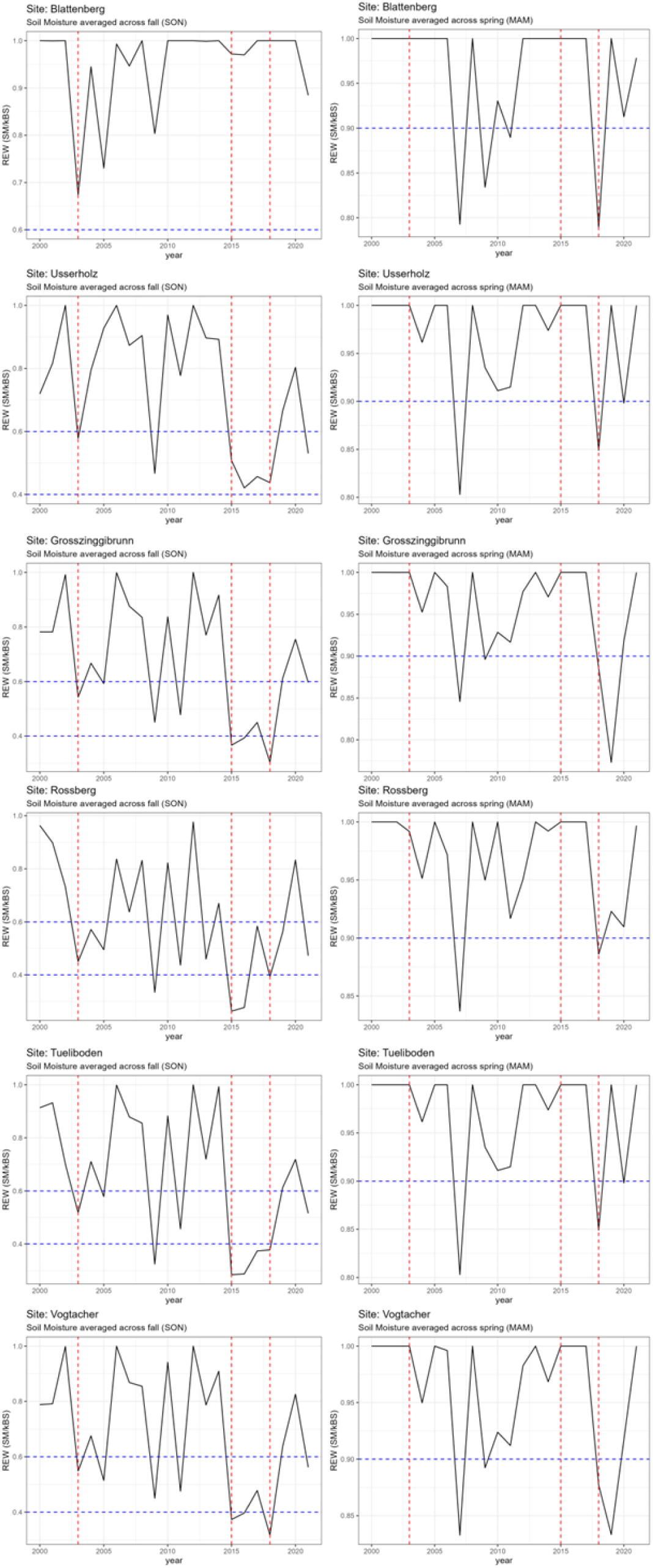
Relative extractable water content variation across the six beech-dominated sites in the fall (SON) and spring (MAM) months.

#### D3.3. Sensitivity analysis for new model parameters

Persistent uncertainty regarding parameter selection is a recurring challenge in ecological modeling (Huber et al., 2020). We confronted this when introducing four new parameters (i.e., *kREW_fall_, kREW_spring_, kDrTh, kEg*) that required rigorous constraints.

From the literature review, we identified the parameter range for *kREW* as presented in section D.3.2, hence the selected ranges for fall [0.7-0.5] and spring [1.0-0.9] were chosen as upper and lower boundary conditions. For the parameter *kDrTh*, we identified a threshold range spanning from 0.1 (lower boundary) to 0.3 (upper boundary) corresponding to a very low to very high drought sensitivity class associated with the drought index (*gDr*, Eq.9 in the main text), which usually varies between 0 (no drought) to 0.2 (medium-strong drought) during the growing season (Figure D3.3.1). For *kEg*, we assumed a variation between 1 (i.e., any deviation from the optimum E/D would count) or a slight threshold such as 0.9. This condition implies that the ratio of transpiration to demand should be sufficiently high during the growing season.

The scarcity of empirical data posed a challenge in this regard. To confine this uncertainty and to choose a plausible combination of parameter values within determined ranges, we adopted a two-fold approach aimed at exploring the parameter space effectively (cf. Table D3.3.1).

##### D3.3.1 Short-term patterns

We focused on identifying a target pattern which operates in the short term, specifically the basal area losses triggered by the droughts of 2018-2020 at the six beech-dominated sites. The analyses of the short-term pattern allowed us to confirm the pre-conceived notion about the optimal parameter ranges. Yet, to ensure the model’s generalizability across diverse environmental conditions and future case studies, we extended our analysis to include sites of the European gradient, particularly those characterized by high aridity. Therefore, we selected a long-term pattern focusing on basal area development over time and the frequency of mortality events reducing it. This strategy, inspired by the pattern-oriented modeling framework (Grimm et al., 2005), facilitated a comprehensive exploration of parameter values that accounted for both short-term disturbances and long-term ecological dynamics, enhancing the overall robustness and applicability of our model.

We developed a scoring system in the range [0,1] to eliminate the scenarios which were failing to reproduce the pattern in the short term. A parameter combination leading to basal area losses in the exact years of observation (2018-2020) would score 1. Furthermore, the scores varied based on how widespread the mortality events were across multiple sites: *i)* if mortality events occurred in more than two out of six sites in the exact year of observation the score was 0.75; *ii)* if mortality events occurred in two out of six sites the score dropped to 0.5; *iii)* if mortality events occurred in one out of six sites the score further dropped to 0.25. Only the scenarios with a score ≥ 0.75 were selected for testing on long-term simulations across the European gradient of sites (cf. main text). The scenarios that reproduced a scoring greater than 0 are shown in Table D3.3.2.

**Table D3.3.1:** Pattern-oriented modelling framework to assess parameter uncertainty (after Huber, 2019)

| Pattern | Point in time | Level |
| --- | --- | --- |
| *Short-term* |  |  |
| Basal area losses | 2018-20<br>(drought-induced mortality event) | Total stand |
| *Long-term* |  |  |
| Frequency in basal area losses | Dynamic equilibrium<br>(years -1500) | Total stand |

**Table D3.3.2:** Scenario definition as combination of the parameters REW in autumn and spring and the two drought tolerance thresholds *kDrTh* (Eq. 8) and *g* (Eq. 9) in the main text. Highlighted in red are the parameter combinations with a score of 0.75. In the red shaded rows, we show the best parameter set, which was chosen in this study.

| Scenario | $kREW_{fl}$ | $kREW_{sp}$ | $kDrTh$ | $kEg$ |
| --- | --- | --- | --- | --- |
| 1 | 0.5 | 1 | 0.1 | 1 |
| 2 | 0.6 | 1 | 0.1 | 1 |
| 3 | 0.7 | 1 | 0.1 | 1 |
| 4 | 0.5 | 0.9 | 0.1 | 1 |
| 5 | 0.6 | 0.9 | 0.1 | 1 |
| 6 | 0.7 | 0.9 | 0.1 | 1 |
| 7 | 0.5 | 1 | 0.2 | 1 |
| 8 | 0.6 | 1 | 0.2 | 1 |
| 9 | 0.7 | 1 | 0.2 | 1 |
| 10 | 0.5 | 0.9 | 0.2 | 1 |
| 11 | 0.6 | 0.9 | 0.2 | 1 |
| 12 | 0.7 | 0.9 | 0.2 | 1 |
| 13 | 0.5 | 1 | 0.3 | 1 |
| 14 | 0.6 | 1 | 0.3 | 1 |
| 15 | 0.7 | 1 | 0.3 | 1 |
| 16 | 0.5 | 0.9 | 0.3 | 1 |
| 17 | 0.6 | 0.9 | 0.3 | 1 |
| 18 | 0.7 | 0.9 | 0.3 | 1 |
| 19 | 0.5 | 1 | 0.1 | 0.9 |
| 20 | 0.6 | 1 | 0.1 | 0.9 |
| 21 | 0.7 | 1 | 0.1 | 0.9 |
| 22 | 0.5 | 0.9 | 0.1 | 0.9 |
| 23 | 0.6 | 0.9 | 0.1 | 0.9 |
| 24 | 0.7 | 0.9 | 0.1 | 0.9 |
| 25 | 0.5 | 1 | 0.2 | 0.9 |
| 26 | 0.6 | 1 | 0.2 | 0.9 |
| 27 | 0.7 | 1 | 0.2 | 0.9 |
| 28 | 0.5 | 0.9 | 0.2 | 0.9 |
| 29 | 0.6 | 0.9 | 0.2 | 0.9 |
| 30 | 0.7 | 0.9 | 0.2 | 0.9 |
| 31 | 0.5 | 1 | 0.3 | 0.9 |
| 32 | 0.6 | 1 | 0.3 | 0.9 |
| 33 | 0.7 | 1 | 0.3 | 0.9 |
| 34 | 0.5 | 0.9 | 0.3 | 0.9 |
| 35 | 0.6 | 0.9 | 0.3 | 0.9 |
| 36 | 0.7 | 0.9 | 0.3 | 0.9 |

We selected a total of 36 scenarios as result of the combination of the 4 parameter values (Table D3.3.2), which were evaluated for the six beech sites. All scenarios led to different degrees of basal area losses (cf. Figure D3.3.2), most commonly including the years 2003-2004, 2008-2009 and the years 2016-2017, except for scenario 28 which was identified as best matching the observations (Figure D3.3.3) and those scenarios with a score greater than 0.5 which were then selected for further testing (red values in Table D3.3.2, cf. also Figure D3.3.2).

**Figure D3.3.1:**
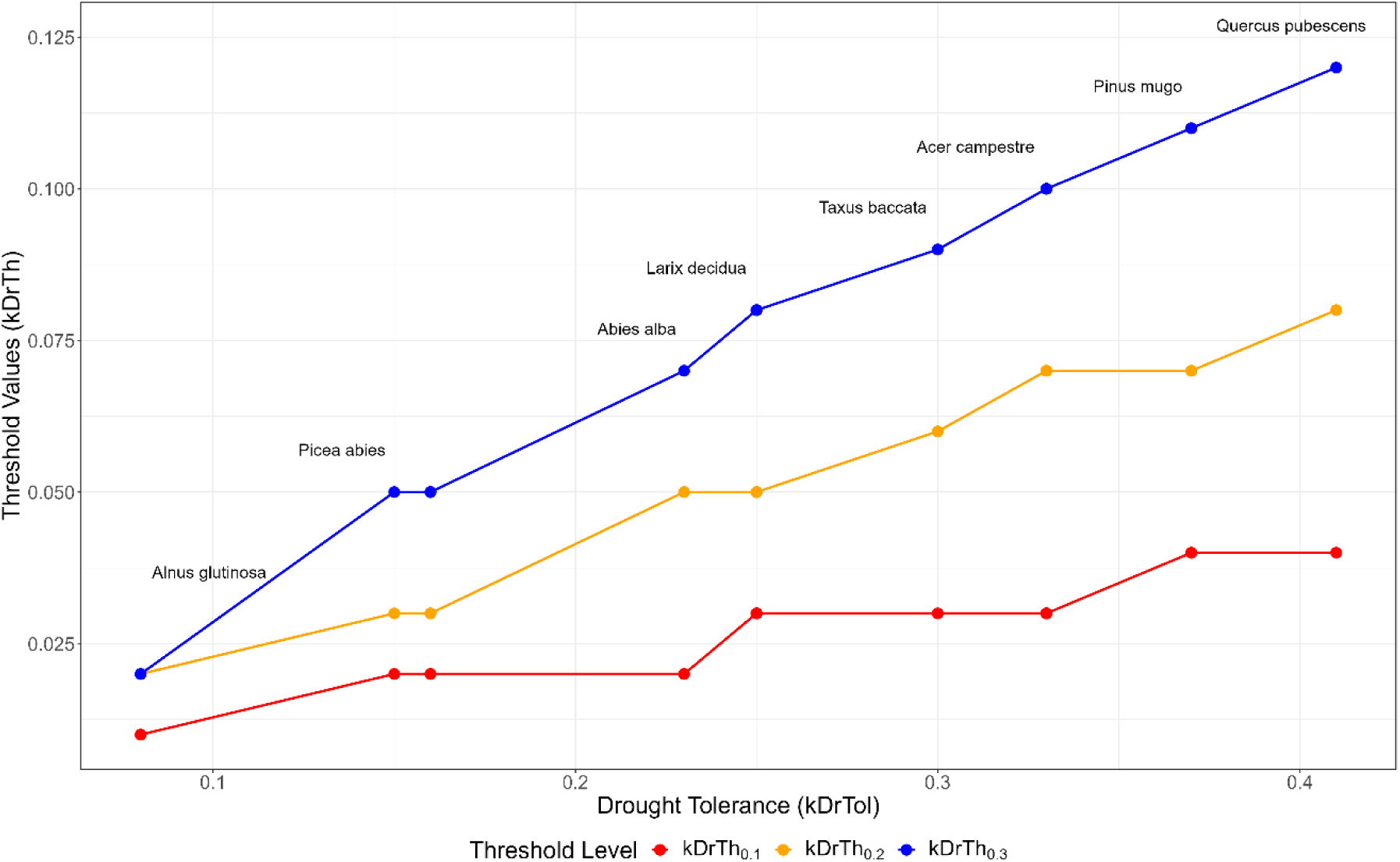
Drought sensitivity threshold classes from very low (0.1) in red to very high (0.3) in blue, in relation to the drought tolerance classes (*kDrTol*) of the tree species parameterized in ForClim. Each class corresponds to different threshold values (*kDrTh*) per species group. As examples, 8 representative species of the 32 species parameterized in ForClim are labelled and shown in the plot.

**Figure D3.3.2:**
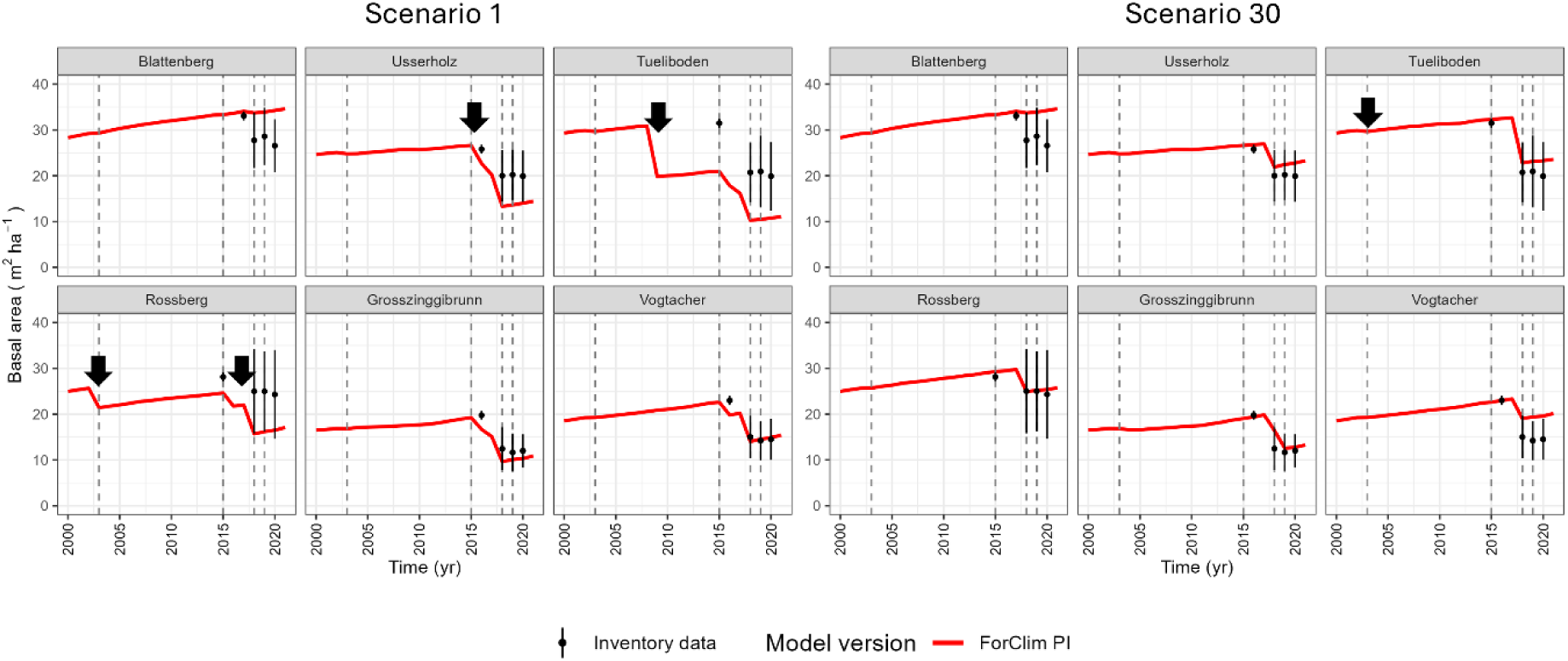
Basal area development in scenarios 1 and 30 resulting from the combination of the four parameters *kREW_fall_, kREW_spring_, kEg* and *kDrTh* (cf. Table D3.3.2). The black arrows indicate the mortality years considered as artifacts.

**Figure D3.3.3:**
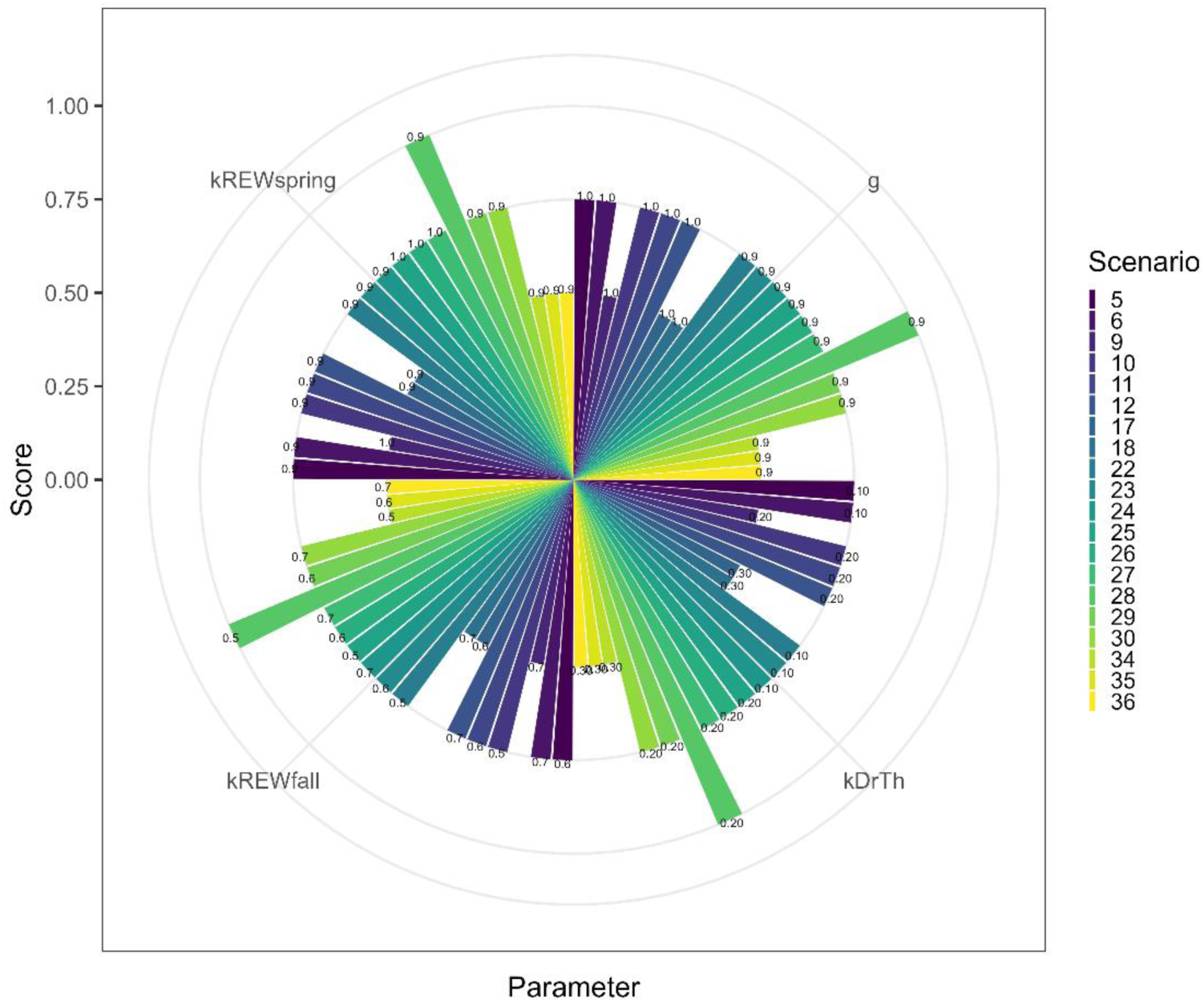
Combination of the four parameters *kREW_fall_, kREW_spring_, kEg* (shown as g in the plot) and *kDrTh* according to the parameter range selected in Table ***D3.3.2***. Each colored bar represents a unique parameter combination (scenario) with its score. The scenarios which scored a value> than 0.25 are shown in the plot.

##### D3.3.2 Long-term patterns

We assessed the long-term patterns by evaluating the mortality frequency on selected default sites, namely Davos, Adelboden and Potsdam, resulting in 33 simulation scenarios. Simulations started in the year 0 and ended in the year 1500, using model variant 22 (Huber et al. 2020), while the simulation setup was the same as in the main experiment. To systematically assess the differences within each scenario (S) - site (N) combination, we developed a cost function (𝐶_𝑠_) that calculates the sum of squared differences between the simulated basal area (Eq. SM6) and its lagged value (time points, (𝑇), cf. Eq. SM7). To facilitate the choice of the optimum parameter set, the scenario with the lowest cost (𝑠^∗^, Eq. SM8.), indicating the most favorable outcome in terms of the evaluated variable, was selected.

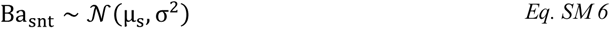

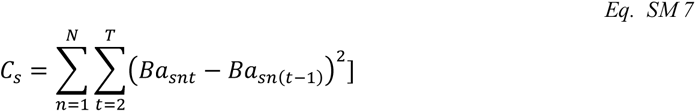

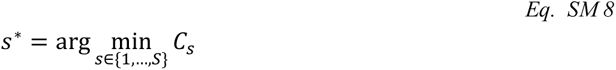

This calculation helped us quantify the deviation or change in the variable under consideration, aiding in the evaluation of scenario performance by assigning higher cost to scenarios that result in larger changes in basal area over time (Figure D3.3.4). While no differences were observed at the site Adelboden, scenario 6 yielded the lowest cost at Davos (cold part of the gradient), while scenarios 6, 12, 24 and 30 resulted in low costs at Potsdam (dry part of the gradient). Therefore, we concluded that the parameter combination of scenario 6 can be used as a second-best alternative to the chosen parameter selection for scenario 28.

**Figure D3.3.4:**
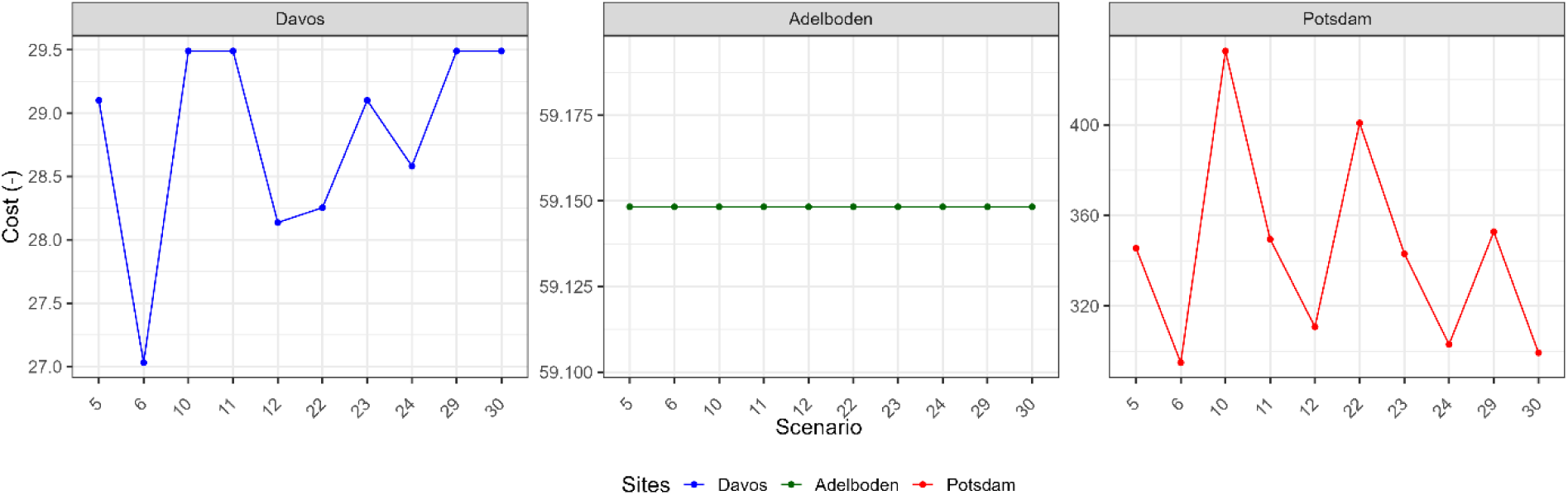
Cost function (see Eq. 14) associated with each scenario for three sites along the European climatic gradient.

#### D4: kDD re-estimation

In the new formulation of the growth equation (cf. Eq. 6 of the main text), we enhanced the growth response to climatic extremes by applying Liebig’s law of the minimum between temperature and soil water dynamics, rather than multiplying the two factors and keeping them within the root function. Thus, the growth reduction associated with temperature (DDGF) and moisture (SMGF) became substantially larger in ForClim v4.1. This has considerable impacts on forest dynamics, particularly in cold and dry conditions. We therefore applied a formal equivalence between the two DDGF formulations, and defined the system:

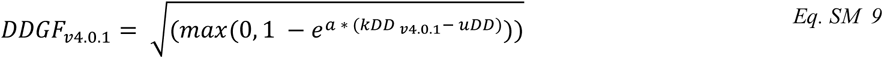

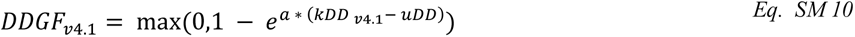

where kDD stands for the species-specific minimal annual degree-day sum. uDD is the monthly sum of degree-days (°C·days), and *a* is the slope parameter. Values for kDD _v4.1_ were approximated iteratively: the Newton-Raphson iteration algorithm starts with an initial guess by setting kDD _v4.1_ to the value of kDD _v4.0.1_ and the function iteratively updates the guess based on the differences between DDGF _v4.0.1_ and DDGF_v4.1_, converging towards a value of kDD _v4.1_ that satisfies a specified tolerance value set to 0.1.

The kDD and uDD values used for testing were retrieved from simulations along the European site gradient. We fitted a linear model to predict the approximate values of kDD _v4.1_ based on kDD _v4.0.1_. The highest performance was obtained with the following model (Table D4.1):

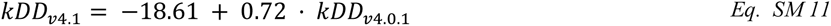

**Table D4.1:**
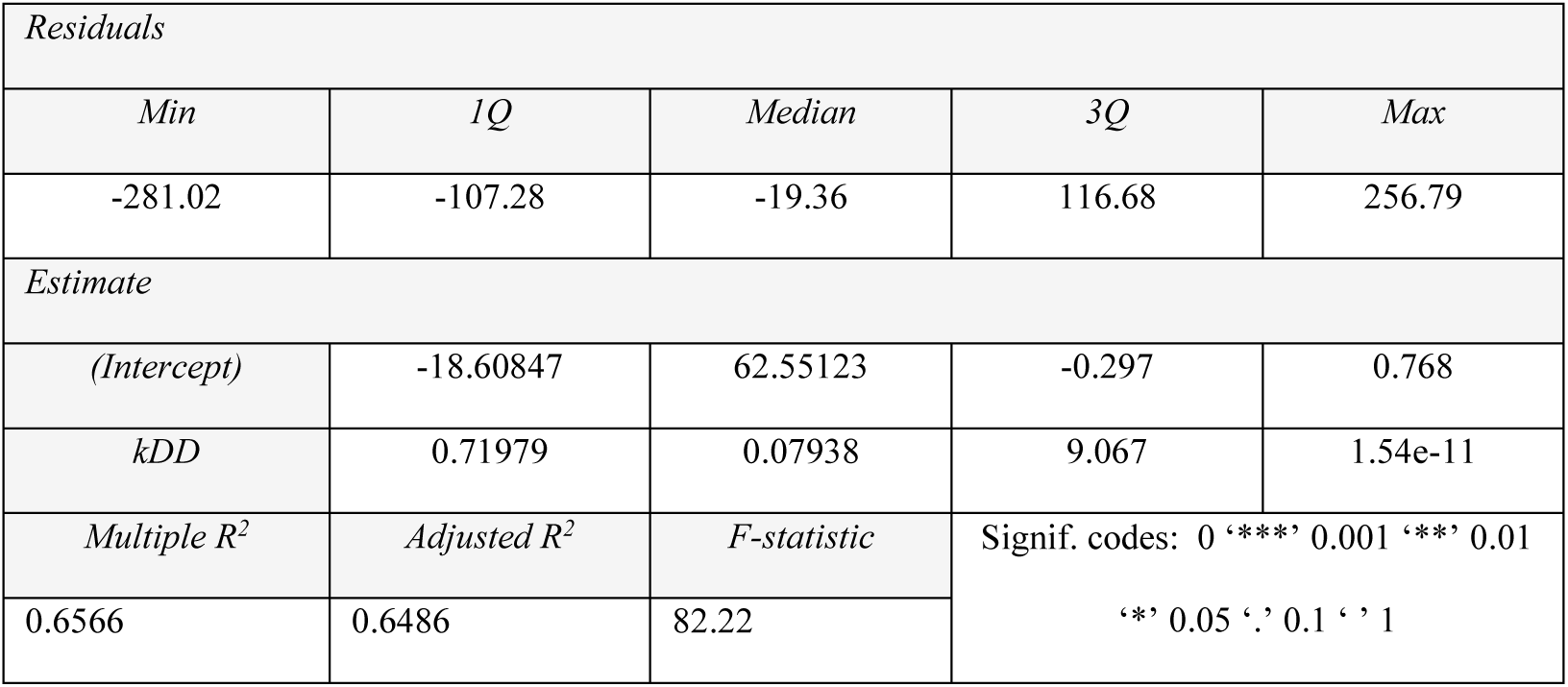
Summary statistics of linear model for kDD*_v4.1_* estimation

The newly estimated parameter values can be found in Table D4.2.

**Table D4.2:** Current (v4.0.1) and newly estimated (v4.1) kDD parameter values for the European tree species in ForClim.

| *kName* | *kDD* v4.0.1 | *kDD* v4.1 |
| --- | --- | --- |
| *Alnus viridis* | 350 | 233 |
| *Larix decidua* | 350 | 233 |
| *Pinus cembra* | 350 | 233 |
| *Picea abies* | 350 | 233 |
| *Betula pendula* | 425 | 287 |
| *Populus tremula* | 425 | 287 |
| *Pinus montana* | 500 | 341 |
| *Sorbus aucuparia* | 500 | 341 |
| *Pinus sylvestris* | 500 | 341 |
| *Alnus incana* | 500 | 341 |
| *Pseudotsuga menziesii* | 633 | 437 |
| *Abies alba* | 650 | 449 |
| *Acer pseudoplatanus* | 650 | 449 |
| *Salix alba* | 650 | 449 |
| *Populus nigra* | 750 | 521 |
| *Alnus glutinosa* | 750 | 521 |
| *Sorbus aria* | 750 | 521 |
| *Tilia cordata* | 750 | 521 |
| *Fagus sylvatica* | 850 | 593 |
| *Corylus avellana* | 850 | 593 |
| *Fraxinus excelsior* | 850 | 593 |
| *Acer platanoides* | 850 | 593 |
| *Ulmus glabra* | 850 | 593 |
| *Tilia platyphyllos* | 850 | 593 |
| *Quercus petraea* | 950 | 665 |
| *Carpinus betulus* | 950 | 665 |
| *Acer campestre* | 950 | 665 |
| *Taxus baccata* | 1050 | 737 |
| *Quercus robur* | 1050 | 737 |
| *Castanea sativa* | 1050 | 737 |
| *Quercus pubescens* | 1150 | 809 |

### D5: kRedMax and kH_min_

#### D5.1. Dynamic calculation of maximum tree height and climatological averaging of drought and degree-days

When utilizing weather time series in the ForClim simulations, it is logical to dynamically calculate the maximum height that trees (*gHMax*) can reach, rather than to determine it prior to the simulation based on the assumption of a constant “current” climate, as done in previous versions of ForClim (cf. Rasche et al. 2012). This change in the approach was necessary because weather time series are inherently non-stationary. By recalculating *gHMax* annually based on a moving window approach, we can reflect the influence of changing climatic conditions on tree growth (“site index”), ensuring that the model is responsive to temporal fluctuations in temperature, precipitation, and other critical environmental factors. This dynamic adjustment is crucial for capturing realistic growth patterns and ecological responses of trees in a varying climate, including drought-induced mortality.

To this end, averages of the drought index and the degree-day sum are calculated over a 30-year period using a moving average, thereby treating these averages as a climatology underlying the calculating of “site index”.

#### D5.2. Minimum height at the species distribution limit

Rasche et al. (2012) used the 𝑘𝑅𝑒𝑑𝑀𝑎𝑥 parameter to quantify the maximum reduction in tree height based on suboptimal temperature and drought conditions, using data from the Swiss National Forest Inventory (NFI) and Growth-and-Yield plots. This parameter was designed to reflect the maximum height reduction due to environmental stressors, particularly drought and degree-days, but its estimation is not suitable for simulations outside productive forests, thus potentially misrepresenting ecological dynamics at the distributional limits of the species (Körner et al. 1998). To address this limitation, we assumed that the parameter 𝑘𝐻_min_ should reflect the minimum height that trees (which survive for a long time) can achieve at their upper and lower distributional limits, in the most extreme case at treeline.

Yet, the definition of a tree height threshold to define forests varies from 5-3 m (Jeník & Lokvenc 1962, Körner 2012) in Northern Central Europe to 2 m (Holtmeier et al. 2009) in Central Southern Europe (Cudlín et al. 2017). Moreover, recent studies (Gelabert et al. 2024) indicate that up to around 1500 m a.s.l., trees reach a fairly constant maximum height in the European Alps, decreasing at a rate of -1.27 m per 100 m of elevation. Despite the limitations of the study, we approximated the minimum height of trees to be 15 m by taking the mean between the estimated maximum tree height at the highest site of the European gradient (2300 m a.s.l., site of GrandeDixence), which amounts to 27 m and the maximum tree height reported by Körner for the Swiss Alps of 3 m. We therefore set 𝑘𝐻_min_ to 15 m and recalculated 𝑘𝑅𝑒𝑑𝑀𝑎𝑥 according to Eq. 19. Furthermore, the model updates the maximum tree height (𝑔𝐻𝑀𝑎𝑥) annually to incorporate the most recent trends and environmental conditions (Eq. SM 19-24). This adjustment ensures that calculated 𝑔𝐻𝑀𝑎𝑥 reflects the latest data trends and environmental conditions, thus enhancing the reliability of our simulations.

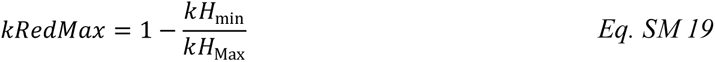

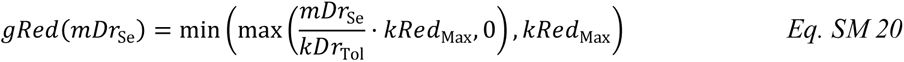

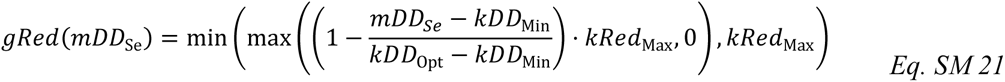

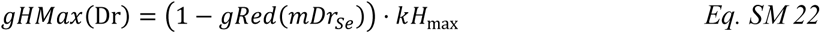

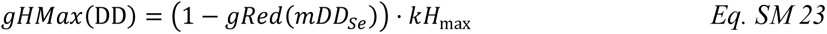

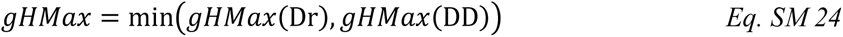

### D6: Site parameters for the six beech sites as well as the LWF sites Lägeren and Visp

**Table D6:** Summary table of site characteristics at the six drought-prone beech sites and at the ICP-Level II plots.

| Site | $kPatchSize$<br>($m^2$) | $kBS_{mean}$<br>($cm$) | $kBS_{min}$<br>($cm$) | $kAvN$<br>($kg\ ha^{-1}\ y^{-1}$) | $SN^1$<br>($p\ ha^{-1}$) |
| --- | --- | --- | --- | --- | --- |
| Blattenberg | 550 | 8.6 | 8.2 | 100 | 180 |
| Grosszinggibrunn | 550 | 15.4 | 14.4 | 180 | 245 |
| Rossberg | 550 | 15.5 | 13.3 | 180 | 187 |
| Tüeliboden | 550 | 14 | 12.6 | 180 | 170 |
| Usserholz | 550 | 11.2 | 10.2 | 180 | 165 |
| Vogtacher | 550 | 13.7 | 12.6 | 180 | 201 |
| Visp | 750 | 7.0 | 9.0 | 100 | 232<sup>(i)</sup> |
| Lägeren | 550 | 8.5 | 7.3 | 180 | 503<sup>(ii)</sup> |

<sup>1</sup> Simulated Stem Number. Tree number with a DBH > 8 cm
<sup>(i)</sup> Observed stem number per hectare 226 p ha<sup>-1</sup> in 1996 (Dobberdin et al 2005)
<sup>(ii)</sup> Observed stem number per hectare was 503 in 2011 (SwissFluxnet, 2024)

### D7: Site parameters along the climatic gradient

**Table D7.1:** Summary table of average temperature (T, °C) and precipitation sum (P, cm) for the 12 European sites. Weather data were obtained from Bugmann & Solomon (2000).

| *Site* | *T* | *P* |
| --- | --- | --- |
| Grande-Dixence | 1.3 | 1017 |
| Bever (S) | 1.5 | 841 |
| Davos | 3 | 1007 |
| Adelboden | 5.5 | 1351 |
| Huttwil | 8.1 | 1287 |
| Bern | 8.4 | 1006 |
| Schaffhausen | 8.6 | 882 |
| Basel | 9.2 | 784 |
| Schwerin | 8.2 | 625 |
| Potsdam | 8.7 | 589 |
| Cottbus | 8.77 | 573 |
| Sion | 9.7 | 597 |

**Table D7.2:** kBS values across the European gradient (a: Swiss sites, b: German sites). Buffer from kBS_min,M_ from Meusbuerger et al, kBS_min,41_ as the value chosen for ForClim v4.1 and kBS for ForClim v4.0.1 as in Bugmann and Solomon (2000).

| **a) Swiss sites** |  |  |  |  |  |  |  |  |
| --- | --- | --- | --- | --- | --- | --- | --- | --- |
| site | $kBS_{min,M}$ | $kBS_{mean,M}$ | $kBS_{min,41}$ | $kBS_{mean,41}$ | kBS | $\Delta[kBS_{mean,M} - kBS_{mean,41}]$ | $\Delta[kBS - kBS_{mean,M}]$ | $\Delta[kBS - kBS_{mean,41}]$ |
| Grande Dixence | 9.5 | 10.0 | 7.0 | 8.0 | 10.0 | 2.0 | 0.0 | 2.0 |
| BeverS | 8.3 | 9.4 | 8.1 | 9.0 | 10.0 | 0.4 | 0.6 | 1.0 |
| Davos | 8.2 | 9.3 | 5.0 | 7.0 | 10.0 | 2.3 | 0.7 | 3.0 |
| Adelboden | 9.1 | 10.6 | 9.0 | 10.0 | 15.0 | 0.6 | 4.4 | 5.0 |
| Huttwil | 13.5 | 15.2 | 12.0 | 13.0 | 20.0 | 2.2 | 4.8 | 7.0 |
| Bern | 13.5 | 14.9 | 9.0 | 14.0 | 20.0 | 0.9 | 5.1 | 6.0 |
| Schaffhausen | 9.5 | 11.6 | 9.0 | 13.0 | 15.0 | -0.5 | 3.4 | 2.0 |
| Basel | 12.4 | 13.9 | 11.0 | 14.0 | 15.0 | -0.1 | 1.1 | 1.0 |
| Sion | 10.0 | 11.4 | 9.0 | 10.0 | 15.0 | 1.4 | 3.6 | 5.0 |
| **b) German sites** |  |  |  |  |  |  |  |  |
| site | $kBS_{min,BGB}$ | $kBS_{mean,BGB}$ | $kBS_{min,41}$ | $kBS_{mean,41}$ | kBS | $\Delta[kBS_{mean,BGB} - kBS_{mean,41}]$ | $\Delta[kBS - kBS_{mean,BGB}]$ | $\Delta[kBS - kBS_{mean,41}]$ |
| Schwerin | 5.0 | 18.5 | 4.0 | 19.0 | 24.0 | -0.5 | 5.5 | 5.0 |
| Potsdam | 5.0 | 23.2 | 5.0 | 25.0 | 24.0 | -1.8 | 0.8 | -1.0 |
| Cottbus | 4.6 | 9.6 | 4.0 | 10.0 | 24.0 | -0.4 | 14.4 | 14.0 |

**Table D7.3:** ForClim v4.1 is based on ForClim v4.0.1 (Huber et al., 2020), which offered a total of 504 model variants. The table shows the setup of ForClim v4.1 and salient differences to ForClim v4.0.1 beyond those explained in the main manuscript. Acronyms: *A* stands for the species-specific light-dependent allocation into height vs diameter growth based on the scaled shade tolerance; *E* represents the establishment probability routine according to which the establishment probability (*kEstP)* is set to 4%. The mortality *M* is distinguished in background mortality (𝑀_0_) and stress-induced mortality. The latter is further differentiated in growth dependent (𝑀_SG_) and the predisposing-inciting stress dependent mortality (𝑀_S-PI_).

| ForClim<br>version | Model<br>variant | Weather<br>approach | Soil<br>bucket | Height<br>reduction | Establishment<br>routine | Allocation of<br>growth | Background<br>mortality | Stress-induced<br>mortality |
| --- | --- | --- | --- | --- | --- | --- | --- | --- |
| 4.0.1. | 22 | Generated | One | kRedMax | $E_6^*$ | $A_2$ | $M_0$ | $M_{S-G}$ |
| 4.1. | 22 | Time series | Distributed | Minimum height | $E_6^*$ | $A_2$ | $M_0$ | $M_{S-PI}$ |

**Figure D7.1:**
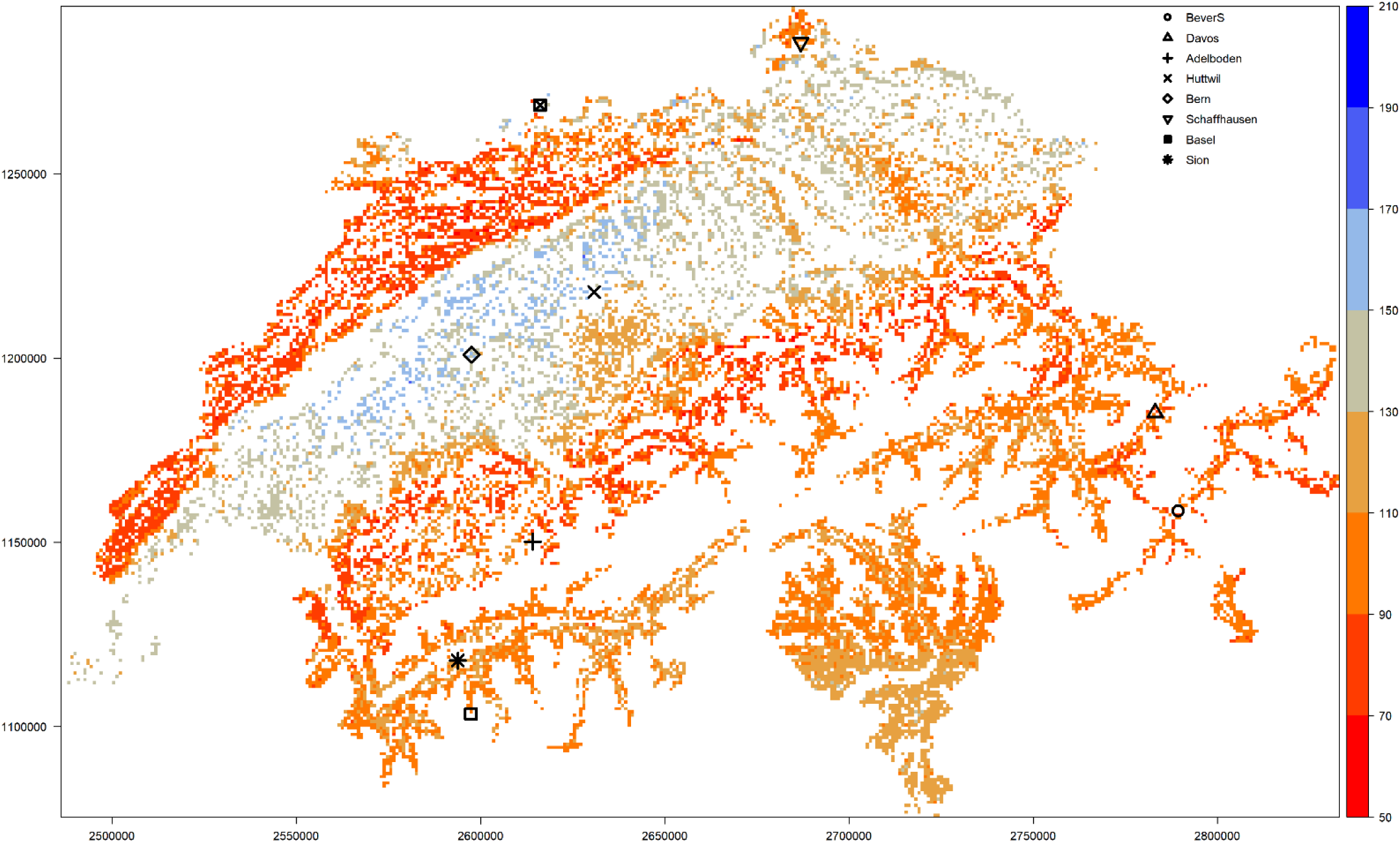
*Available Water Capacity (mm) maps by Meusburger et al. (2022).* Symbols indicate the nine Swiss sites of the European gradient.

**Figure D7.2:**
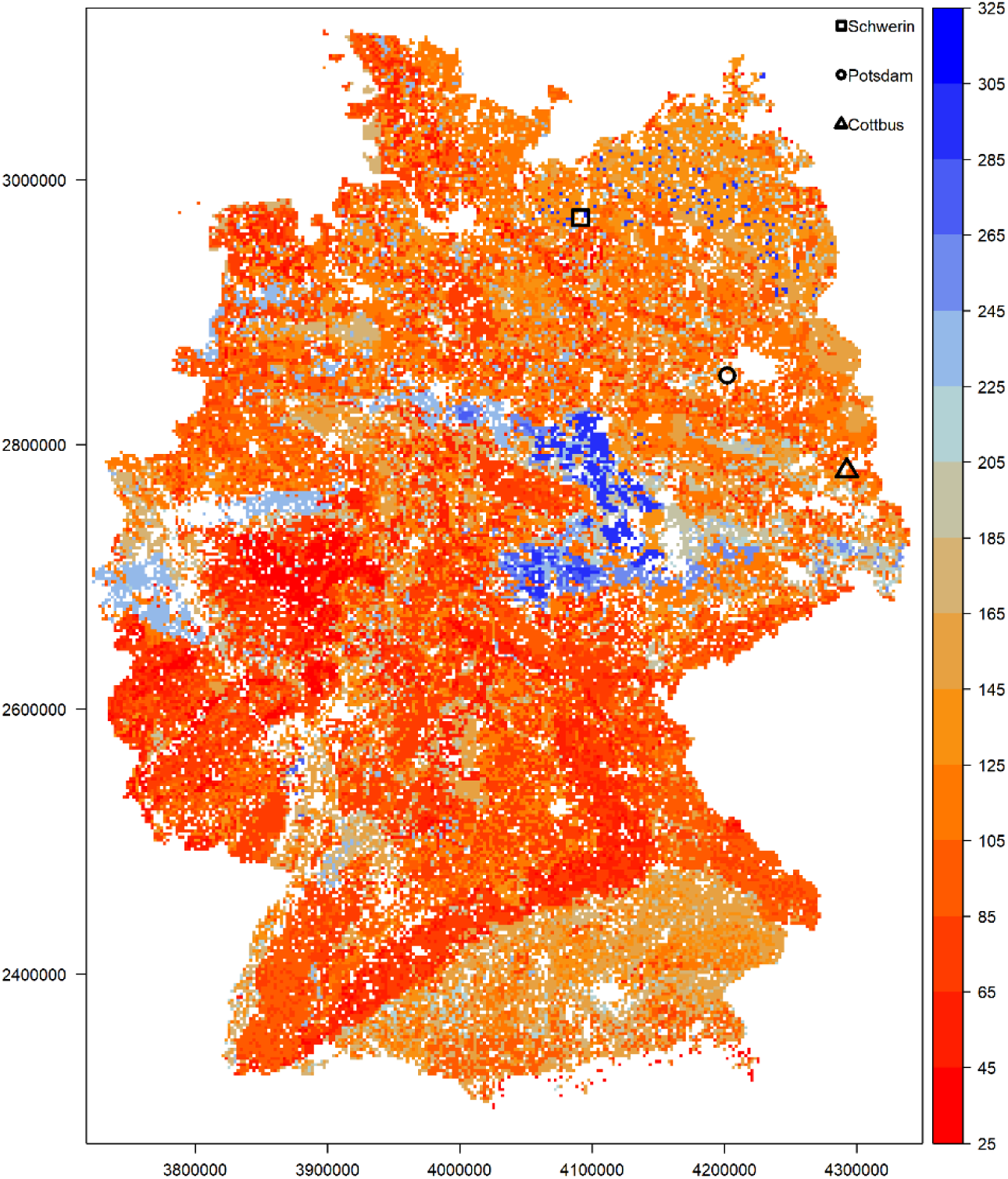
*Available Water Capacity (mm) maps produced by BGB.* Symbols indicate the three German sites of the European gradient.

**Figure D7.3:**
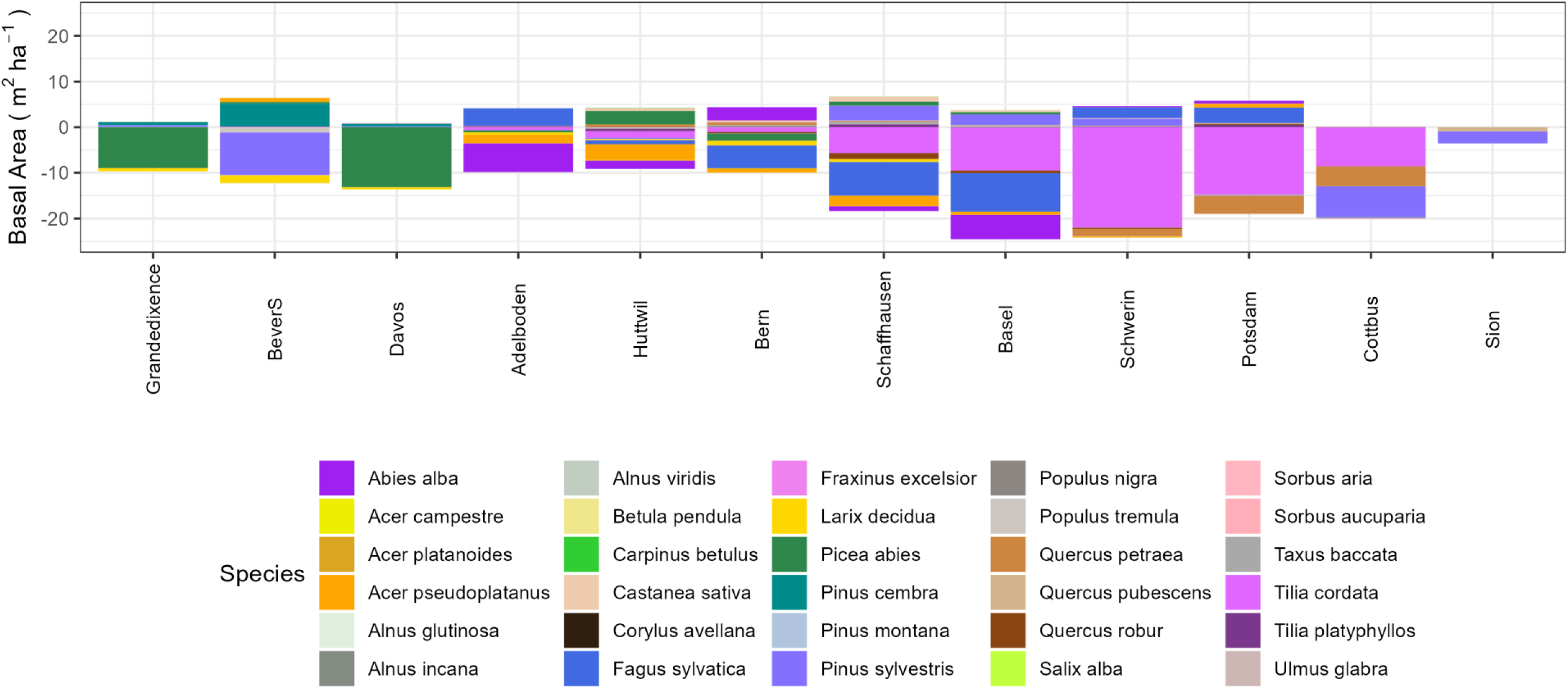
Basal area differences (m^2^ ha^-1^) between ForClim v4.0.1 variant 22 minus ForClim v4.1 variant 22 across the 12 default sites.

